# High risk glioblastoma cells revealed by machine learning and single cell signaling profiles

**DOI:** 10.1101/632208

**Authors:** Nalin Leelatian, Justine Sinnaeve, Akshitkumar M. Mistry, Sierra M. Barone, Kirsten E. Diggins, Allison R. Greenplate, Kyle D. Weaver, Reid C. Thompson, Lola B. Chambless, Bret C. Mobley, Rebecca A. Ihrie, Jonathan M. Irish

## Abstract

Recent developments in machine learning implemented dimensionality reduction and clustering tools to classify the cellular composition of patient-derived tissue in multi-dimensional, single cell studies. Current approaches, however, require prior knowledge of either categorical clinical outcomes or cell type identities. These algorithms are not well suited for application in tumor biology, where clinical outcomes can be continuous and censored and cell identities may be novel and plastic. Risk Assessment Population IDentification (RAPID) is an unsupervised, machine learning algorithm that identifies single cell phenotypes and assesses clinical risk stratification as a continuous variable. Single cell mass cytometry evaluated 34 different phospho-proteins, transcription factors, and cell identity proteins in tumor tissue resected from patients bearing *IDH* wild-type glioblastomas. RAPID identified and characterized multiple biologically distinct tumor cell subsets that independently and continuously stratified patient outcome. RAPID is broadly applicable for single cell studies where atypical cancer and immune cells may drive disease biology and treatment responses.

## Introduction

Malignant cells in human tumors are remarkably diverse in their functional cell identities and this intra-tumor cellular heterogeneity is closely linked to patient outcomes ^1, 2^. However, bench and computational tools that have driven our understanding of altered phospho-protein signaling networks in cancer have historically been under-used in solid tumor research due to a lack of technology and samples. In blood cancers, single cell profiles of signaling networks have revealed cancer cells present at diagnosis whose abundance is closely linked to subsequent clinical outcomes, including patient survival ^3–7^. This single cell snapshot proteomics approach focuses on a select set of key proteins that govern cell functional identity and can robustly measure targets, such as phospho-proteins, that are inaccessible to sequencing modalities. Suspension mass cytometry is a valuable platform for solid tumor analysis, as it is relatively low cost, well-powered to detect rare and novel cell types, and able to sensitively measure phosphorylated transcription factors and other mechanistic determinants of cancer cell identity ^8, 9^.

Quantitative analysis of single cell cytometry data has recently moved from an era of human-driven identification of cell types using guide markers (expert gating) and embraced machine learning tools that can automatically reveal and characterize novel and abnormal cells ^10–13^. In building an automated cytometry workflow, algorithm developers need to decide whether users will supervise the discovery of cell subsets using clinical knowledge. CITRUS is an automated cell subset discovery tool that uses prior knowledge of categorical labels, such as “disease” or “healthy”, to identify cell clusters associated with those labels ^14^ CellCNN is another supervised analysis tool that requires prospective assignment of samples to categories and uses convolutional neural networks to learn a filter that predicts whether new cells match one of the groups ^15^. Other cell subset discovery approaches do not supervise the analysis with knowledge of clinical outcomes but do use prior biological knowledge to identify cell subpopulations and then test whether differential outcomes are associated these cell subsets ^5, 6, 16^. In mass cytometry analysis, another common approach is to use tools for automated, unsupervised cell discovery and characterization, including SPADE ^17^, t-SNE ^18^, UMAP ^19^, FlowSOM ^20^, and Marker Enrichment Modeling (MEM ^21^). These tools help explore the structure of multidimensional data and reveal subpopulations that can be overlooked in expert manual analysis ^10, 12, 13, 22^. However, while it is possible to quickly review enriched features of the groups ^21^, it would also be powerful to test whether groups with similar phenotypes share an association with differential risk of death ^23^. RAPID, a fully unsupervised workflow presented here, implements t-SNE, FlowSOM, and MEM analysis of single cell mass cytometry data to reveal risk stratifying cell populations ^23^.

Mass cytometry has recently been developed for human solid tumors ^11, 16, 24, 25^, including glioblastoma, the most common primary malignant brain tumor in adults ^26^. The median survival of glioblastoma patients after diagnosis has remained approximately 12-15 months for over a decade ^27, 28^. These highly aggressive tumors are composed of both tumor and stromal cells, which harbor diverse genomic, transcriptomic, and proteomic expression profiles reflecting abnormal neural lineages ^24, 29, 30^. Previous studies in glioblastomas have either measured signaling states in bulk primary tumors ^31–33^ or characterized genomic and transcriptomic profiles in a modicum of single cells (<1000) ^29, 30, 34, 35^. These approaches, however, have not yet improved clinical practice or outcome for patients with glioblastoma. Although both inter- and intra-tumoral dysregulation of signaling in glioblastoma, particularly the disruption of receptor tyrosine kinase (RTK) homeostasis, is hypothesized to drive disease aggressiveness, very little is known about the activation states of signaling effector proteins in glioblastoma and how these signaling changes may be associated with cell subpopulations and patient clinical outcomes ^36, 37^. Novel, molecularly-driven criteria may give valuable insights into the biology of tumor progression and identify patients more likely to benefit from targeted therapeutics in development for this devastating malignancy.

Here, two new technologies were created in parallel: 1) a tailored set of 34 antibodies for single cell mass cytometry of glioblastoma focused on phospho-protein signaling effectors, stem cell proteins, and transcription factors critical to neural development, and 2) an unsupervised cell discovery workflow termed RAPID (<u>R</u>isk <u>A</u>ssessment <u>P</u>opulation <u>Id</u>entification). These technologies were combined to reveal and characterize novel populations of risk stratifying glioblastoma cells.

## Comprehensive patient-specific analysis reveals glioblastoma cells with potentiated signaling and aberrant lineage protein co-expression

Tissue samples were collected from 28 patients with *IDH* wild-type glioblastoma after primary surgical resection (Supplementary Table 1). As of February 2019, the median progression free survival (PFS) and overall survival (OS) after diagnosis were 6.3 and 13 months, respectively, similar to those observed in larger populations of patients undergoing standard therapy ^27^. Resected tissues were immediately dissociated into single-cell suspensions as previously reported ^38^. Cells were stained with a customized antibody panel for mass cytometry designed to capture the expression of known cell surface proteins, intracellular proteins, and phospho-signaling events (Figure 1, 2, Supplementary Table 2, Supplemental Information) that are critical for gliomagenesis and pathogenesis ^32, 39, 40, 30, 34, 35^. Collectively, the antigens included in this panel positively identified >99% of viable single cells within any given tumor sample.

**Figure 1:**
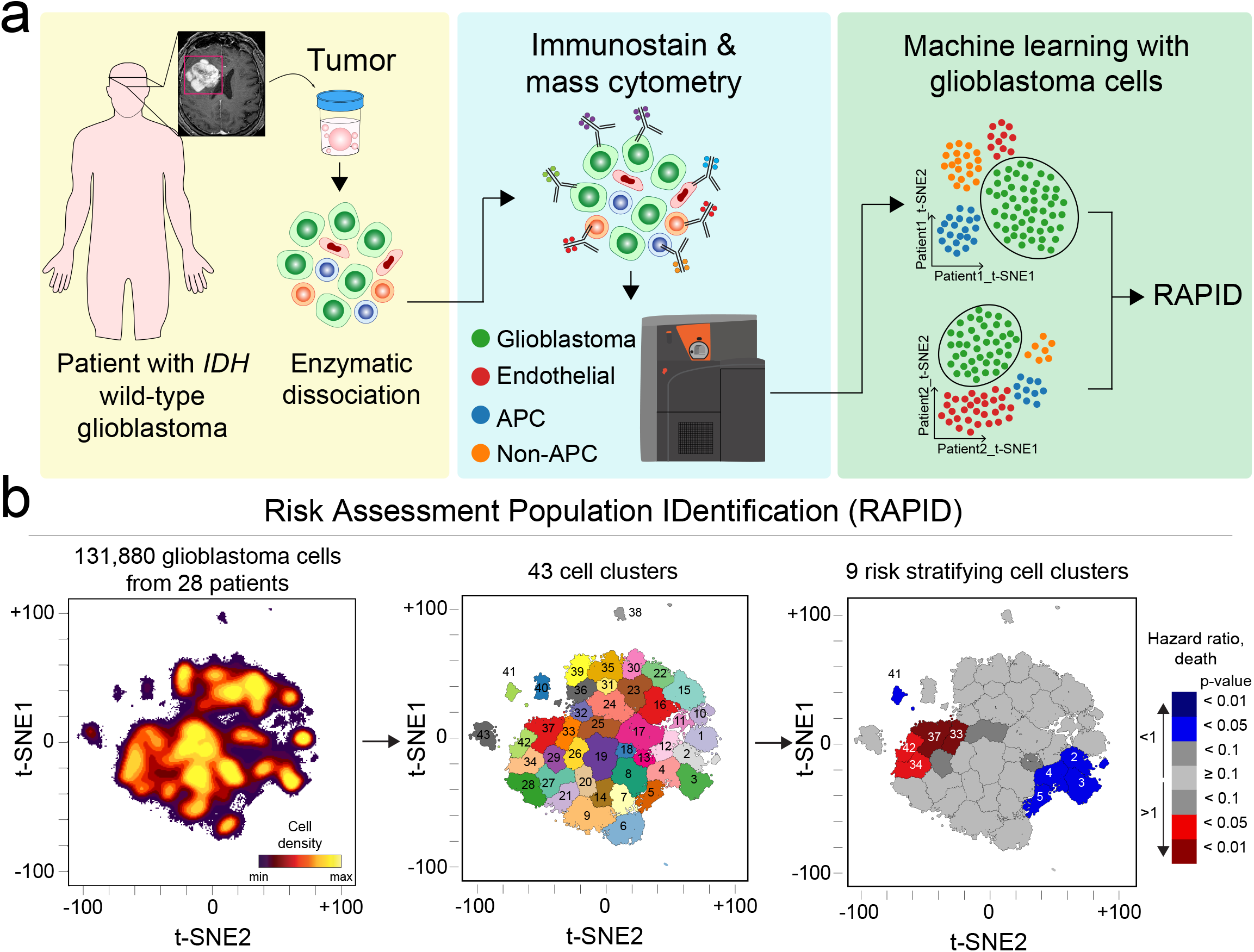
RAPID identifies single cell phenotypes and assesses clinical risk stratification as a continuous variable. (a) Graphic of tumor processing and computational workflow. (b) Glioblastoma cells were identified from 28 patients and computationally pooled for a t-SNE analysis. Cell subsets were automatically identified by FlowSOM and were systematically assessed for association with patient overall or progression-free survival. 43 glioblastoma cell subsets were identified and were color-coded based on hazard ratio of death and p-values (HR>1, red; HR<1, blue) Cell density, FlowSOM cluster, and cluster significance are depicted on t-SNE plots.

**Figure 2:**
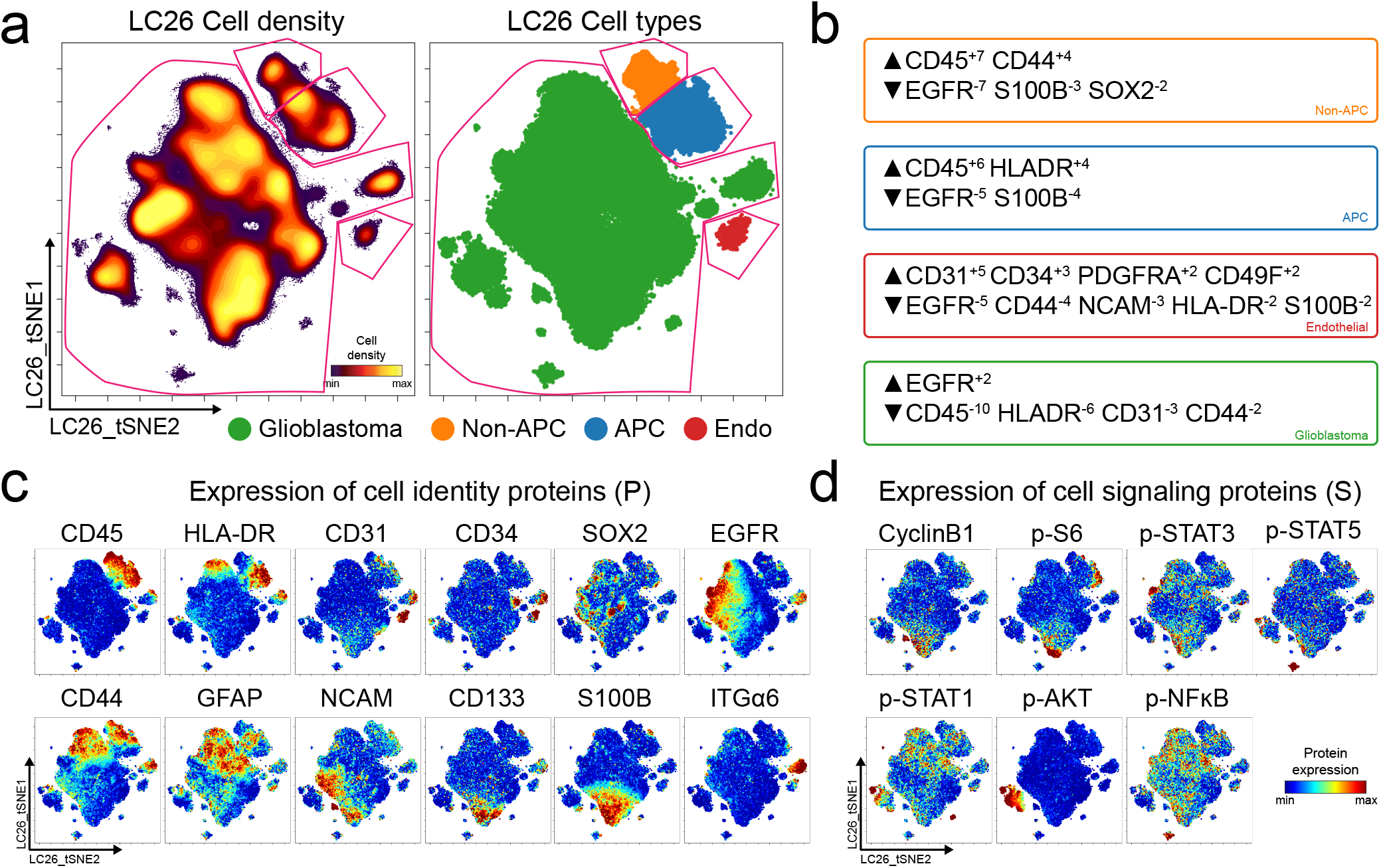
Single-cell quantification of identity proteins and phospho-protein signaling in glioblastoma. (a) t-SNE plots of cell density (left) and major cell types in a patient tumor colored by expert gating (right) for antigen presenting cells (APC, blue), other immune cells (non-APC, orange), endothelial cells (Endo, red), and glioblastoma cells (green) using CD45, CD31, and HLA-DR to identify cells. Pink lines indicate where expert gates were drawn. (b) MEM protein enrichment scores for populations indicated by color in (a) (APC, blue; non-APC, orange; endothelial cells, red; glioblastoma cells, green). (c) Per-cell expression levels of 12 identity proteins, (d) 6 phosphorylated signaling effectors, and proliferation marker cyclin B1 are depicted. Heat indicates protein or phospho-protein expression per cell (scale is specific to each measured feature).

The first round of data analysis was patient-specific and used to computationally isolate the glioblastoma cells from the stromal cells. A patient-specific t-SNE view of single-cell protein expression was generated for all tumor and stromal cells from each patient’s tumor (Supplemental Information). These patient-specific t-SNE maps were generated using 26 of the 34 measured markers ^18^ (Supplemental Table 2). Patient specific t-SNE maps revealed nonglioblastoma populations of immune (CD45^+^) and endothelial (CD45^−^CD31^+^) cells, consistent with prior mass cytometry studies of gliomas ^11, 21, 24^. Non-immune, non-endothelial cells were computationally isolated from each individual patient prior to subsequent downstream analysis of tumor-intrinsic phenotypic parameters (Figure 1, 2). These remaining CD45^−^CD31^−^ cells were labeled as glioblastoma cells. Ion counts for mass-tagged antibody reporters spanned from 0 to nearly 10,000, representing protein expression from a sensitivity limit of around 400 molecules per cell to 1 x 10^7 molecules per cell ^8^. This sensitivity and the ability to capture at least 4,500 live glioblastoma cells from every patient provided excellent statistical power to observe rare cell types representing as little as 1% of the cancer cells (which might themselves be as little as 25% of the tumor cells).

Within a single patient’s tumor, plots of cell density across the t-SNE embedding revealed 5 or more phenotypically distinct subpopulations of glioblastoma cells (Figure 2, Supplemental Information). These intra-tumoral subsets of glioblastoma cells were distinguished by differences in expression of core neural identity proteins and by aberrant co-expression of neural lineage and stem cell proteins. For example, in tumor LC26, abnormal glioblastoma cell subsets were apparent and distinguished by lineage aberrancy. Common abnormal co-expression phenotypes in glioblastoma cells included expression of astrocytic S100B and stem-like CD133 or co-expression of markers associated with different molecular subtypes of glioblastoma, such as mesenchymal (CD44) and classical (EGFR) (Figure 2) ^32^.

Additional intra-tumoral diversity in glioblastoma cells was revealed by quantification of the phosphorylation states of eight signaling effectors (Figure 2; p-STAT5^Y694^, p-STAT3^Y705^, p-S6^S235/S236^, pSTAT1^Y701^, p-NFĸB (p65) ^S529^, p-AKT^S473^, p-ERK1/2^T202/Y204^ .and p-p38^T180/Y182^). Subsets of cells distinguished by abnormal lineage expression typically displayed potentiated basal phospho-protein signaling. For example, simultaneous phosphorylation of S6, STAT5, and STAT3 was commonly observed in glioblastoma cells that expressed S100B, but not in cells that expressed EGFR, GFAP, or CD44 (Supplemental Information). In summary, multiple biologically distinct glioblastoma cells, distinguished by combinations of cellular identity proteins and potentiated signaling features within individual tumor specimens were revealed by per patient analysis.

## RAPID identifies prognostic cell subsets in glioblastoma disease

The second round of data analysis used an equal number of each patient’s glioblastoma cells to create a single, common t-SNE map of glioblastoma cell phenotypes across all patients (N = 131,880 cells; 4,710 cells x 28 patients). Prior to creating this common map, mass cytometry standardization beads were used to remove batch effects and to set the variance stabilizing arcsinh scale transformation for each channel following field-standard protocols ^11, 38, 41^. This common t-SNE map was generated using 24 of 34 measured markers (Supplementary Table 2) and was used for automated analysis of risk stratifying cell subsets.

Once a common, low-dimensional view of all patients was established, the RAPID algorithm used statistical analysis of cell density, feature variance, and population abundance to automatically set all computational analysis parameters. Critically, RAPID was designed to set analysis parameters independent of clinical outcomes. To identify an appropriate number of stable clusters containing phenotypically homogenous cells, RAPID used iteratively executed unsupervised self-organizing maps from FlowSOM ^42^. RAPID repeatedly tested a range of clusters (5-50) and identified the number of clusters that minimized intra-cluster variance for each feature while maintaining cluster stability (see Supplemental Methods). Within the glioblastoma patients examined here, RAPID identified 43 phenotypically distinct glioblastoma cell subsets (Figure 1). RAPID assigned patients to high or low abundance for a given cluster based on a cut-point, set as the interquartile range of the population abundance across the samples (see Supplemental Methods). For example, for cluster 24, the interquartile range was 0.67% to 3.36%, resulting in a cut point of 2.69% (Supplementary Table 4). Those patients with < 2.69% were designated ‘low’ for cluster 24 while those with > 2.69% were assigned to the ‘high’ group. Finally, RAPID applied a univariate Cox survival analysis to determine the correlation between the abundance of tumor cells in each cluster and patient survival outcome.

The output of RAPID, when using t-SNE and FlowSOM, is a PDF containing a color-coded, 2D t-SNE plot depicting all FlowSOM clusters, a 2D t-SNE plot colored by clusters which were significantly associated with patient outcome, and Kaplan-Meier survival estimates of patients for each subset (additional files described in Methods) (Figure 1b).

## Distinct glioblastoma cellular phenotypes associate with patient prognostic outcomes

RAPID identified 43 phenotypically distinct glioblastoma cell clusters. Of these, 7 clusters were considered “universal” because cells from every tumor were observed in these clusters (ranging from 0.02% to 28.05%, Supplemental Table 4). The abundance of the 43 clusters varied extensively across patients. Tumors contained a median of 14 clusters at >1% with a range from 5 cell clusters in LC06 to a maximum of 27 cell clusters LC25. Overall, the presence of a greater number of GBM cell clusters at >1% abundance within a tumor was not observed to be associated with differential survival (ρ=0.047, p=0.812).

In contrast, the abundance of 9 glioblastoma cell clusters was closely correlated with overall survival (Fig. 3,4). Clusters were identified here as prognostic by assessing the hazard ratio (HR) of death in patients who were either high or low for the cell cluster. Negative and positive prognostic clusters were colored red or blue in graphs if they were significantly associated (p<0.05) with an HR that was >1 or <1, respectively.

**Figure 3:**
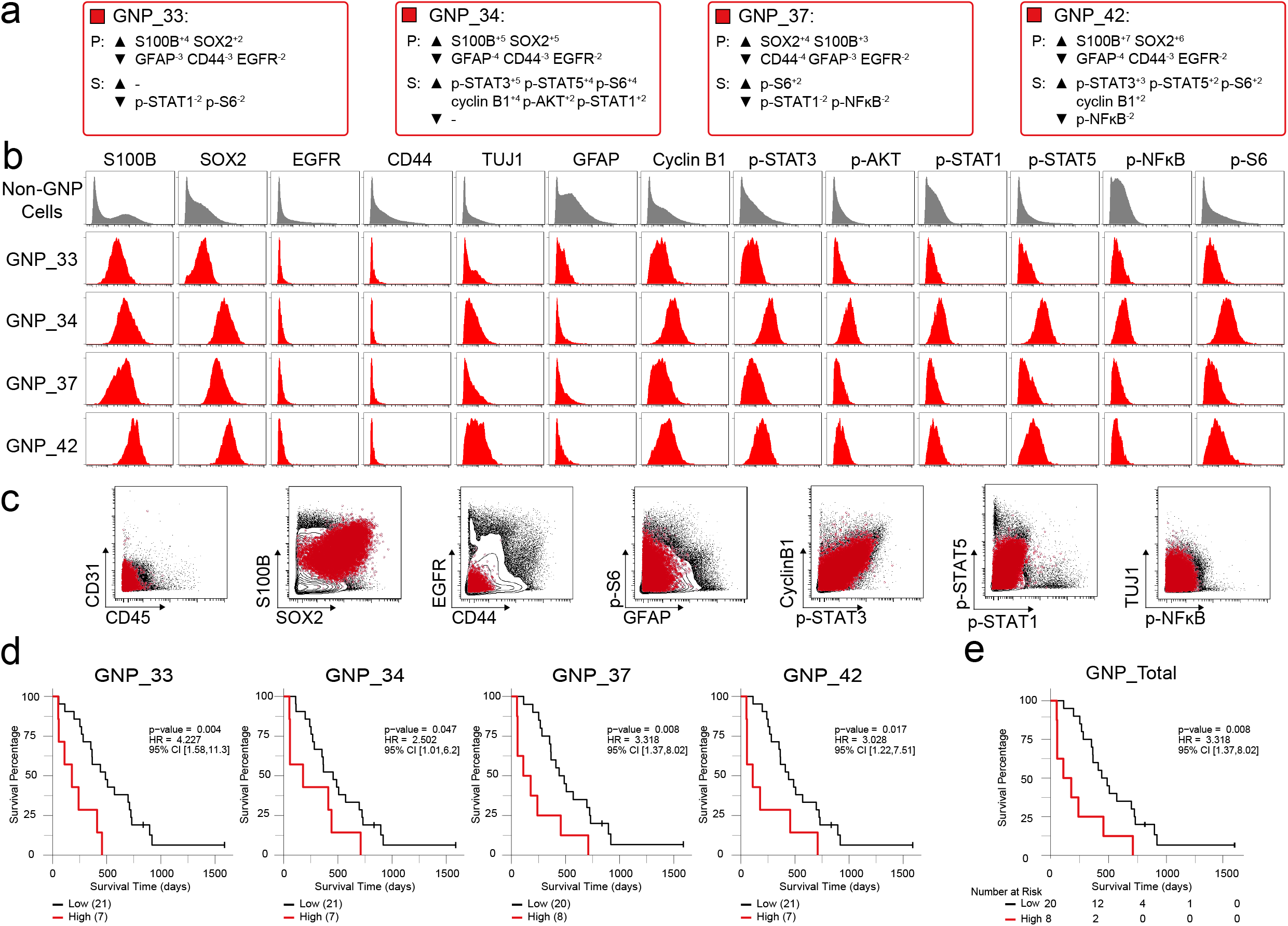
Four cell populations identified by RAPID were negatively associated with patient outcome. (a) Enrichment (upwards arrowhead) or lack (downwards arrowhead) of identity proteins (P) and phosphorylated signaling effectors (S) on Glioblastoma Negative Prognostic cell subsets was quantified using MEM. (b) Histogram plots of each GNP cell subset (red) and all other glioblastoma cells (gray) illustrate the expression of identity proteins and phosphorylated signaling effectors. (c) Combined GNP cell subsets (red circles) were mapped over biaxial plots of all other tumor cells (black contours). (d) For each subset, overall survival was compared between patients with high vs low cell abundance (see Supplementary Methods). (e) Overall survival of patients for high (> 3.1%) total GNP content compared to patients with low (< 3.1%) GNP content.

**Figure 4:**
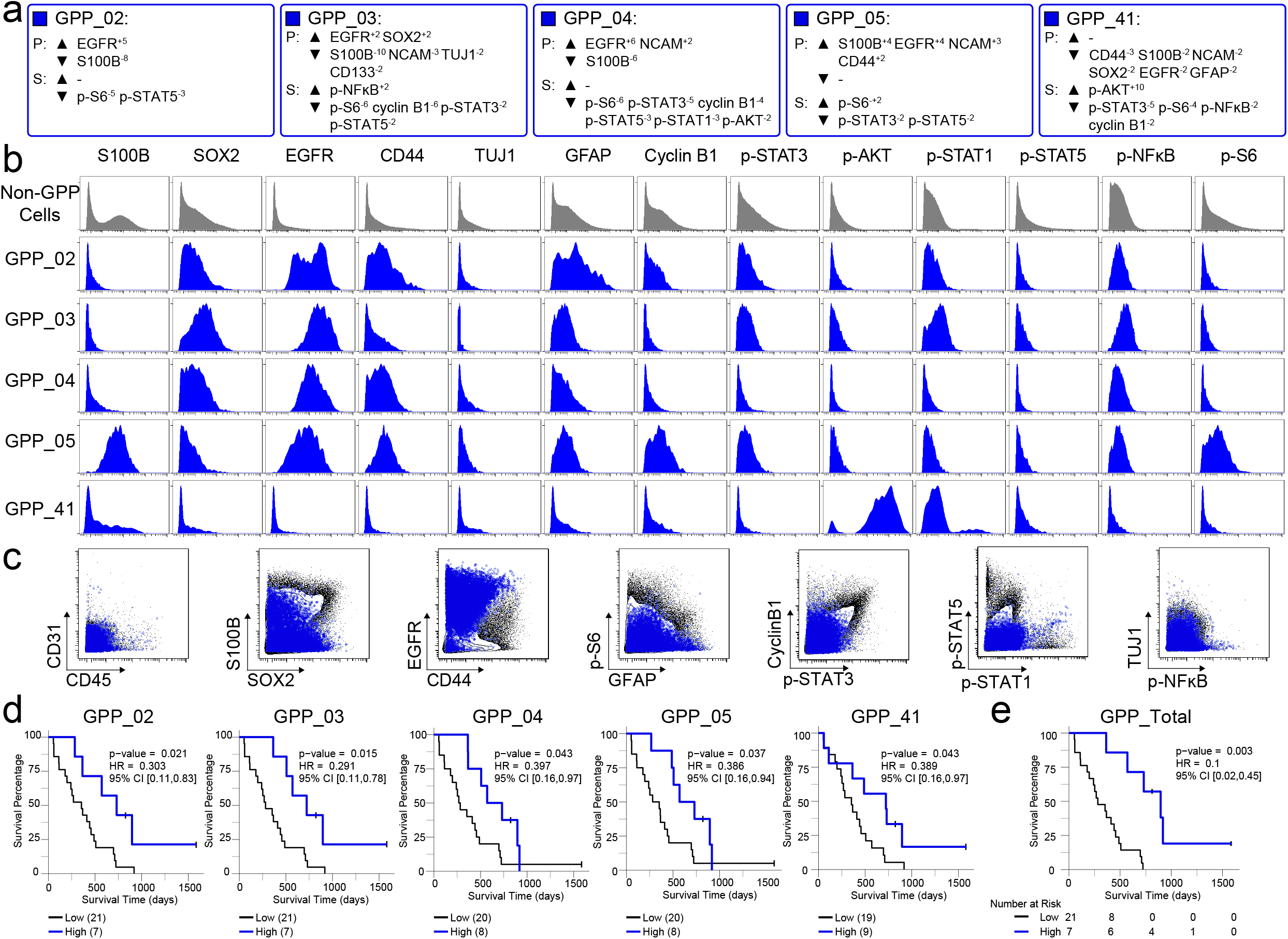
Five cell populations identified by RAPID were positively associated with patient outcome. (a) Enrichment (upwards arrowhead) or lack (downwards arrowhead) of identity proteins (P) and phosphorylated signaling effectors (S) on Glioblastoma Positive Prognostic cell subsets was quantified using MEM. (b) Histogram plots of each GPP cell subset (blue) and all other glioblastoma cells (gray) illustrate the expression of proteins and phosphorylated signaling effectors. (c) Combined GPP cell subsets (blue circles) were mapped over biaxial plots of all other tumor cells (black contours). (d) For each subset, overall survival was compared between patients with high vs low cell abundance (see Supplementary Methods). (e) Overall survival of patients for high (> 8.58%) total GPP content compared to patients with low (< 8.58%) GPP content.

Four Glioblastoma Negative Prognostic (GNP) clusters (red; clusters 33, 34, 37, and 42) and five Glioblastoma Positive Prognostic (GPP) clusters (blue; clusters 2,3,4,5, and 41) were identified (Figure 3, 4). The remaining 34 clusters were not associated with differential prognosis. RAPID was also used to identify glioblastoma cell clusters with differential PFS, as opposed to OS. Assessing PFS can be especially useful for cancers with longer median survival where progression-free survival is the most useful clinical assessment. Of the 43 subsets identified by RAPID, 4 subsets were significantly associated with PFS (subsets 20, 33, and 43 with negative PFS and subset 3 with positive PFS, Supplemental Figure S1).

To determine if the effect of cell subset abundance was continuous and independent of other features known to stratify glioblastoma survival, a multivariate Cox proportional-hazards model analysis was performed incorporating known features and GNP or GPP cell abundance. Known predictors included were age ^43, 44^, *MGMT* promoter methylation status ^45, 46^, and treatment variables including the extent of surgical resection ^47, 48^, therapy with temozolomide ^27^, and radiation ^49, 50^. Multivariate survival analysis of GNP cell abundance on a continuous scale, keeping the other predictors constant, indicated that each 1% increase in GNP cells was associated with an approximately 7% increase in mortality compared to baseline (OS HR=1.07 [95% CI 1.03-1.12], p=0.001). Similarly, a 1% increase in GPP cells was associated with an approximately 6.5% decrease in mortality rate (OS HR=0.935 [0.877-0.997], p=0.04) and an approximately 3.6% decrease in time to tumor progression, as compared to baseline (PFS HR=0.964 [0.930-0.999], p=0.04). When GNP and GPP were assessed simultaneously, abundance of GNP cells was the primary predictor of mortality (OS HR=1.06 [1.01-1.10], p=0.02), while abundance of GPP cells was the primary predictor of time to tumor progression (PFS HR =0.96 [0.93-1.00]; p=0.04). Thus, the abundances of GNP and GPP cell subsets were associated with distinct and contrasting patient outcomes (Figure 3, 4, Supplemental Figure S1), and their predictive value was independent of each other and known prognostic factors of patient survival.

## Enrichment of divergent signaling effectors in prognostic glioblastoma cell subsets

MEM was used in RAPID to quantify the enriched features of the 43 clusters identified by RAPID, including GNP and GPP clusters, in a compact label of cell identity. Protein enrichment was reported on a +10 to −10 scale, where +10 indicates that protein’s expression was especially enriched and −10 indicated that the protein’s expression was excluded from those cells, relative to other glioblastoma cell clusters (Supplemental Figure S2, S3). MEM labels were calculated for both total proteins (P), such as S100B and EGFR, and signaling effectors (S), such as p-STAT5, in the prognostic GNP (Figure 3a) and GPP (Figure 4a) clusters. The MEM label of each cluster is thus an objective description of what makes that population distinct from the other 42 clusters identified by RAPID (Supplemental Figure S3). GNP cells aberrantly coexpressed neural-lineage proteins (astrocytic S100B and stem-like SOX2). Additionally, GNP cells displayed phosphorylation of RTK signaling effectors known to promote cell survival, growth, and proliferation (e.g. p-STAT5, p-S6, p-STAT3, cyclin B1) (Figure 3b). The median and standard deviation of the MEM protein enrichment values for GNP cells included neural lineage determinants (▲S100B^+5±1.6^, SOX2^+4.3±1.7^) and phospho-proteins (▲p-STAT3^+2.8±1.4^, p-STAT5^+2±1.4^) and identified proteins that were specifically lacking in GNP cells relative to other GBM cell clusters (▼EGFR^−2.3±0.1^, GFAP^−3.6±0.8^, CD44^−3.5±0^) (Figure 3). In contrast, GPP cells were positively enriched for EGFR (▲EGFR^+3.7±3.4^) and consistently lacked proliferation (▼cyclin B1^−2.4±2.9^) and pro-survival phospho-proteins (▼p-S6^−3.8±3.2^, p-STAT5^−2±0^, p-STAT3^−3±2^) (Figure 4).

Immune cells were intentionally excluded from initial RAPID analyses and subsequent biaxial gating confirmed that the GNP and GPP subsets did not contain any unexpected residual CD45- or CD31-positive cells (99.50% and 98.64% non-immune, non-endothelial cells, respectively, Figure 3, 4). However, infiltrating immune cells can comprise a large proportion of non-cancer cells in glioblastomas and have highly variable overall abundance across patients ^51–53^. Notably, GPP-high patients’ tumors all contained > 9% CD45^+^ cells, whereas GNP-high patients’ tumors were not observed to contain more than 9% CD45^+^ cells (p < 0.001, Supplementary Figure 4).

## Comparable identification of prognostic glioblastoma cells with different subsampling and dimensionality reduction tools

The cells input to RAPID should be equal in number from each patient in order to remove the possibility that a single patient would disproportionately impact the identification of cell clusters and risk assessment. However, this limits the RAPID analysis to a number of cells equal to the smallest observed from any one patient and it creates the possibility that the cells randomly selected from tumors where many cells were measured might not be representative. For the tumors studied here, the number of glioblastoma cells measured ranged from 4,710 to 330,000 cells per patient.

To test whether the cells sampled for RAPID were representative of the tumor from which they were selected, 9 additional t-SNE analyses were created, each with a different sample of 4,710 cells selected at random, with replacement, from each patient. Each of these 9 t-SNE maps were then used in a new RAPID analysis, creating 10 total analyses (the original and 9 new tests). In these analyses, RAPID identified different numbers of optimal clusters ranging from 18 to 48. Of these, a total of 48 clusters from the 9 new runs were considered prognostic. Because the 10 RAPID analyses ran on different subsampling of cells, the f-measure could not be calculated on a cell-by-cell basis. However, the average f-measure based on patient categorization (GNP high, GNP and GPP low, and GPP high) was 0.79 between t-SNE runs.

Thus, to quantify the degree of similarity between the 48 newly identified clusters and the 9 original GNP and GPP clusters, the root-mean-square deviation (RMSD) in the MEM enrichment values was calculated as a way of determining if the phenotype of the newly identified clusters was stable, even when different cells were sampled from the tumors. GNP subsets from subsequent runs were highly similar to the GNP subsets identified by the initial analysis described above and the same was observed for GPP subsets (Supplemental Figure S5; GNP v GNP average RMSD = 92.5, GPP v GPP average RMSD = 88.2, and GNP v GPP average RMSD = 80.8).

To test whether RAPID could use different types of dimensionality reduction values as input parameters, the algorithm was implemented replacing t-SNE with UMAP (Uniform Manifold Approximation and Projection), a tool that emphasizes both local and global data structure ^19^ RAPID identified 31 populations using UMAP input; 4 of these were prognostic and significantly associated with OS (1 GNP_UMAP_ and 3 GPP_UMAP_) (Figure 5). GNP_UMAP_ MEM scores reflected the characteristic S100B and SOX2 expression observed in the GNP populations along with an active pro-survival basal signaling status. GPP_UMAP_ subsets were similarly defined by co-expression of EGFR and CD44 and a general lack of the measured phosphorylated signaling effectors (Figure 5). When the cells identified using t-SNE were overlaid on the UMAP axes, they occupied similar phenotypic space as UMAP-identified clusters, and vice versa (f-measure for cell assignment to GNP, GPP, or neither = 0.872, Figure 5). Thus, when UMAP was used in the RAPID algorithm, GNP and GPP populations were identified that had comparable phenotypes to those identified previously in t-SNE analyses, confirming that RAPID is not dependent upon a specific dimensionality reduction tool (Figure 5).

**Figure 5:**
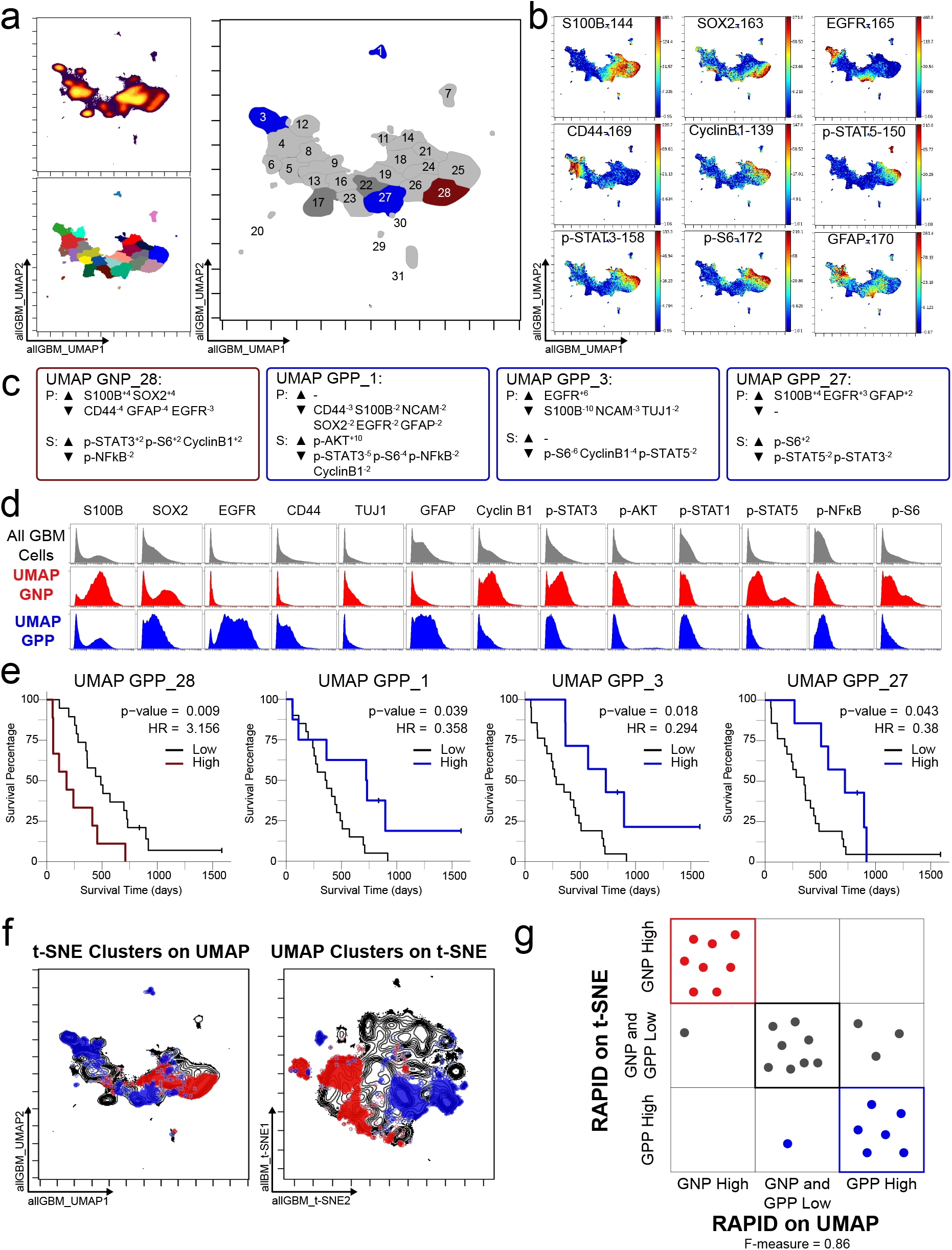
GNP and GPP cells were also identified using dimensionality reduction tool UMAP in the RAPID algorithm. (a) UMAP analysis of 131,880 cells from 28 patients. Upper left plot - heat on cell density; lower left plot – colored by FlowSOM cluster; right plot – colored by GNP(red)/GPP(blue) designation and p-value. (b) Per-cell expression levels of 5 identity proteins, 3 phosphorylated signaling effectors, and proliferation marker cyclin B1 are depicted. (c) Enrichment of identity proteins (P) and phosphorylated signaling effectors (S) of glioblastoma cell subsets was quantified using MEM. GNP and GPP cells are labeled in red and blue, respectively. (d) Histogram analysis depicts the expression of key identity proteins and phosphorylation signaling effectors of GNP (red) and GPP (blue) compared to all other cells (gray, top row). (e) Overall survival curves for four UMAP-identified populations associated with survival. Cox-proportional hazard model was used to determine a hazard ratio (HR) of death. Censored patients are indicated by vertical ticks. (f) GNP (red) and GPP (blue) cells identified via t-SNE (“t-SNE GNP” or “t-SNE GPP”) and UMAP (“UMAP GNP” or “UMAP GPP”) are overlaid on either UMAP or t-SNE axes. Localization of GPP cell subsets identified by one tool in the same region as those identified by the other suggests similar phenotypes across dimensionality reduction methods. (g) Categorization of each patient (dots) based on GNP high (red), GPP high (blue), or neither (gray) according to abundance based on RAPID using t-SNE or RAPID using UMAP (f measure = 0.86).

## Towards tracking clinically distinct glioblastoma cells in the clinic

After patterns are recognized by a machine learning approach, it can be valuable to determine whether the learned features can be identified using simpler models that can be applied by experts or machines to new datasets. One approach is to create a decision tree using one- or two-dimensional gating ^23^, consistent with traditional strategies in immunology and hematopathology. Such gates make the identification of cells computationally less intensive and more pragmatic for wide-spread clinical use and have been previously used in glioblastoma mass cytometry ^24^. Therefore, a traditional, lower-dimensional strategy was developed to use a small number of simple gates to capture the GNP and GPP cell populations (Figure 6).

**Figure 6:**
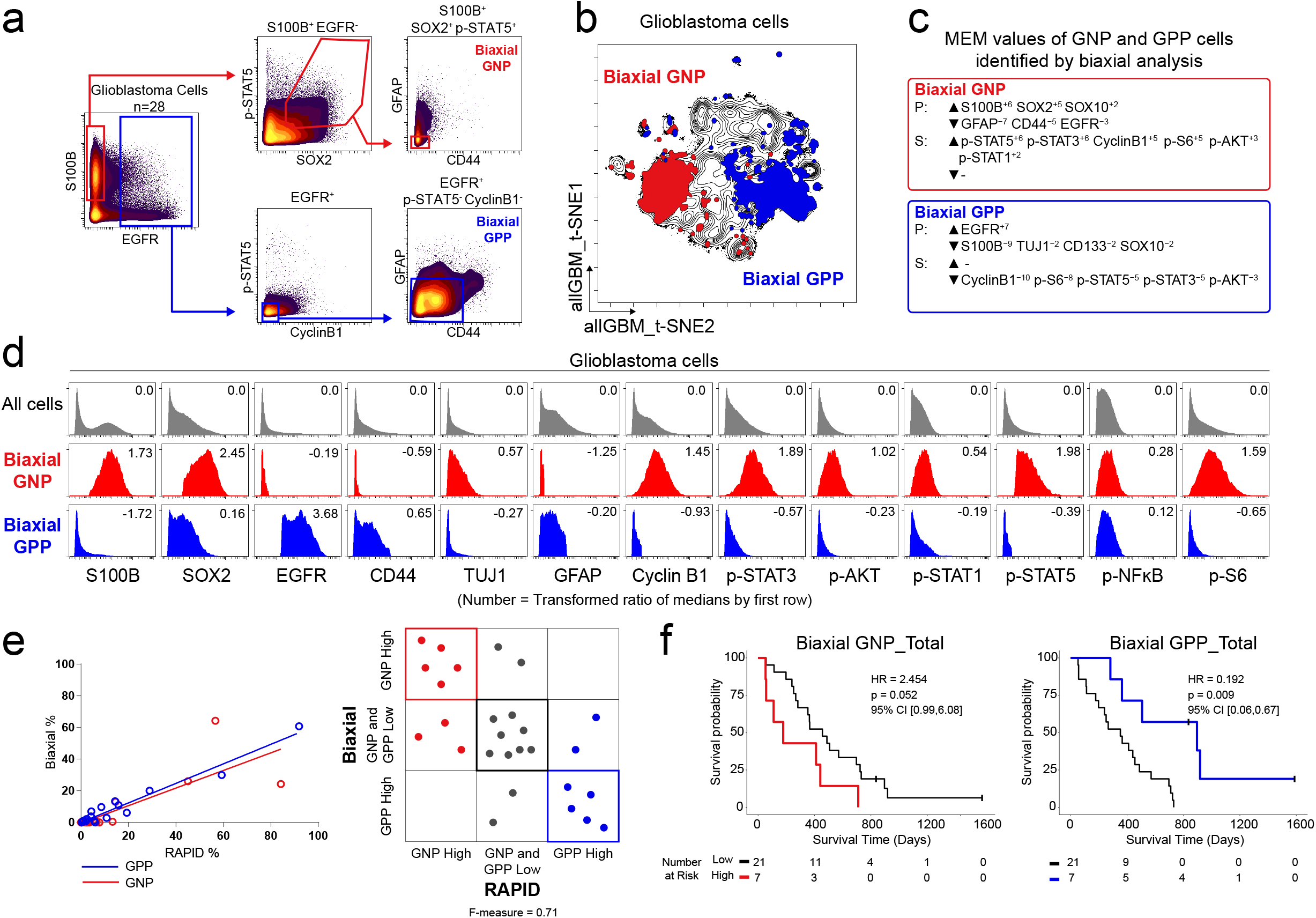
A biaxial gating workflow based on 7 protein markers derived from RAPID effectively identified clinically distinct glioblastoma cell subsets. (a) Biaxial plots demonstrating a sequential gating scheme compatible to a standard clinical flow cytometry workflow. Biaxial GNP and GPP cells were identified using red and blue gates respectively. (b) Biaxial GNP (red) and biaxial GPP (blue) are overlaid over contours of glioblastoma cells from 28 tumors on common t-SNE axes (as in Figure 1). (c) MEM analysis was used to quantify enriched identity proteins (P) and phosphorylated signaling effectors (S) of biaxial GNP and GPP cells. (d) Histogram analysis depicts the expression of key identity proteins and phosphorylated signaling effectors of biaxial GNP (red) and biaxial GPP (blue) compared to all glioblastoma cells (gray, top row). The transformed ratio of medians for each marker on GNP or GPP cells compared to all glioblastoma cells is shown in the upper right corner of each histogram. (e) Correlation between GNP (red, ρ = 0.81) and GPP (blue, ρ = 0.98) population abundance identified via biaxial gating and RAPID (left). Categorization of each patient (dots) based on GNP high (red), GPP high (blue), or GNP and GPP low status according to abundance based on biaxial gating or RAPID. Dots are colored by RAPID categorization (f measure = 0.71). (f) Overall survival of patients for high total GNP content (left) or GPP content (right) compared to patients with low GNP or GPP content based on biaxial gating.

A population that was consistent with both the phenotype and risk stratification of GNP cells was identifiable using 3 gates and 6 proteins (S100B, EGFR, SOX2, p-STAT5, GFAP, and CD44, Figure 6). Similarly, GPP cells could be identified with 3 gates and 6 proteins (S100B, EGFR, cyclin B1, p-STAT5, GFAP, and CD44, Figure 6). This gating scheme accurately captured both GNP and GPP cells (f-measure of 0.826 for categorizing cells as GNP, GPP, or neither using RAPID cell populations as truth and biaxial gating as test). GNP and GPP cells identified by traditional gating were also mapped back onto the t-SNE axes and largely occupied the same regions of the biaxial t-SNE map as the cells identified by the RAPID algorithm (Figure 6). The cells identified by traditional gating were quantitatively comparable in their phenotype, as seen by a comparable MEM label for the gating identified GNP and GPP cell subsets (Figure 6). Thus, a simple gating model of GNP or GPP cell identity successfully recovered GNP and GPP cells by assessing only 7 total key features observed to be enriched following RAPID. This indicated that, once revealed, GNP and GPP cell subsets were phenotypically cohesive in a traditional cell biological sense and could be reliably quantified by traditional approaches compatible with standard clinical flow cytometric profiling.

## Discussion

RAPID is a novel automated workflow that identifies cell subpopulations associated with patient outcomes. The RAPID workflow automatically assigned single tumor cells from *IDH* wild-type glioblastomas into computational clusters based on phenotypic similarity, generated a quantitative phenotypic descriptor of each population, and determined the correlation between the abundance of populations and clinical outcomes. Two significant glioblastoma cell types were identified: Glioblastoma Negative Prognostic (GNP) cells, characterized by high expression of S100B, SOX2, p-STAT3, and p-STAT5, were associated with a decreased overall survival, while Glioblastoma Positive Prognostic (GPP) cells, characterized by high expression of EGFR and CD44, were associated with longer overall survival. Critically, therapeutically targetable signaling events were identified as a signature of prognostic cell populations identified by RAPID, suggesting potentially novel therapeutic strategies for patients with these characteristics.

High-dimensional cytometry was critical to revealing novel prognostic glioblastoma cells in two ways. First, assessment of a large number of cells per tumor enabled the use of an unsupervised approach in the identification of rare and novel cell subsets across patients. Second, per-cell quantification of phosphorylated signaling effector proteins revealed potential mechanisms of tumor cell regulation that are not inherently apparent in bulk tumor data. Supervised analysis of single cell data has previously uncovered signaling events tied to patient survival in hematologic malignancies ^3, 4, 7^. To our knowledge, our findings are the first to reveal a similar connection in a solid malignancy using an automated, unsupervised approach, reinforcing the importance of cell signaling in multiple human malignancies

Although other workflows and algorithms have been developed to identify cell populations of interest in cancer samples (CITRUS ^14^, DDPR ^6^, Phenograph ^4^, Cytofast ^54^), these largely require a level of prior knowledge which may not always be available, especially for solid tumors. For example, Levine et al. used Phenograph ^4^ and an understanding of the coupling of surface markers and signaling status in healthy bone marrow to classify negative prognostic leukemia cells. Similarly, a map of the healthy developmental lineage was instrumental in using DDPR to identify features of negative prognostic leukemia cells ^6^. Supervised methods, including CITRUS and Cytofast, require that samples to be grouped at the beginning of the analysis before generating an overview of cell cluster phenotypes in cytometry data ^54^. These methods, however, require that the data have already been clustered and that each sample be prospectively assigned to a group, whereas RAPID enables analysis with continuous, ungrouped data. In studies of diseased human tissue, it is difficult to anticipate the number of expected unique phenotypic subsets and once identified, these subsets can be challenging to manually annotate as they may reasonably be assigned to one or more cell-type categories (this study and ^29^). It is particularly valuable to be independent of prior knowledge of expected cell clusters in studies of diseases like primary glial tumors, where healthy samples are infrequently obtained and the developmental lineage is largely quiescent. RAPID is designed to be free from supervision in the identification of the number of clusters and also in the assessment of cluster abundance in tumors. RAPID also employs MEM to automatically provide a quantitative description of the features which are most selectively enriched on each cell cluster. Furthermore, RAPID is modular, such that different dimensionality reduction tools (t-SNE, UMAP) can be used with different clustering algorithms (dbSCAN, FlowSOM) within the workflow. A benefit of RAPID is the streamlined, unsupervised application of these tools such that a user can input raw data files (for example, FCS files from cytometry platforms) or equivalent data types in conjunction with patient survival data, and RAPID will output quantitatively described cell clusters and their significance with respect to patient outcome. For the data set used in this study (131,880 cells), RAPID ran in 15 minutes from start to finish.

In this study, RAPID analysis of glioblastoma patient samples demonstrated a link between altered signaling and possible abnormal lineage programs in glioblastoma ^55^. The GNP signature was defined by abnormal neural development features and simultaneous high basal phosphorylation of multiple signaling effectors downstream of RTKs (Fig. 3). STAT5 signaling, a common feature of all GNP cell subsets, is required in development of many tissues to block apoptosis and drive cell cycle entry ^2^. For example, p-STAT5 is an essential feature of negative prognostic acute myeloid leukemia signaling profiles ^3, 4^. Here, RAPID identified the connection between p-STAT5 and glioblastoma outcome, previously unidentified in primary patient samples. STAT3 and S6 phosphorylation, identified here in GNP cells, agreed with prior studies indicating the importance of p-STAT3 in T cell suppression ^56^ and mTOR-dependent signaling in tumor formation ^57, 58^. These phosphorylation signaling events in GNP cells should be explored as a potential therapeutic target and a biomarker of therapy response.

In contrast, the time-to-progression-prolonging GPP signature was defined by EGFR and CD44 co-enrichment, diminished evidence of proliferation, and specific lack of STAT5 phosphorylation. Previous molecular subtyping predicts EGFR expression in the classical subset of tumors and CD44 expression in mesenchymal tumors ^32^. As these studies were based on bulk tumor data, cells co-expressing EGFR and CD44 (classified as GPP cells in this study) may have been missed; single glioma cells have been shown to co-express pro-tumor receptors ^29, 36^. Genetically, glioblastomas commonly have amplified *EGFR* ^31, 32^; however, we noted examples of tumors with robust EGFR amplification that contained both high and low percentages of GPP cells (data not shown), highlighting the importance of measuring protein expression in addition to genomic content. Although EGFR signaling through mTOR and EGFRvIII has been linked with increased p-S6 and p-STAT3/5 respectively, we did not observe these associations in the GNP or GPP subsets ^59, 60^. Instead, these cells showed enrichment of p-NFĸB (Fig. 3), a transcription factor that activates pro-apoptotic programs ^61, 62^.

Recent studies have revealed significant variation in immune cell abundance and relative proportions of immune cell subsets across glioblastoma ^63, 64^. Here, unfavorable GNP cells were associated with diminished tumor-infiltrating immune cells and GPP cells were associated with higher proportions of immune cells in the tumor microenvironment. These results invite the question of whether an altered immune microenvironment precedes development of an aggressive glioblastoma or whether more aggressive tumors suppress anti-tumor immunity. These findings argue that immunotherapy is likely to be more efficacious in tumors containing GPP cells, but that additional research is needed to understand whether GNP cells directly suppress microglia or immigrant leukocytes.

The GNP and GPP subsets correlated with survival independent of the effects of other widely accepted prognostic factors (age ^43, 44^, *MGMT* promoter methylation status ^45, 46^, and treatment including extent of surgical resection ^47, 48^, therapy with temozolomide ^27^, and radiation ^49, 50^). These cells were identified in pre-therapy, untreated patient samples, suggesting that these phenotypes are linked to biological mechanisms of therapy response or tumor detection by the immune system. Future studies of recurrent glioma samples would illuminate the persistence of these populations. If GNP subsets have the capacity to evade therapy and retain their active proliferation properties, recurrent tumors would be expected to contain higher proportions of GNP cells and have a more uniform phenotype. Although much has been made of loss or gain of genetic aberrations post-temozolomide and radiation therapies, little is known about signaling in recurrent tumor cells and thus it is unclear if clonal evolution and/or a shift in activated phospho-proteins is necessary for tumor cell survival and repopulation. Other factors have recently been shown to correlate with patient outcomes including the location of the tumor with respect to the largest neural stem cell niche in the adult brain, the ventricular-subventricular zone (V-SVZ) ^65^. In the future, one fascinating question will be to determine whether V-SVZ contacting tumors, which correlate with worse outcomes, contain more cells with a GNP-like phenotype and fewer GPP-like cells.

Critically, these discoveries using RAPID led to a development of a lower-dimensional pipeline which can be immediately adopted for clinical stratification. Moreover, the combination of single-cell snapshot proteomics and the automated RAPID algorithm can be immediately applied to the discovery of critical onco-signaling events in other types of intractable human malignancies, providing a needed complement to genomic classification.

## Supporting information

Supplemental Single Cell Patient Information

## Acknowledgements

We thank the Irish and Ihrie labs at Vanderbilt University for helpful discussions, and Dr. Christopher V.E. Wright for his invaluable suggestions during the preparation of this manuscript.

## Author contributions

N.L., R.A.I., and J.M.I. designed the study. N.L. and J.S. performed experimental work. J.S., S.M.B, N.L, A.M.M., R.A.I., and J.M.I. performed data analysis, developed figures, and wrote the manuscript. A.M.M. and S.M.B. compiled patient data and performed survival analysis. S.M.B, A.M.M., K.E.D., A.R.G., and J.M.I. developed R scripts for data analysis and visualization. K.D.W., R.C.T., and L.B.C. provided clinical samples. B.C.M. provided pathological diagnosis and tumor molecular status. R.A.I. and J.M.I. provided financial support. All authors contributed in reviewing the manuscript. Correspondence and requests for materials should be addressed to.

## Materials and Methods

### Patient samples

Surgical resection specimens of 28 IDH-wildtype glioblastomas collected at Vanderbilt University Medical Center between 2014 and 2016 were processed into single cell suspensions following an established protocol ^38^. Only samples that were confirmed to be IDH-wildtype glioblastomas by standard pathological diagnosis were used. All samples were collected with patient informed consent in compliance with the Vanderbilt Institutional Review Board (IRBs #030372, #131870, #181970), and in accordance with the declaration of Helsinki.

### Patient characteristics and collection of clinical data

All patients were adults (≥ 18 years of age) at the time of their maximal safe surgical resection of their cerebral (supratentorial) glioblastomas. Extent of surgical resection was independently classified as either gross total or subtotal resection by a neurosurgeon and a neuroradiologist. Gross total resection was defined as agreement by both viewers of no significant residual tumor enhancement on patients’ gadolinium-enhanced magnetic resonance imaging (MRI) of the brain obtained within 24 hours after surgery. All patients were considered for treatment with postoperative chemotherapy (temozolomide) and radiation according to the standard of care ^27^, after determination of *MGMT* promoter methylation status by pyrosequencing (Cancer Genetics, Inc., Los Angeles, CA, USA). Multiplex polymerase chain reaction (PCR) was used to determine *IDH1/2* mutational status. Patients’ postoperative course was followed until February 2019, noting time to first, definitive radiographic progression or recurrence of glioblastoma as agreed upon by the treating neuro-oncologist and neuroradiologist, and the time to patients’ death. All deaths were deemed to be due to the natural course of patients’ glioblastoma. Median overall survival of the analyzed 28 patients with IDH wild-type glioblastoma was 388.5 days (13 months) and median PFS was 187.5 days (6.3 months), which is typical for the disease^26, 27^.

### Mass cytometry analysis

Cells derived from patient samples were prepared as previously described ^38^. A multi-step staining protocol was used, which included 1) live surface stain, 2) 0.02% saponin permeabilization intracellular stain, and 3) intracellular stain after permeabilization with ice-cold methanol (Supplementary Table 2). After staining, cells were resuspended in deionized water containing standard normalization beads (Fluidigm)^53^, and collected on a CyTOF 1.0 instrument located in the Mass Cytometry Core Facility at Vanderbilt University. Rhodium viability stain and cleaved caspase-3 antibody were included in staining to exclude non-viable and apoptotic cells, respectively. Detection of total histone H3 was used to identify intact, nucleated cells ^24^. A 32-dimensional mass cytometry antibody panel was used to analyze over 2 million viable cells from 28 tumors (ranging from 4,860 to 336,284 cells per tumor). Data were normalized with MATLAB-based normalization software ^53^, and were arcsinh transformed (cofactor 5), prior to analysis using the Cytobank platform ^66^.

### Implementation of RAPID in R

FCS files for each patient sample (28) containing only cells of interest (non-immune, non-endothelial cells) were input in R. Cell subset identification was performed using the previously published FlowSOM R package ^42^. t-SNE values (t-SNE1_glioblastoma and t-SNE2_glioblastoma) from t-SNE (or UMAP) analysis of CD45^−^CD31^−^ glioblastoma cells from 28 patients were used as parameters for cell subset clustering. Within the RAPID workflow, the optimal number of clusters was determined by first identifying, for each feature, the smallest number of clusters that minimizes the intra-cluster signal variance for that feature. Then, the optimal cluster number of the data set was determined by taking the median of the optimal numbers for each individual feature. Once the cluster number was determined, the abundance of cell subsets and their clinical significance was assessed using outcome-guided analysis. Patients were divided into Low and High groups, based on the distribution (interquartile variance) of the abundance of a given cell subset across the cohort. A univariate Cox regression analysis was then used to estimate the effect size (hazard ratio, HR, of death) on survival and quantify its statistical significance with a p-value. The RAPID program output included: 1) two t-SNE (or UMAP) plots (.png), one color coded by each FlowSOM cluster and one color coded by prognostic status and p-value; 2) Kaplan-Meier survival curves for cell subsets; 3) .txt files of MEM and Median values for each feature, enrichment scores, and IQR values; 4) a new FCS file with File ID, cluster ID, and prognostic status for each cell; and 5) an .rds file with survival statistics for each cluster. In this study, abundance of Glioblastoma Negative Prognostic (GNP) and Glioblastoma Positive Prognostic (GPP) cells in each tumor was quantified as percentages per total glioblastoma cells (i.e. immune and endothelial cells were already excluded). MEM analysis was performed in R, using the previously published R package ^21^. In short, MEM captured and quantified cell subset-specific feature enrichment by scaling the magnitude (median) differences between clusters, depending on the spread (interquartile variance, IQR) of the data. These values were then computed in comparison to the remaining cells in a given dataset. MEM values were interpreted as either being positively enriched (▲, UP positive values) or negatively enriched (▼, DN negative values). The variation of a given cellular feature across GNP or GPP cell subsets was quantified as ± standard deviations (SD).

### Survival and statistical analysis

Time from surgical resection to death (overall survival, OS) and time from surgical resection to the initial radiographic recurrence or death before radiographic assessment (progression free survival, PFS) were plotted and analyzed in R. Survival time points were censored if, at last follow up, the patient was known to be alive or had not had radiographic progression. Differences in the survival curves were compared using the Cox univariate regression model, reporting a hazard ratio (HR) between the survival curves.

A Cox proportional-hazards regression model was created to assess the influence of GNP and GPP cells on OS and PFS as continuous variables while accounting for other factors known to affect survival, including age at diagnosis, *MGMT* promoter methylation status, extent of surgical resection (EOR), treatment with temozolomide (TMZ), and radiation (XRT). The hazard model can be written as:

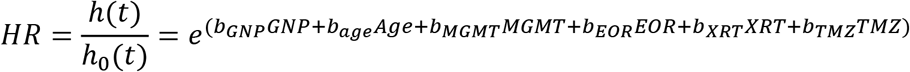

where 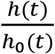 represents the ratio of hazard comparing the risk of death at time *t* to the baseline hazard (obtained when all variables are equal to zero) and *e^b_x_^* represents the hazard ratio of variable *x*. The data were fit using R software, version 3.5 (R foundation for Statistical Computing, Vienna, Austria). The proportional-hazards assumption was tested in all multivariate models and supported by a non-significant relationship between Schoenfeld residuals and time for each covariate included in the model (p > 0.38; degree of freedom = 1) and the overall model (p = 0.96; degrees of freedom = 6 and 7). Statistical significance α was set at 0.05 for all statistical analyses, one- or two-tailed noted in figure legends.

An f-measure was used to quantify the level of agreement between classifications of patients or cells between alternative analysis strategies as wells as multiple RAPID iterations. The f-measure is the harmonic mean of the precision and recall given by the equation F = 2 * (Precision * Recall) / (Precision + Recall) where Precision = True Positive / (True Positive + False Positive) and Recall = True Positive / (True Positive + False Negative). An f-measure of 1 indicates perfect agreement between two different strategies or iterations as opposed to an f-measure of 0 which would mean no agreement between classifications of patients or cells from two strategies or iterations. Patients could be classified as GNP high, GNP and GPP low, or GPP high, while cells were classified as GNP, GPP, or neither. To calculate the f-measure of patient categorization, the classification of the 28 patients into the three prognostic groups from the t-SNE implementation of RAPID was used as the reference point from which to compare patient classification resulting from the UMAP implementation of RAPID or the biaxial gating strategy. Similarly, the stability of the RAPID workflow in assigning cells to GNP, GPP, or nonsignificant clusters was tested by using the t-SNE implementation of RAPID (FlowSOM seed 38) as the reference from which to compare 100 iterations of RAPID (random FlowSOM seed per iteration). Calculation of the f-measure was implemented using R software, version 3.5.

### Data availability

Files will be made available in FlowRepository upon publication following peer review.

---

**Supplemental Figure S1:**
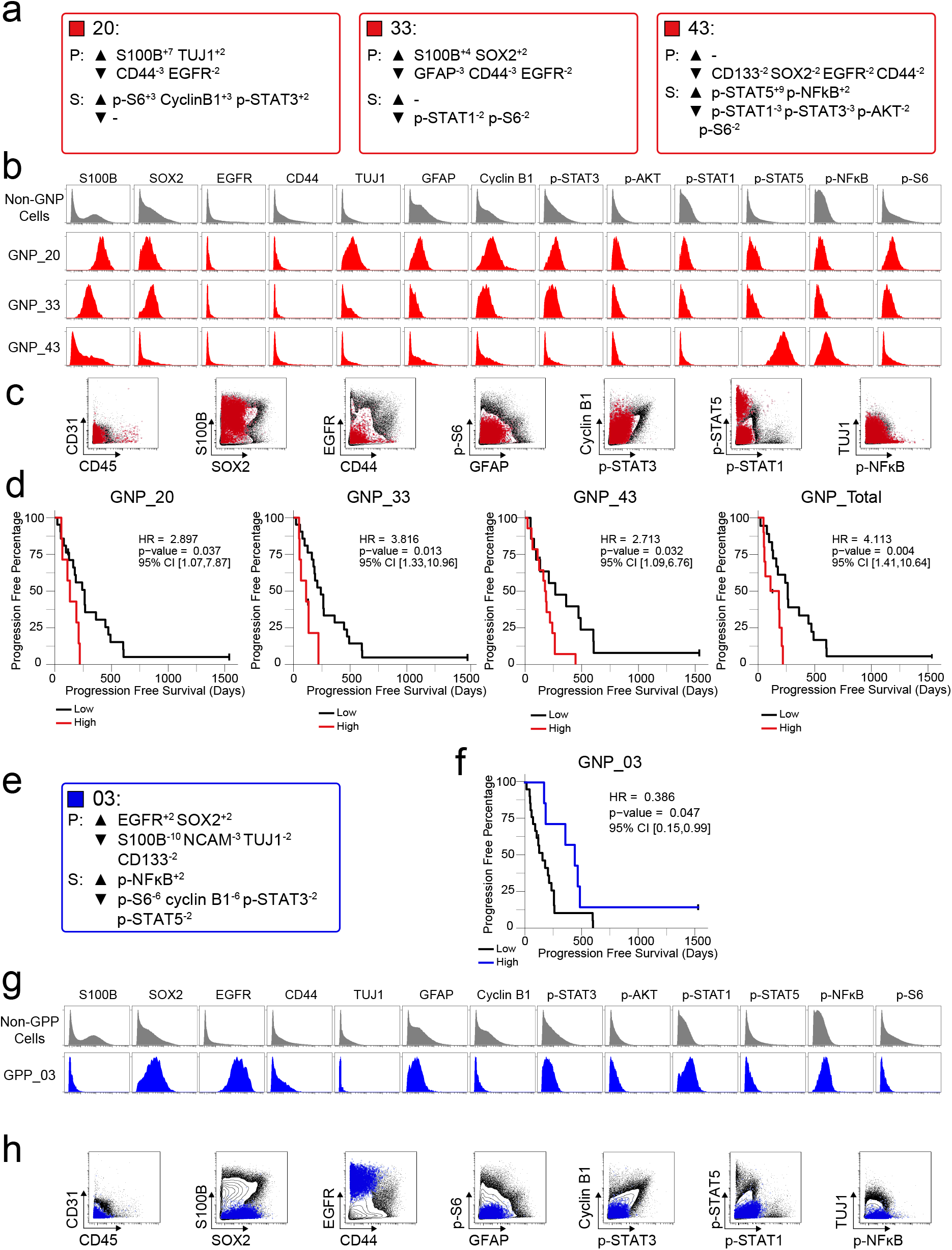
RAPID identified four populations associated with time to patient progression. (a) Enrichment of identity proteins (P) and phosphorylated signaling effectors (S) of PFS GNP cell subsets was quantified using MEM. (b) Histogram plots of each GNP cell subset (red) and all other glioblastoma cells (gray) illustrate the expression of proteins and phosphorylated signaling effectors. (c) Combined GNP cell subsets (red circles) were mapped over biaxial plots of all other tumor cells (black contours). (d) For each subset, progression free survival was compared between patients with high vs low cell abundance (see Supplementary Methods). (e) Enrichment of identity proteins (P) and phosphorylated signaling effectors (S) of the PFS Glioblastoma Positive Prognostic cell subset was quantified using MEM. (f) Progression free survival was compared between patients with high vs low GPP cell abundance (g) Histogram plots of the GPP cell subset (blue) and all other glioblastoma cells (gray) illustrate the expression of proteins and phosphorylated signaling effectors. (h) The GPP cell subset (blue circles) was mapped over biaxial plots of all other tumor cells (black contours).

**Supplemental Figure S2:**
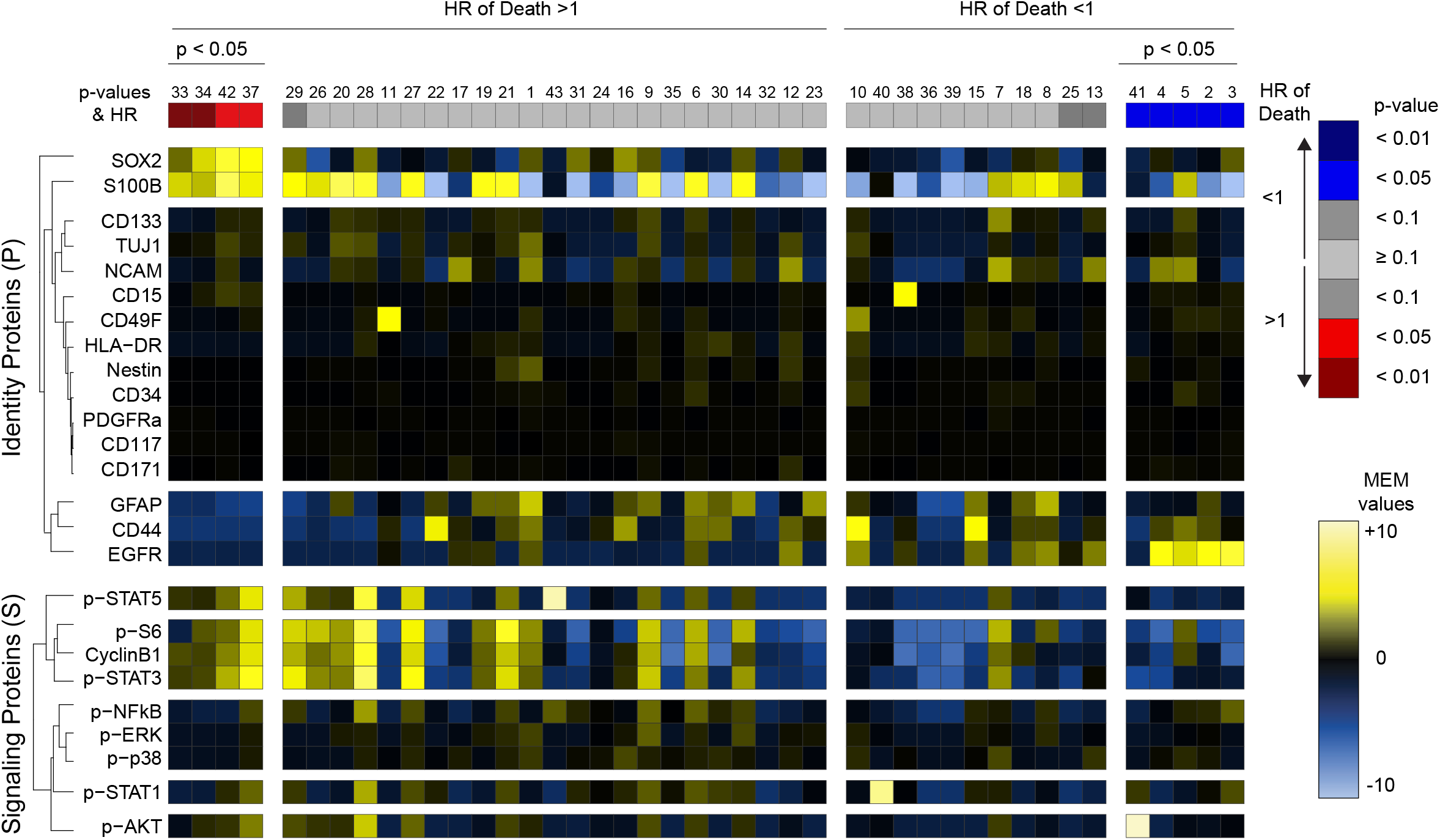
Glioblastoma cell subsets showed differential enrichment of identity proteins and phosphorylated signaling effectors. Forty-three glioblastoma cell subsets automatically identified by FlowSOM are arranged according to their associations with overall survival (HR>1, left; HR<1, right) and statistical significance of that association (p-values). A heatmap represents the MEM values of glioblastoma cell subsets (columns). GNP cells are labeled in red, while GPP cells are labeled in blue. Hierarchical clustering was performed based on MEM values and is depicted on the left of the heatmap for measured features. HR = hazard ratio of death.

**Supplemental Figure S3:**
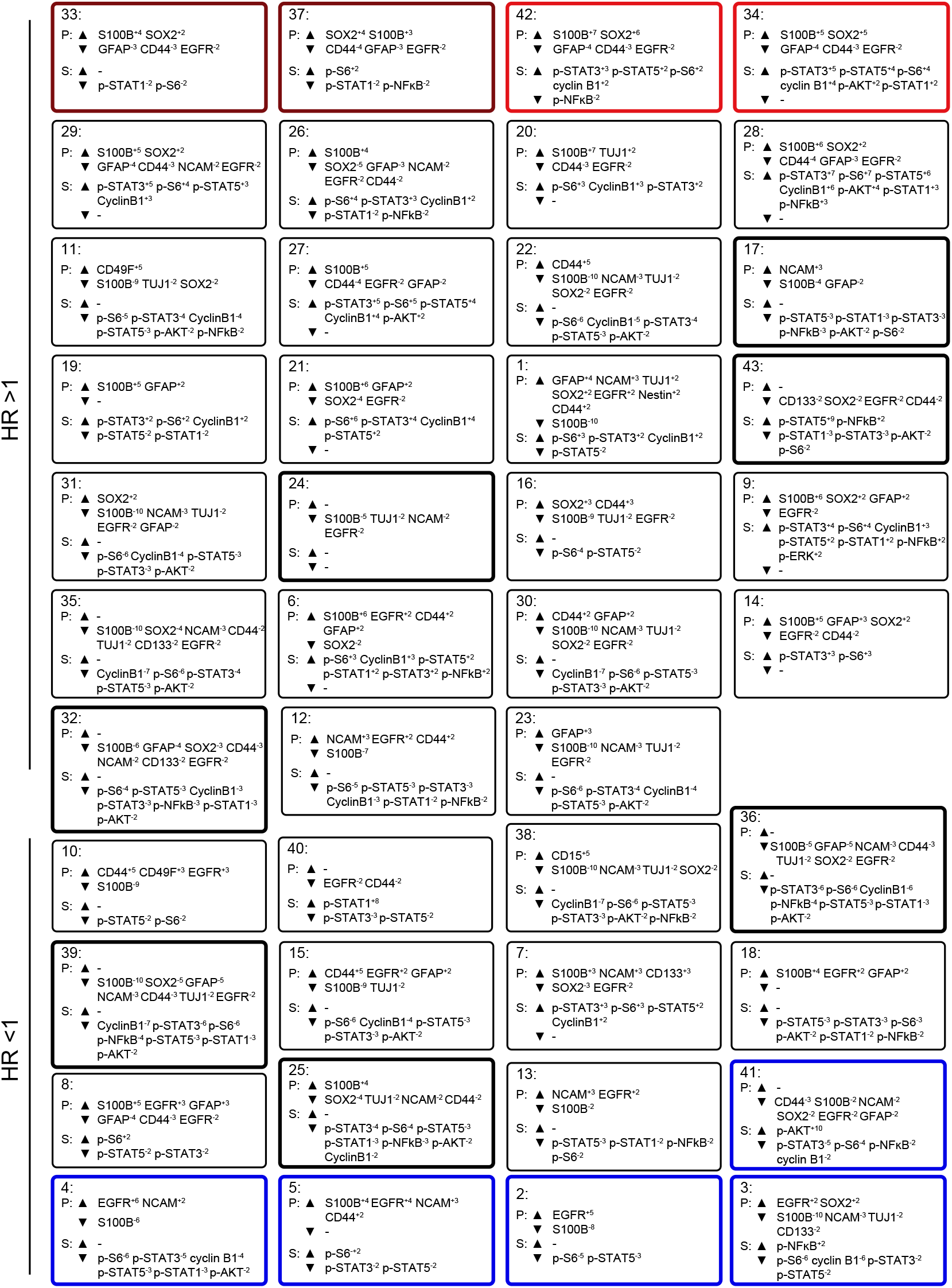
Quantitative MEM labels of the enriched identity proteins and signaling features of all glioblastoma cell subsets identified by RAPID. Enrichment of identity proteins (P) and phosphorylated signaling effectors (S) of glioblastoma cell subsets identified by RAPID was quantified using MEM. GNP and GPP cells are labeled in red and blue, respectively. Populations detected in every patient sample (abundances ranging from 0.02% to 28.05) are outlined in bold.

**Supplemental Figure S4:**
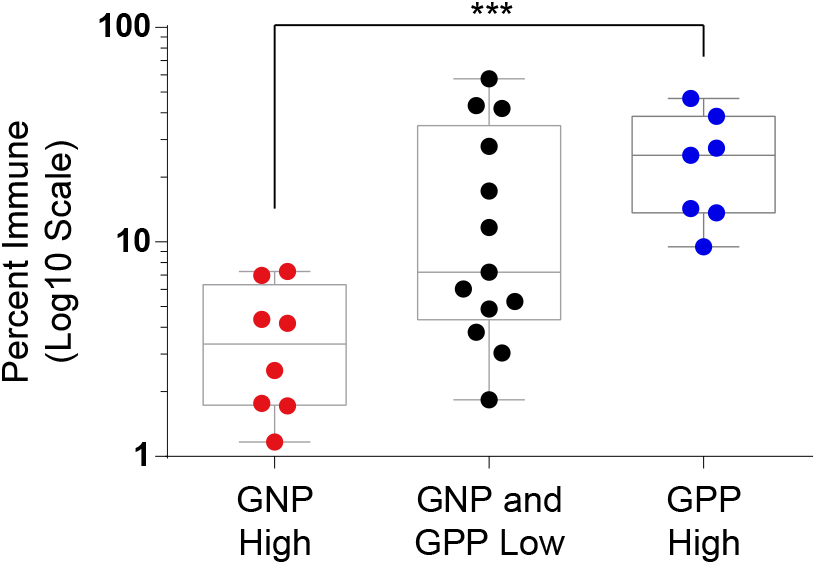
Abundance of immune cells correlated with the abundance of prognostic cell subsets. Box and whisker plot of immune abundance (%, log10 scale) on the y-axis and patients divided into three groups: GNP high (red, >3.1% GNP cells), GPP high (blue, >8.58% GPP), or GNP and GPP low (gray). Box encompasses the 25^th^ to 75^th^ percentile, gray horizontal line indicates the median, and whiskers extend to the minimum and maximum values. *** p=0.0008, two-tailed t-test.

**Supplemental Figure S5:**
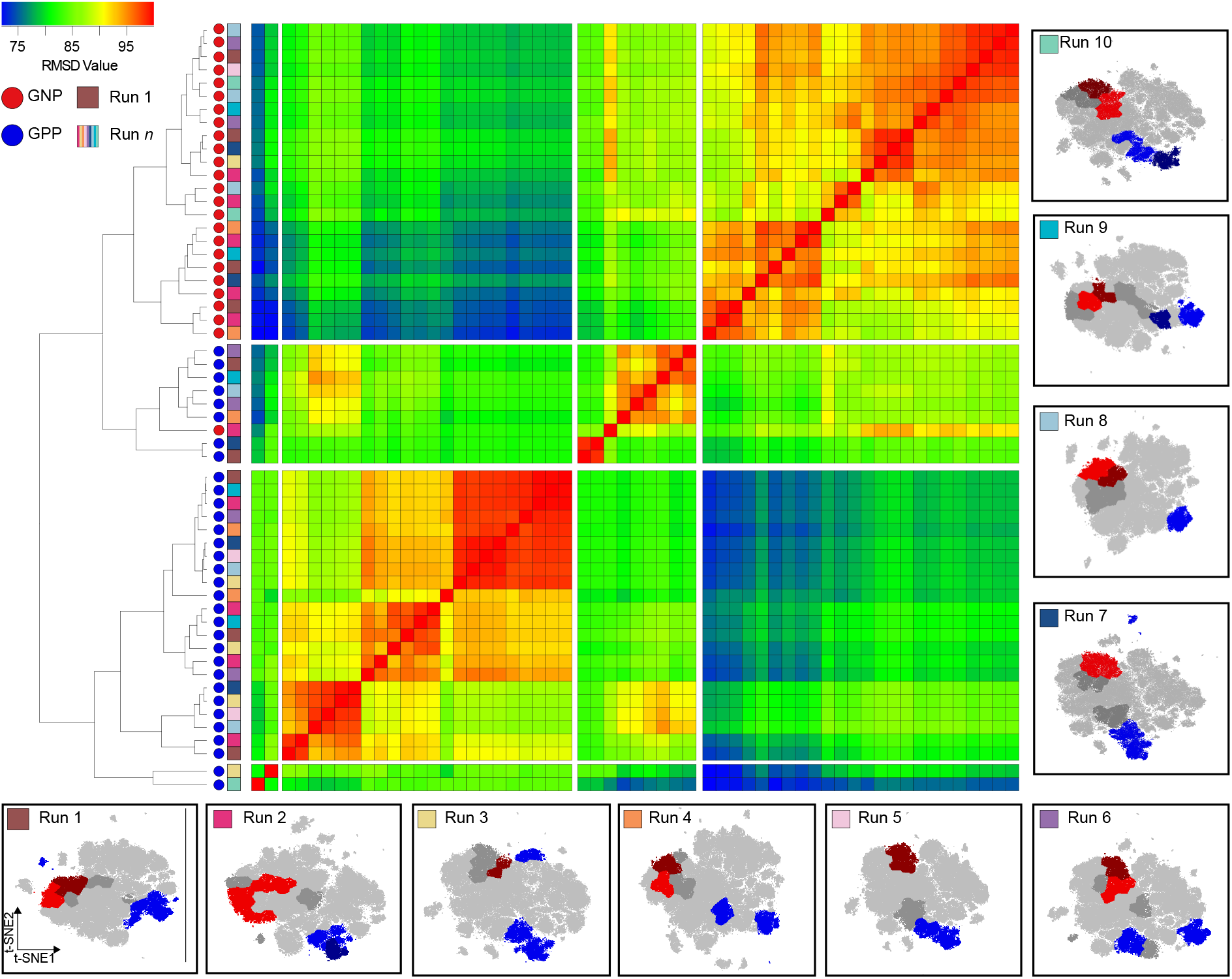
Subsampling of glioblastoma cells repeatedly resulted in GNP and GPP subsets with similar phenotypes. RMSD map comparing MEM scores for GNP and GPP subsets identified in the main figures and from nine additional t-SNE runs. GNP subsets are noted by red circles and GPP subsets are noted by blue circles. Colored boxes to the right of the red or blue circles indicate the t-SNE run from which the subset is derived. t-SNE runs are plotted around the heatmap with the corresponding colored box in the upper left of each plot.

Supplemental Information: **Patient specific view of population abundance and mass signal for all analyzed patients in this study**. Additional patient specific plots of the data are available in supplementary materials online. Each patient is shown on an individual page, with progression-free and overall survival data (also included in Supplementary Tables 1 and 3) shown at top right. At top left, the common t-SNE plot derived from analyzing an equal number of glioblastoma (non-immune, non-endothelial) cells from each of 28 patients (as in Figure 1), with contours indicating event abundance, is shown. Second from left, the density of events of the individual patient’s tumor is shown. Second from right, the assignment of cells from the patient to FlowSOM clusters is shown, and furthest right indicates the distribution of these clusters on the map of all GBM patients’ cells. FlowSOM clusters and the abundance of these clusters in the individual patient’s sample are shown in column at right side (also included in Supplementary Table 3). Clusters identified as “high” based on the criteria detailed in Methods are indicated in bold. Below, t-SNE plots and “heat” for each measured channel on cells from the individual patient are shown. (https://www.biorxiv.org/content/biorxiv/early/2019/05/10/632208/DC1/embed/media-1.pdf?download=true)

**Supplementary Table 1 –.**
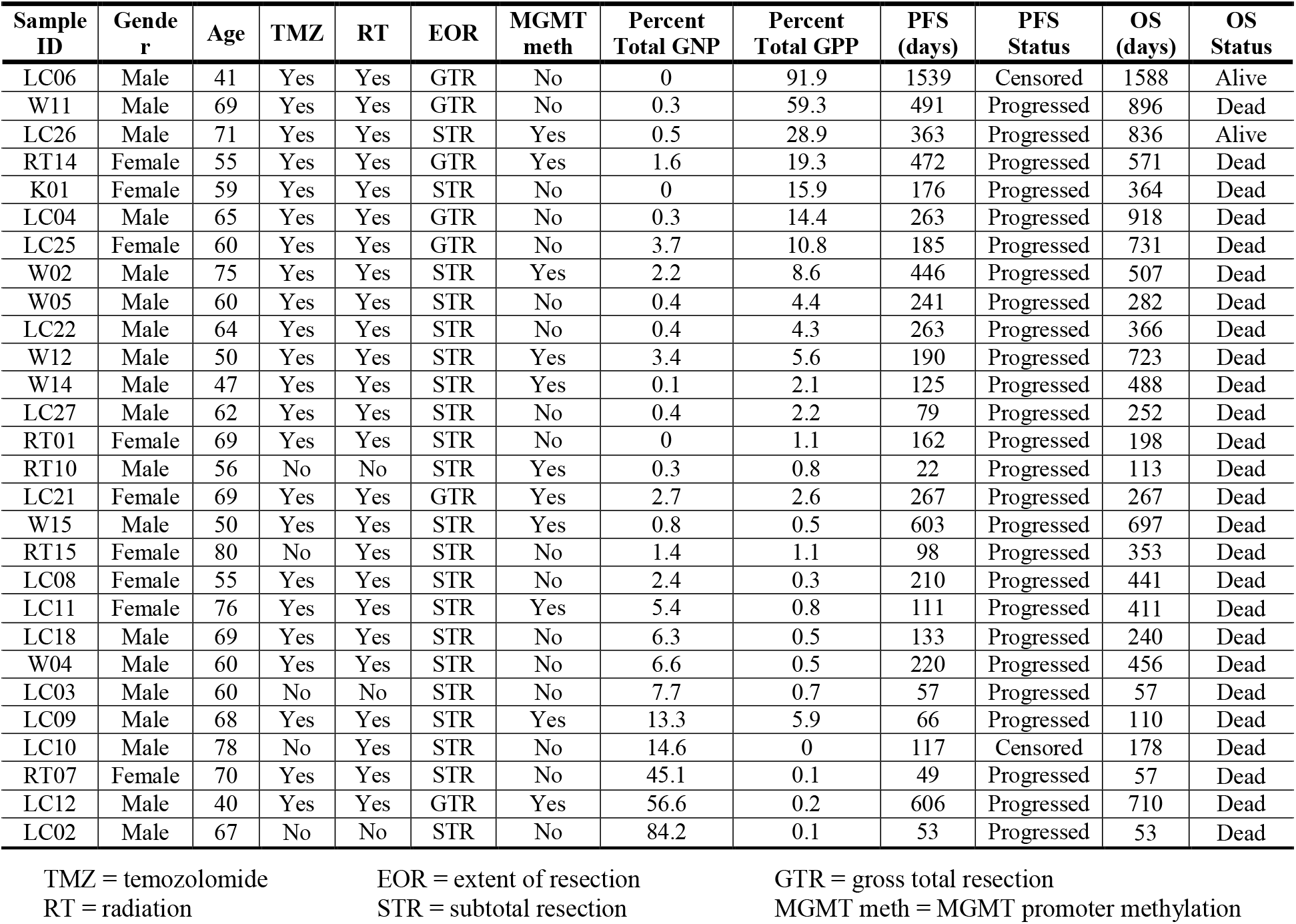
Glioblastoma Patient Characteristics

**Supplementary Table 2 –.** Mass cytometry antibody panels

| Target | Mass | Clone | Signaling & proteins |  | Stain |  |  |
| --- | --- | --- | --- | --- | --- | --- | --- |
|  |  |  | Panel | t-SNE | Live | Sap | MeOH |
| Rhodium | 103 | - | ● |  | ✓ |  |  |
| Cyclin B1 | 139 | GNS-1 | ● |  |  |  | ✓ |
| TUJ1 | 141 | TUJ1 | ● | ■ |  |  | ✓ |
| cCasp3 | 142 | 5A1E | ● |  |  |  | ✓ |
| CD117 | 143 | 104D2 | ● | ■ | ✓ |  |  |
| S100B | 144 | 19/S100B | ● | ■ |  |  | ✓ |
| CD31 | 145 | WM59 | ● | ■* | ✓ |  |  |
| γH2AX | 147 | JBW301 | ● |  |  |  | ✓ |
| CD34 | 148 | 581 | ● | ■ | ✓ |  |  |
| p-4E-BP1 (T37/T46) | 149 | 236B4 | ● |  |  |  | ✓ |
| p-STAT5 (Y694) | 150 | 47 | ● | ■ |  |  | ✓ |
| BMX | 151 | 40/BMX | ● |  |  |  | ✓ |
| p-AKT (S473) | 152 | D9E | ● | ■ |  |  | ✓ |
| p-STAT1 (Y701) | 153 | 58D6 | ● | ■ |  |  | ✓ |
| CD45 | 154 | HI30 | ● | ■* | ✓ |  |  |
| NCAM/CD56 | 155 | HCD56 | ● | ■ | ✓ |  |  |
| p-p38 (T180/Y182) | 156 | D3F9 | ● | ■ |  |  | ✓ |
| p-STAT3 (Y705) | 158 | 4/P-STAT3 | ● | ■ |  |  | ✓ |
| ITGα6/CD49F | 159 | GoH3 | ● | ■ | ✓ |  |  |
| CD133 | 160 | AC133 | ● | ■ | ✓ |  |  |
| PDGFRα | 161 | 16A1 | ● | ■ | ✓ |  |  |
| SOX2 | 163 | O30-678 | ● | ■ |  | ✓ |  |
| SSEA-1/CD15 | 164 | W6D3 | ● | ■ | ✓ |  |  |
| EGFR | 165 | AY13 | ● | ■ | ✓ |  |  |
| p-NFκB p65 (S529) | 166 | K10-895.12.50 | ● | ■ |  |  | ✓ |
| L1CAM | 167 | 5G3 | ● | ■ | ✓ |  |  |
| Nestin | 168 | 10C2 | ● | ■ |  |  | ✓ |
| CD44 | 169 | BJ18 | ● | ■ | ✓ |  |  |
| GFAP | 170 | 1B4 | ● | ■ |  |  | ✓ |
| p-ERK1/2 (T202/Y204) | 171 | D13.14.4E | ● | ■ |  |  | ✓ |
| p-S6 (S235/S236) | 172 | N7-548 | ● | ■ |  |  | ✓ |
| SOX10 | 173 | A-2 | ● |  |  |  | ✓ |
| HLA-DR | 174 | L243 | ● | ■ | ✓ |  |  |
| p-HH3 | 175 | HTA28 | ● |  |  |  | ✓ |
| Histone H3 | 176 | D1H2 | ● |  |  |  | ✓ |

● = included in the panel
■ = included for generation of t-SNE map
\* Excluded from t-SNE analyses of only glioblastoma cells
Live = live surface stain
Sap = 0.02% saponin stain
MeOH = stain after ice-cold methanol permeabilization

**Supplemental Table 3 -.**
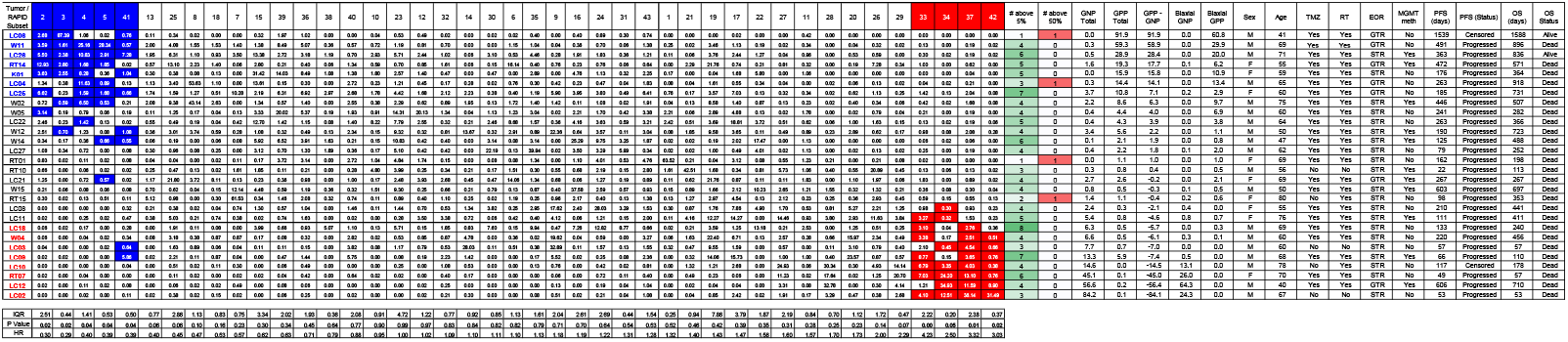
Subset and Patient Information

## References

1. Irish, J.M. & Doxie, D.B. High-dimensional single-cell cancer biology. Current topics in microbiology and immunology 377, 1–21 (2014).

2. Irish, J.M., Kotecha, N. & Nolan, G.P. Mapping normal and cancer cell signalling networks: towards single-cell proteomics. Nature reviews. Cancer 6, 146–155 (2006).

3. Irish, J.M. et al. Single cell profiling of potentiated phospho-protein networks in cancer cells. Cell 118, 217–228 (2004).

4. Levine, J.H. et al. Data-Driven Phenotypic Dissection of AML Reveals Progenitor-like Cells that Correlate with Prognosis. Cell 162, 184–197 (2015).

5. Irish, J.M. et al. B-cell signaling networks reveal a negative prognostic human lymphoma cell subset that emerges during tumor progression. Proceedings of the National Academy of Sciences of the United States of America 107, 12747–12754 (2010).

6. Good, Z. et al. Single-cell developmental classification of B cell precursor acute lymphoblastic leukemia at diagnosis reveals predictors of relapse. Nature medicine 24, 474–483 (2018).

7. Myklebust, J.H. et al. Distinct patterns of B-cell receptor signaling in non-Hodgkin lymphomas identified by single-cell profiling. Blood 129, 759–770 (2017).

8. Mistry, A.M., Greenplate, A.R., Ihrie, R.A. & Irish, J.M. Beyond the message: advantages of snapshot proteomics with single-cell mass cytometry in solid tumors. The FEBS journal 286, 1523–1539 (2019).

9. Spitzer, M.H. & Nolan, G.P. Mass Cytometry: Single Cells, Many Features. Cell 165, 780–791 (2016).

10. Irish, J.M. Beyond the age of cellular discovery. Nature immunology 15, 1095–1097 (2014).

11. Greenplate, A.R. et al. Computational Immune Monitoring Reveals Abnormal Double-Negative T Cells Present across Human Tumor Types. Cancer Immunol Res 7, 86–99 (2019).

12. Saeys, Y., Gassen, S.V. & Lambrecht, B.N. Computational flow cytometry: helping to make sense of high-dimensional immunology data. Nature reviews. Immunology 16, 449–462 (2016).

13. Diggins, K.E., Ferrell, P.B., Jr. & Irish, J.M. Methods for discovery and characterization of cell subsets in high dimensional mass cytometry data. Methods 82, 55–63 (2015).

14. Bruggner, R.V., Bodenmiller, B., Dill, D.L., Tibshirani, R.J. & Nolan, G.P. Automated identification of stratifying signatures in cellular subpopulations. Proceedings of the National Academy of Sciences of the United States of America 111, E2770–2777 (2014).

15. Arvaniti, E. & Claassen, M. Sensitive detection of rare disease-associated cell subsets via representation learning. Nature communications 8, 14825 (2017).

16. Gonzalez, V.D. et al. Commonly Occurring Cell Subsets in High-Grade Serous Ovarian Tumors Identified by Single-Cell Mass Cytometry. Cell Rep 22, 1875–1888 (2018).

17. Bendall, S.C. et al. Single-cell mass cytometry of differential immune and drug responses across a human hematopoietic continuum. Science 332, 687–696 (2011).

18. Amir el, A.D. et al. viSNE enables visualization of high dimensional single-cell data and reveals phenotypic heterogeneity of leukemia. Nat Biotechnol 31, 545–552 (2013).

19. Becht, E. et al. Dimensionality reduction for visualizing single-cell data using UMAP. Nat Biotechnol (2018).

20. Van Gassen, S. et al. FlowSOM: Using self-organizing maps for visualization and interpretation of cytometry data. Cytometry. Part A: the journal of the International Society for Analytical Cytology (2015).

21. Diggins, K.E., Greenplate, A.R., Leelatian, N., Wogsland, C.E. & Irish, J.M. Characterizing cell subsets using marker enrichment modeling. Nature methods 14, 275–278 (2017).

22. Newell, E.W. & Cheng, Y. Mass cytometry: blessed with the curse of dimensionality. Nature immunology 17, 890–895 (2016).

23. Gandelman, J.S. et al. Machine learning reveals chronic graft-versus-host disease phenotypes and stratifies survival after stem cell transplant for hematologic malignancies. Haematologica (2018).

24. Leelatian, N. et al. Single cell analysis of human tissues and solid tumors with mass cytometry. Cytometry B Clin Cytom 92, 68–78 (2017).

25. Doxie, D.B. et al. BRAF and MEK inhibitor therapy eliminates Nestin-expressing melanoma cells in human tumors. Pigment cell & melanoma research 31, 708–719 (2018).

26. Ostrom, Q.T. et al. CBTRUS Statistical Report: Primary brain and other central nervous system tumors diagnosed in the United States in 2010-2014. Neuro Oncol 19, v1–v88 (2017).

27. Stupp, R. et al. Radiotherapy plus concomitant and adjuvant temozolomide for glioblastoma. N Engl J Med 352, 987–996 (2005).

28. Gilbert, M.R. et al. A randomized trial of bevacizumab for newly diagnosed glioblastoma. N Engl J Med 370, 699–708 (2014).

29. Patel, A.P. et al. Single-cell RNA-seq highlights intratumoral heterogeneity in primary glioblastoma. Science 344, 1396–1401 (2014).

30. Wei, W. et al. Single-Cell Phosphoproteomics Resolves Adaptive Signaling Dynamics and Informs Targeted Combination Therapy in Glioblastoma. Cancer Cell 29, 563–573 (2016).

31. Brennan, C.W. et al. The somatic genomic landscape of glioblastoma. Cell 155, 462–477 (2013).

32. Verhaak, R.G. et al. Integrated genomic analysis identifies clinically relevant subtypes of glioblastoma characterized by abnormalities in PDGFRA, IDH1, EGFR, and NF1. Cancer Cell 17, 98–110 (2010).

33. Brennan, C. et al. Glioblastoma subclasses can be defined by activity among signal transduction pathways and associated genomic alterations. PLoS One 4, e7752 (2009).

34. Johnson, H. & White, F.M. Quantitative analysis of signaling networks across differentially embedded tumors highlights interpatient heterogeneity in human glioblastoma. J Proteome Res 13, 4581–4593 (2014).

35. Stommel, J.M. et al. Coactivation of receptor tyrosine kinases affects the response of tumor cells to targeted therapies. Science 318, 287–290 (2007).

36. Snuderl, M. et al. Mosaic amplification of multiple receptor tyrosine kinase genes in glioblastoma. Cancer cell 20, 810–817 (2011).

37. Meyer, M. et al. Single cell-derived clonal analysis of human glioblastoma links functional and genomic heterogeneity. Proc Natl Acad Sci U S A 112, 851–856 (2015).

38. Leelatian, N. et al. Preparing Viable Single Cells from Human Tissue and Tumors for Cytomic Analysis. Curr Protoc Mol Biol 118, 25C 21 21–25C 21 23 (2017).

39. Cancer Genome Atlas Research, N. et al. Comprehensive, Integrative Genomic Analysis of Diffuse Lower-Grade Gliomas. N Engl J Med 372, 2481–2498 (2015).

40. Singh, S.K. et al. Identification of human brain tumour initiating cells. Nature 432, 396–401 (2004).

41. Leelatian, N., Diggins, K.E. & Irish, J.M. Characterizing Phenotypes and Signaling Networks of Single Human Cells by Mass Cytometry. Methods in molecular biology 1346, 99–113 (2015).

42. Van Gassen, S. et al. FlowSOM: Using self-organizing maps for visualization and interpretation of cytometry data. Cytometry A 87, 636–645 (2015).

43. Ohgaki, H. et al. Genetic pathways to glioblastoma: a population-based study. Cancer Res 64, 6892–6899 (2004).

44. Shapiro, W.R. et al. Randomized trial of three chemotherapy regimens and two radiotherapy regimens and two radiotherapy regimens in postoperative treatment of malignant glioma. Brain Tumor Cooperative Group Trial 8001. J Neurosurg 71, 1–9 (1989).

45. Hegi, M.E. et al. MGMT gene silencing and benefit from temozolomide in glioblastoma. N Engl J Med 352, 997–1003 (2005).

46. Brown, C.E. et al. Regression of Glioblastoma after Chimeric Antigen Receptor T-Cell Therapy. N Engl J Med 375, 2561–2569 (2016).

47. Brown, T.J. et al. Association of the Extent of Resection With Survival in Glioblastoma: A Systematic Review and Meta-analysis. JAMA Oncol 2, 1460–1469 (2016).

48. Grabowski, M.M. et al. Residual tumor volume versus extent of resection: predictors of survival after surgery for glioblastoma. J Neurosurg 121, 1115–1123 (2014).

49. Mirimanoff, R.O. et al. Radiotherapy and temozolomide for newly diagnosed glioblastoma: recursive partitioning analysis of the EORTC 26981/22981-NCIC CE3 phase III randomized trial. J Clin Oncol 24, 2563–2569 (2006).

50. Walker, M.D. et al. Randomized comparisons of radiotherapy and nitrosoureas for the treatment of malignant glioma after surgery. N Engl J Med 303, 1323–1329 (1980).

51. Hussain, S.F. et al. The role of human glioma-infiltrating microglia/macrophages in mediating antitumor immune responses. Neuro Oncol 8, 261–279 (2006).

52. Behbehani, G.K. et al. Transient partial permeabilization with saponin enables cellular barcoding prior to surface marker staining. Cytometry. Part A: the journal of the International Society for Analytical Cytology 85, 1011–1019 (2014).

53. Finck, R. et al. Normalization of mass cytometry data with bead standards. Cytometry. Part A: the journal of the International Society for Analytical Cytology 83, 483–494 (2013).

54. Beyrend, G., Stam, K., Hollt, T., Ossendorp, F. & Arens, R. Cytofast: A workflow for visual and quantitative analysis of flow and mass cytometry data to discover immune signatures and correlations. Comput Struct Biotechnol J 16, 435–442 (2018).

55. Lan, X. et al. Fate mapping of human glioblastoma reveals an invariant stem cell hierarchy. Nature 549, 227–232 (2017).

56. Wei, J. et al. miR-124 inhibits STAT3 signaling to enhance T cell-mediated immune clearance of glioma. Cancer Res 73, 3913–3926 (2013).

57. Dolma, S. et al. Inhibition of Dopamine Receptor D4 Impedes Autophagic Flux, Proliferation, and Survival of Glioblastoma Stem Cells. Cancer Cell 29, 859–873 (2016).

58. Fan, Q. et al. A Kinase Inhibitor Targeted to mTORC1 Drives Regression in Glioblastoma. Cancer Cell 31, 424–435 (2017).

59. Fan, Q.W. et al. EGFR phosphorylates tumor-derived EGFRvIII driving STAT3/5 and progression in glioblastoma. Cancer Cell 24, 438–449 (2013).

60. Fan, Q.W. et al. EGFR signals to mTOR through PKC and independently of Akt in glioma. Sci Signal 2, ra4 (2009).

61. Fujioka, S. et al. Stabilization of p53 is a novel mechanism for proapoptotic function of NF-kappaB. J Biol Chem 279, 27549–27559 (2004).

62. Schneider, A. et al. NF-kappaB is activated and promotes cell death in focal cerebral ischemia. Nat Med 5, 554–559 (1999).

63. Venteicher, A.S. et al. Decoupling genetics, lineages, and microenvironment in IDH-mutant gliomas by single-cell RNA-seq. Science 355 (2017).

64. Wang, Q. et al. Tumor Evolution of Glioma-Intrinsic Gene Expression Subtypes Associates with Immunological Changes in the Microenvironment. Cancer Cell 32, 42–56 e46 (2017).

65. Mistry, A.M. et al. Decreased survival in glioblastomas is specific to contact with the ventricular-subventricular zone, not subgranular zone or corpus callosum. J Neurooncol 132, 341–349 (2017).

66. Kotecha, N., Krutzik, P.O. & Irish, J.M. Web-based analysis and publication of flow cytometry experiments. Current protocols in cytometry / editorial board, J. Paul Robinson, managing editor … [et al.] Chapter 10, Unit10 17 (2010).

