## Supplemental Single Cell Patient Information for "High risk glioblastoma cells revealed by machine learning and single cell signaling profiles"

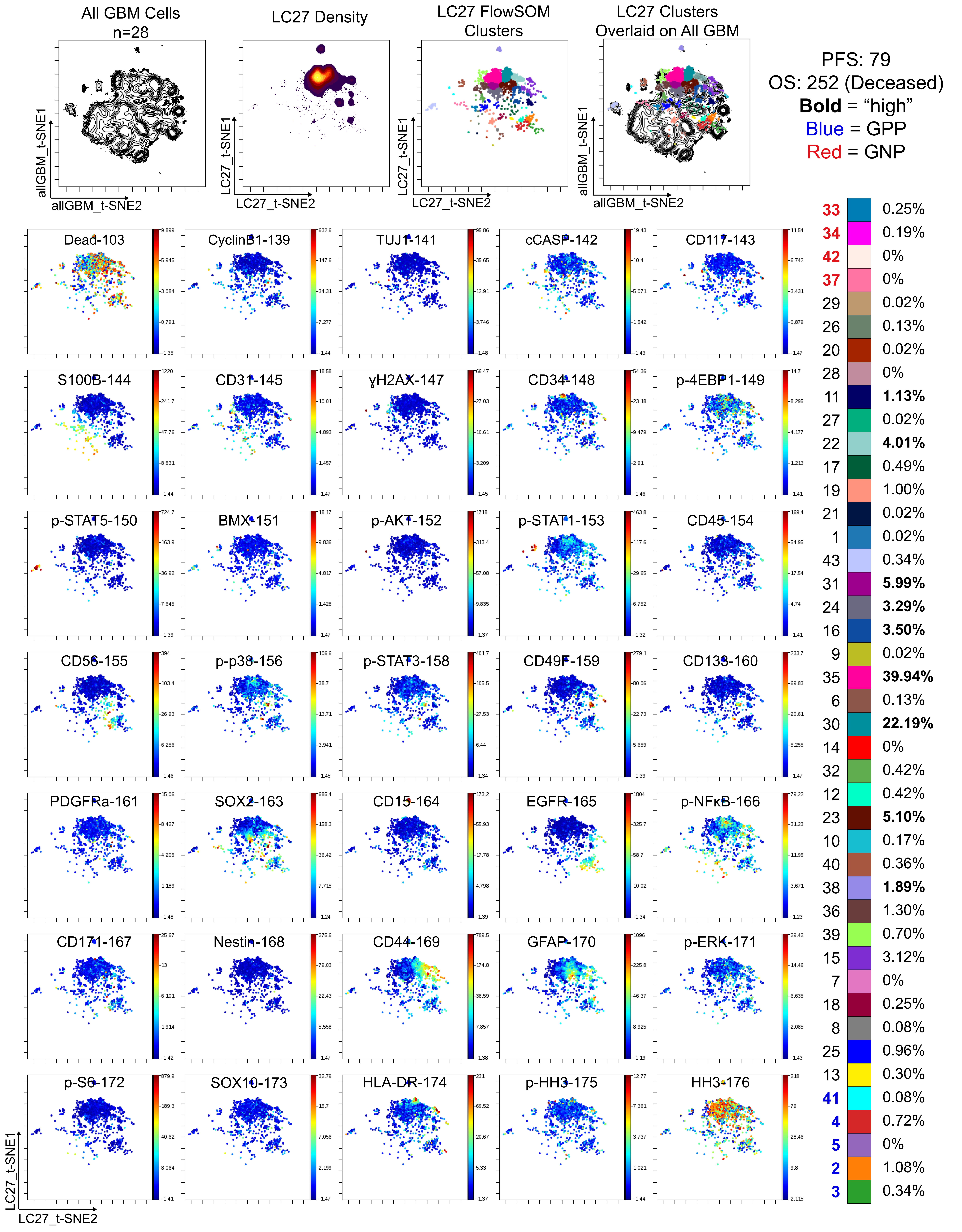

Supplemental Single Cell Patient Information

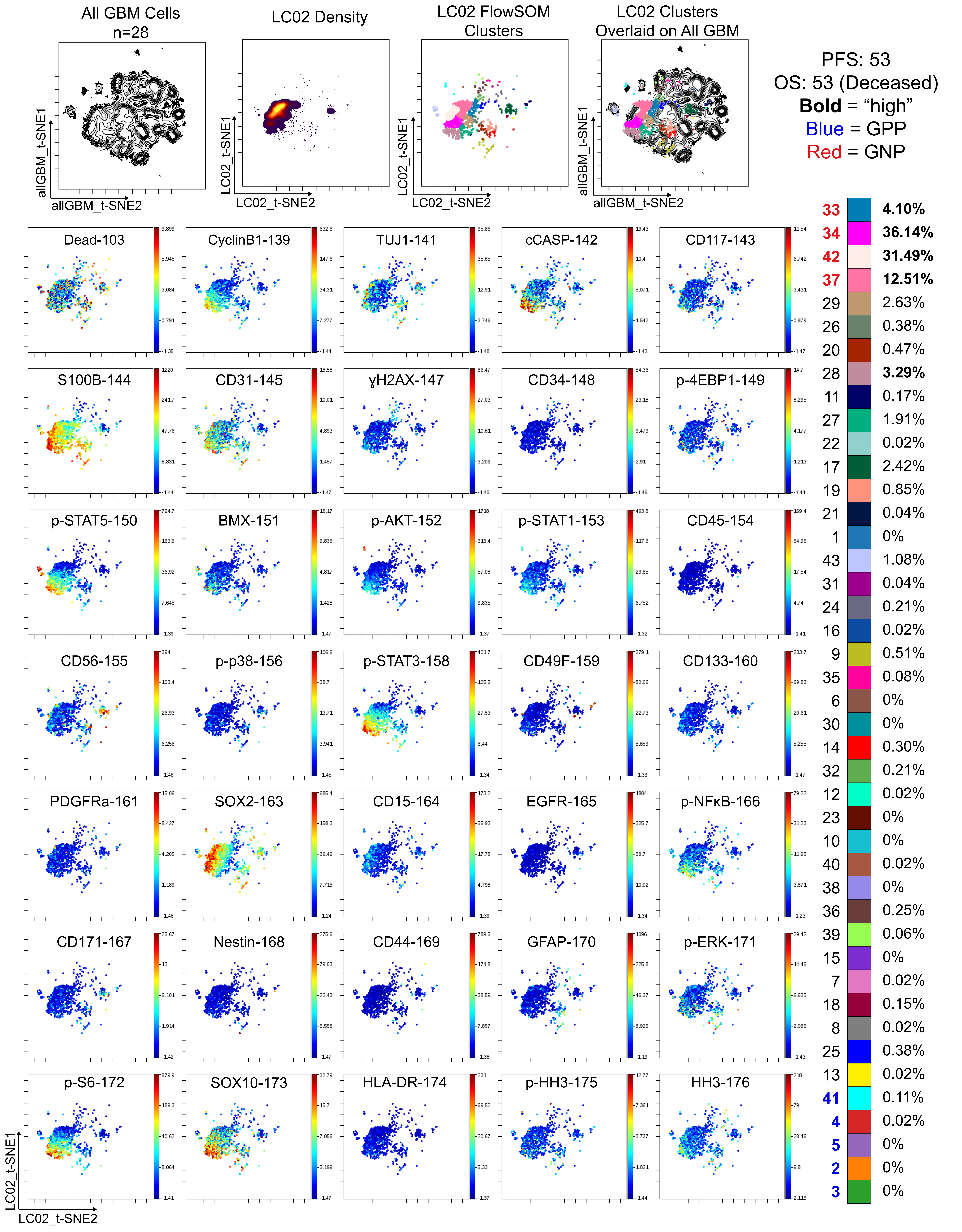

Supplemental Single Cell Patient Information

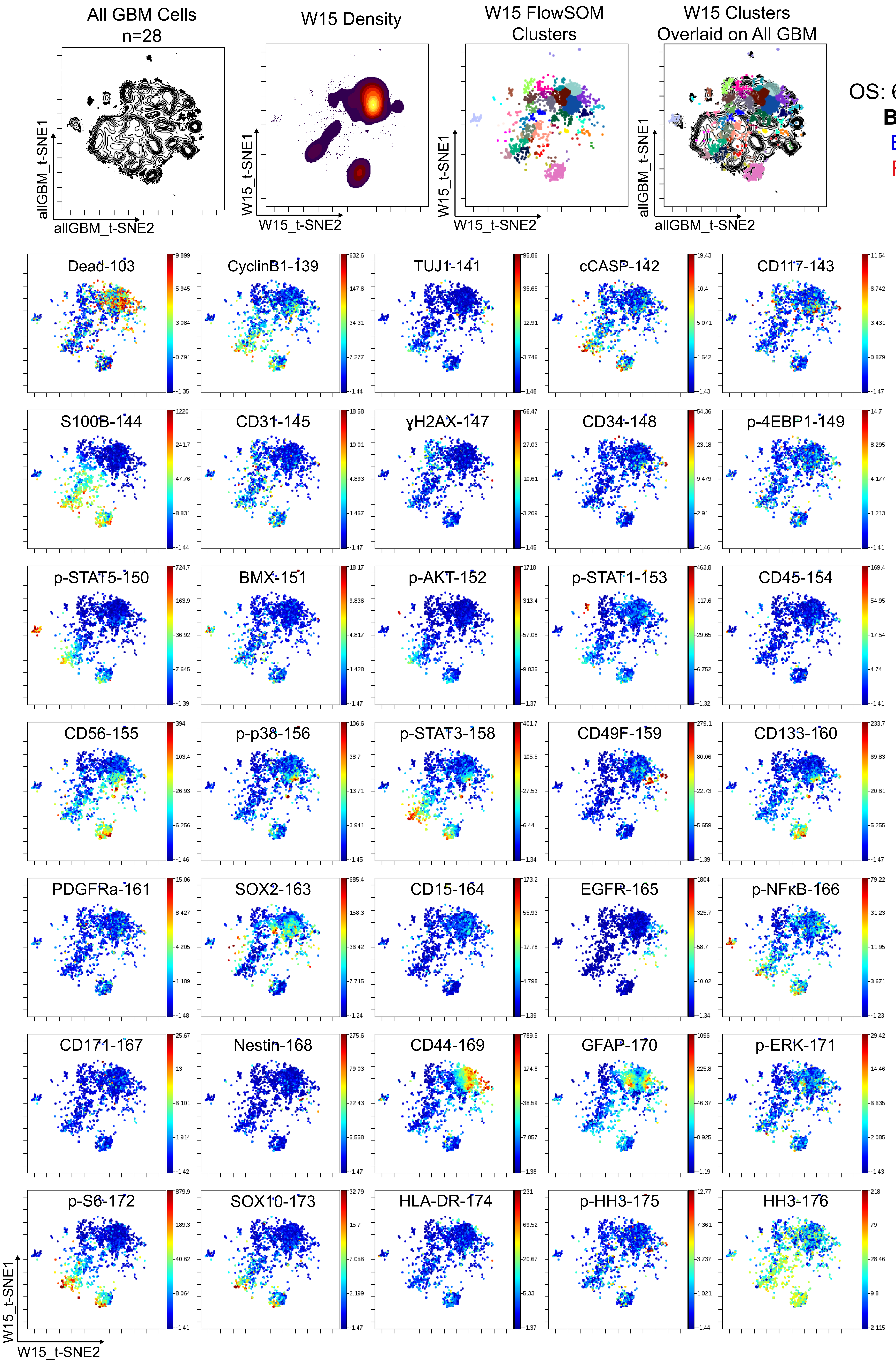


PFS: 603

OS: 697 (Deceased)

**Bold** = "high"

Blue = GPP

Red = GNP

| Cluster | Percentage |
|---|---|
| **33** | 0.36% |
| **34** | 0.30% |
| **42** | 0.04% |
| **37** | 0.08% |
| 29 | 0.21% |
| 26 | 1.32% |
| 20 | 0.32% |
| 28 | **1.55%** |
| 11 | **1.21%** |
| 27 | **2.65%** |
| 22 | **10.23%** |
| 17 | 2.12% |
| 19 | 1.66% |
| 21 | 0.89% |
| 1 | 0.15% |
| 43 | 0.93% |
| 31 | **0.57%** |
| 24 | 2.59% |
| 16 | **37.58%** |
| 9 | 0.40% |
| 35 | 0.87% |
| 6 | 0.13% |
| 30 | 0.79% |
| 14 | 0.21% |
| 32 | 0.66% |
| 12 | 0.25% |
| 23 | **9.30%** |
| 10 | **1.51%** |
| 40 | 0.32% |
| 38 | **0.36%** |
| 36 | 1.19% |
| 39 | 0.59% |
| 15 | **4.48%** |
| 7 | **12.14%** |
| 18 | 0.15% |
| 8 | 0.04% |
| 25 | 0.62% |
| 13 | 0.70% |
| **41** | 0.08% |
| **4** | 0.08% |
| **5** | 0.06% |
| **2** | 0.21% |
| **3** | 0.06% |

Supplemental Single Cell Patient Information

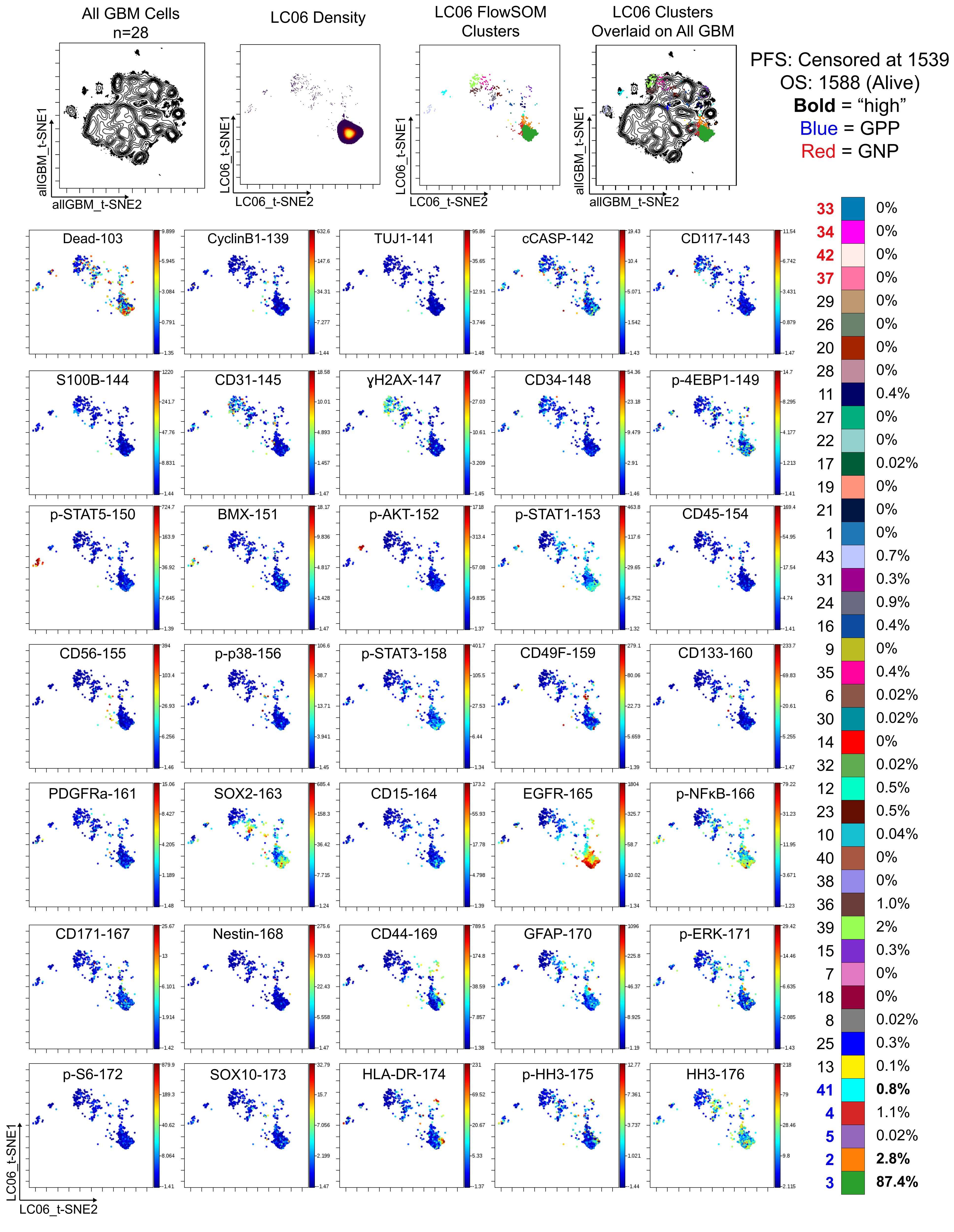

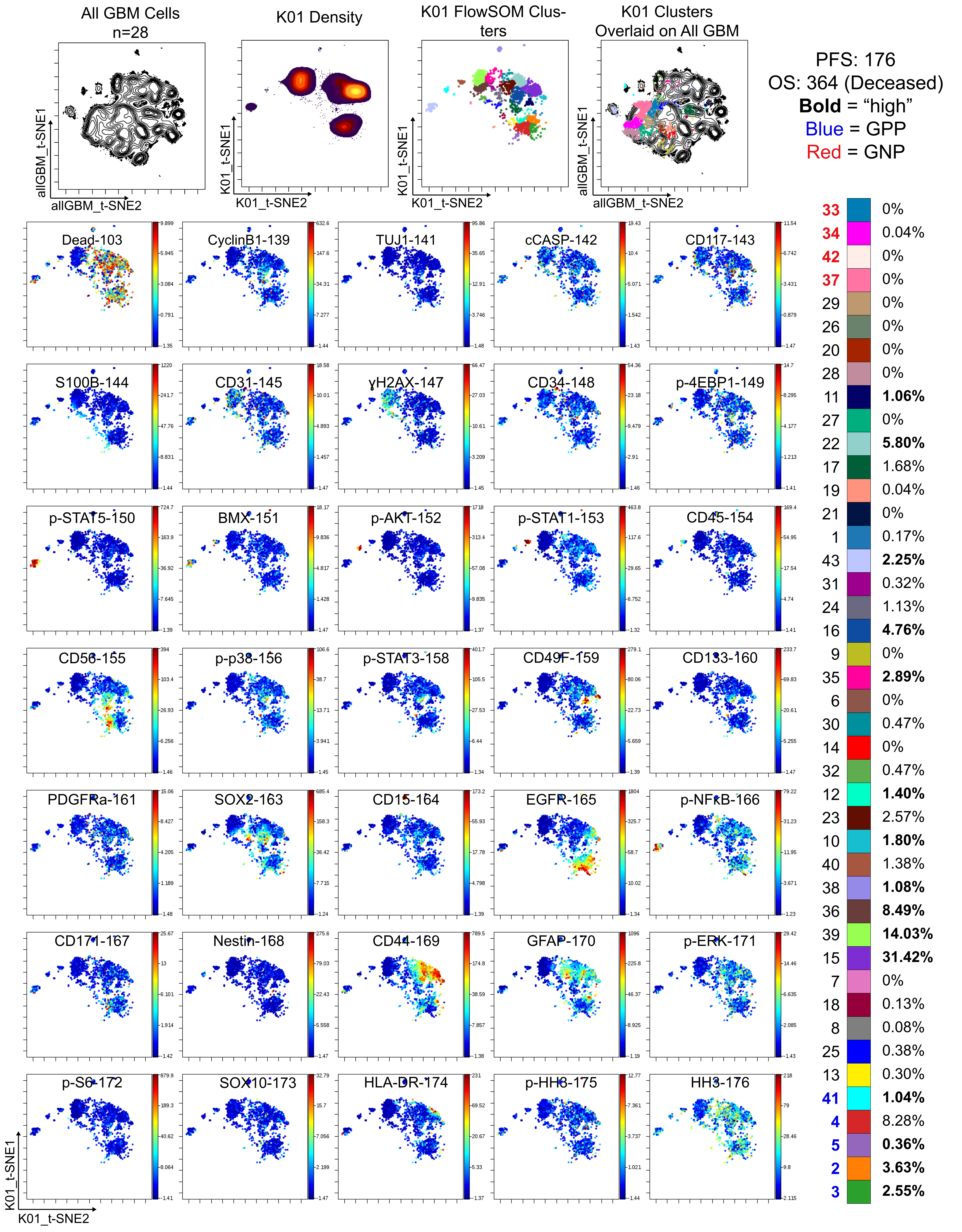
All GBM Cells
n=28
K01 Density
K01 FlowSOM Clusters
K01 Clusters Overlaid on All GBM
allGBM_t-SNE1
allGBM_t-SNE2
K01_t-SNE1
K01_t-SNE2
PFS: 176
OS: 364 (Deceased)
Bold = "high"
Blue = GPP
Red = GNP
Dead-103
CyclinB1-139
TUJ1-141
cCASP-142
CD117-143
S100B-144
CD31-145
γH2AX-147
CD34-148
p-4EBP1-149
p-STAT5-150
BMX-151
p-AKT-152
p-STAT1-153
CD45-154
CD56-155
p-p38-156
p-STAT3-158
CD49F-159
CD133-160
PDGFRa-161
SOX2-163
CD15-164
EGFR-165
p-NFkB-166
CD171-167
Nestin-168
CD44-169
GFAP-170
p-ERK-171
p-S6-172
SOX10-173
HLA-DR-174
p-HH3-175
HH3-176
33 0%
34 0.04%
42 0%
37 0%
29 0%
26 0%
20 0%
28 0%
11 1.06%
27 0%
22 5.80%
17 1.68%
19 0.04%
21 0%
1 0.17%
43 2.25%
31 0.32%
24 1.13%
16 4.76%
9 0%
35 2.89%
6 0%
30 0.47%
14 0%
32 0.47%
12 1.40%
23 2.57%
10 1.80%
40 1.38%
38 1.08%
36 8.49%
39 14.03%
15 31.42%
7 0%
18 0.13%
8 0.08%
25 0.38%
13 0.30%
41 1.04%
4 8.28%
5 0.36%
2 3.63%
3 2.55%

Supplemental Single Cell Patient Information

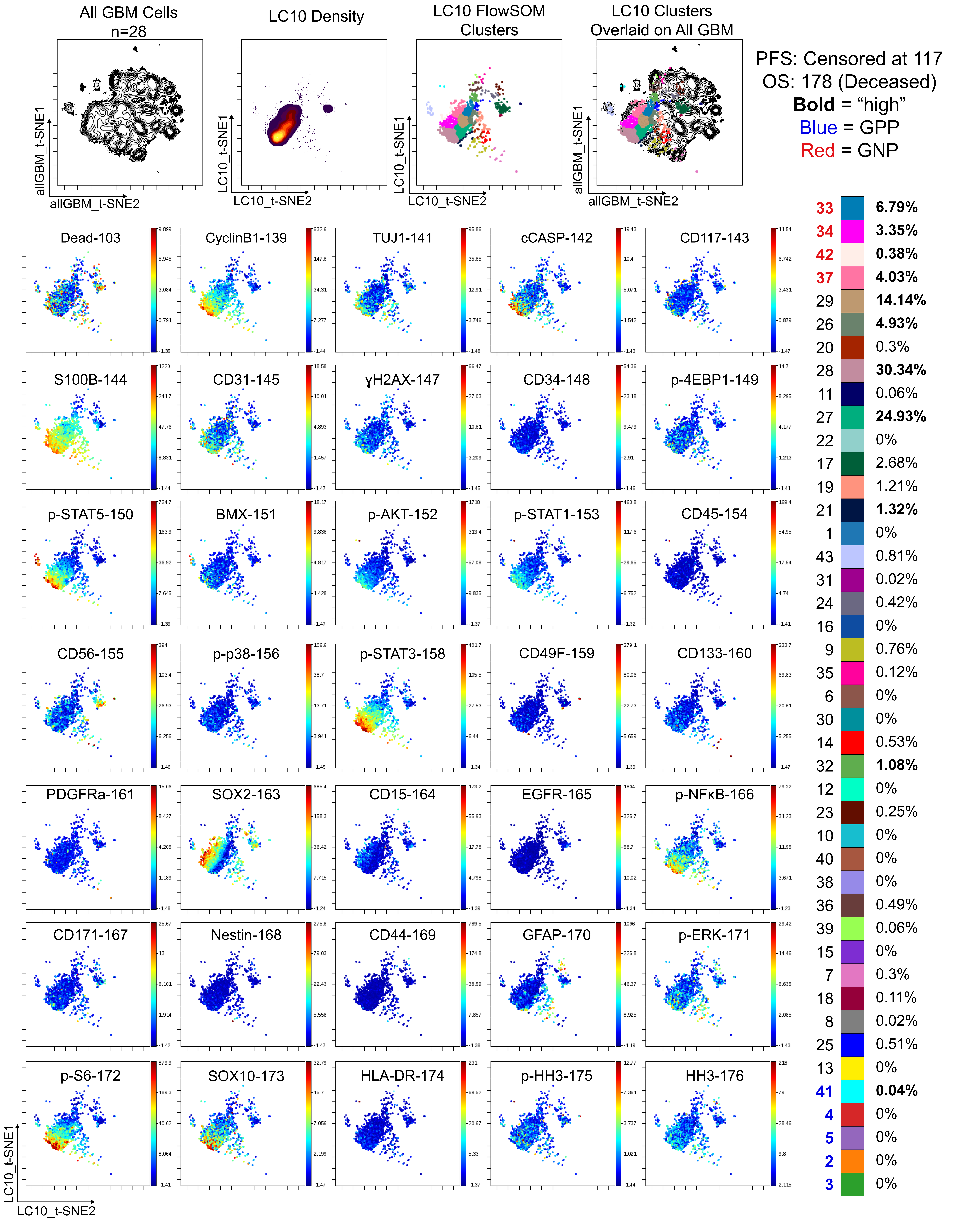

Supplemental Single Cell Patient Information

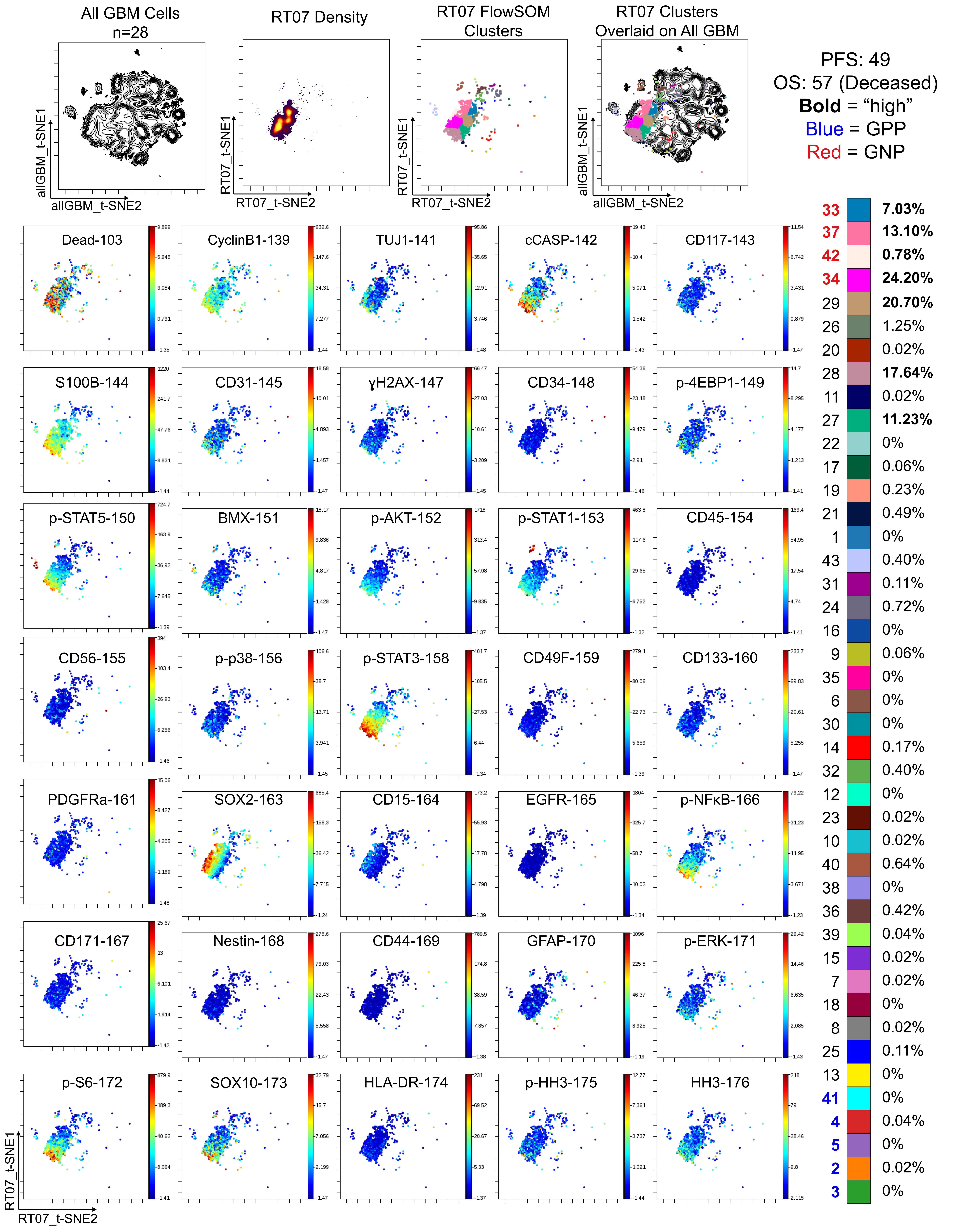

Supplemental Single Cell Patient Information

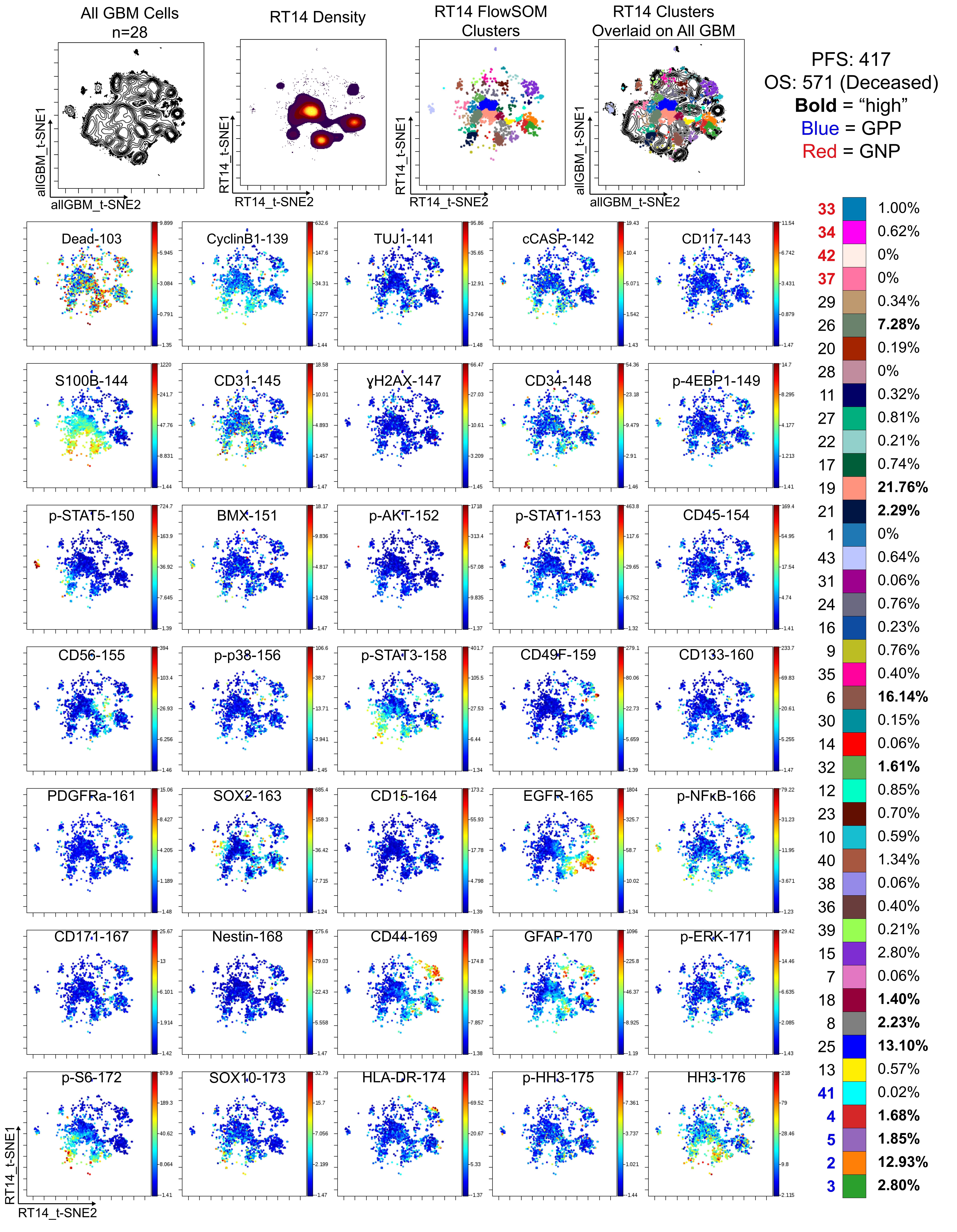

### Supplemental Single Cell Patient Information

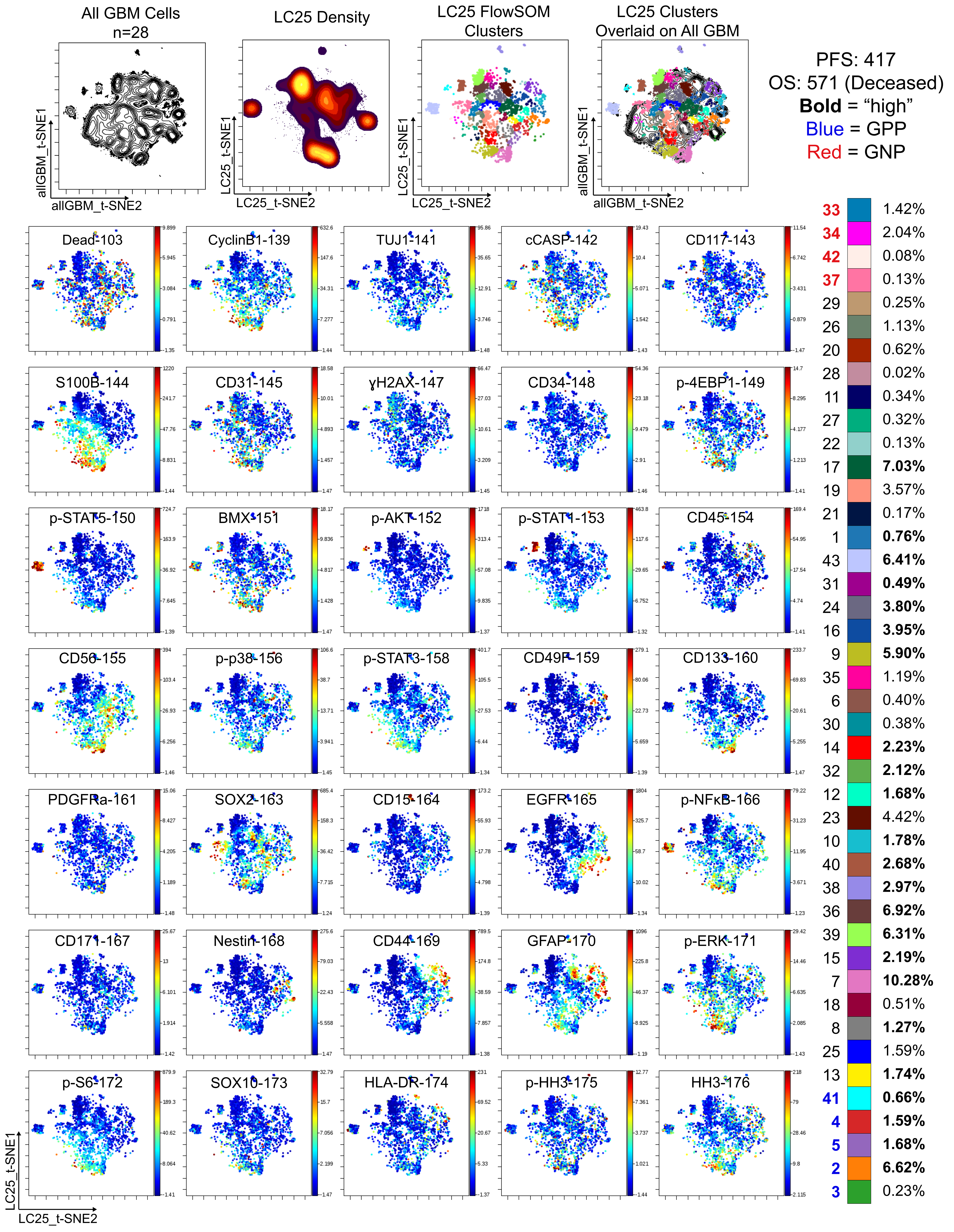

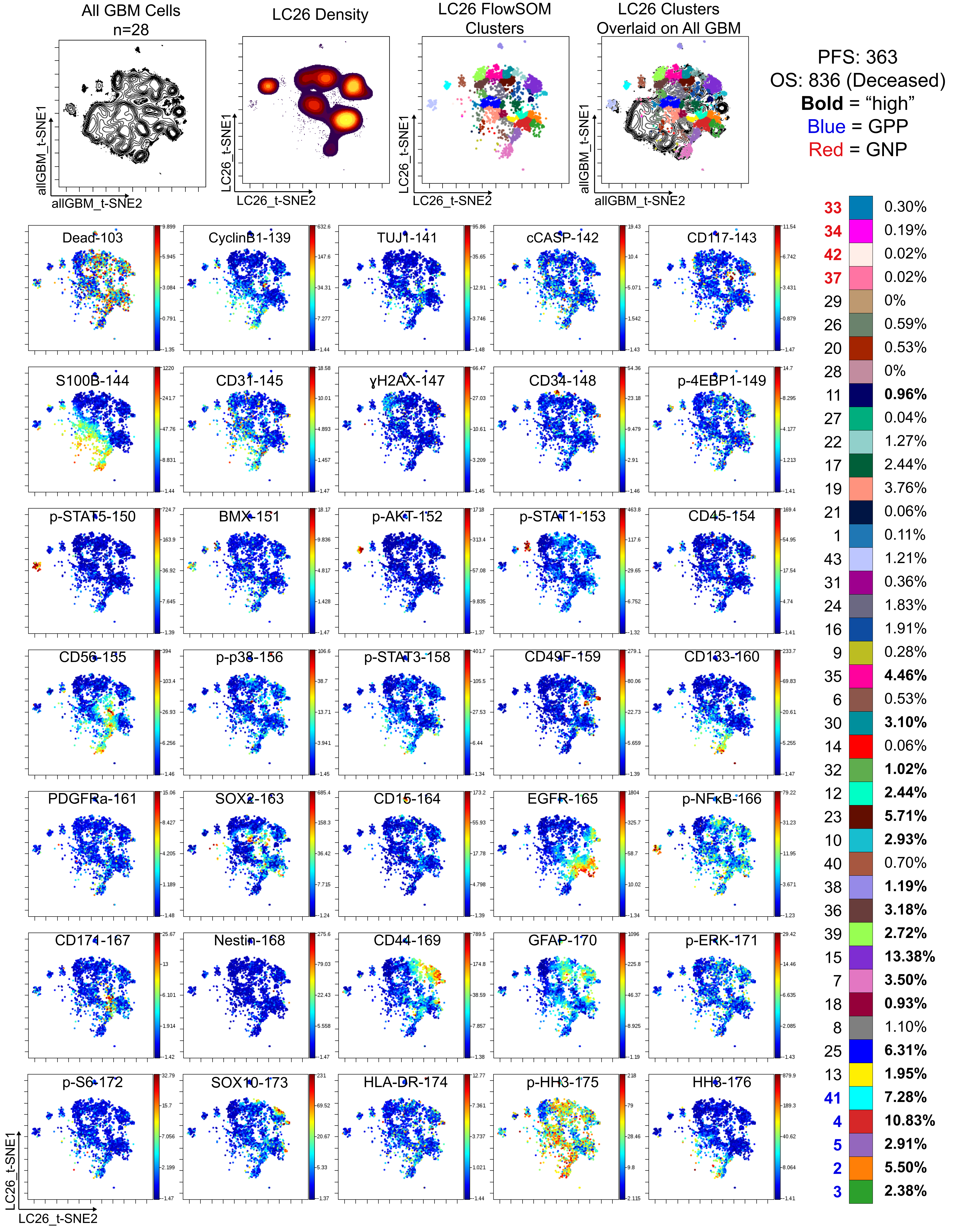
All GBM Cells
n=28
LC26 Density
LC26 FlowSOM Clusters
LC26 Clusters Overlaid on All GBM
allGBM_t-SNE1
allGBM_t-SNE2
LC26_t-SNE1
LC26_t-SNE2
PFS: 363
OS: 836 (Deceased)
Bold = "high"
Blue = GPP
Red = GNP
33 0.30%
34 0.19%
42 0.02%
37 0.02%
29 0%
26 0.59%
20 0.53%
28 0%
11 0.96%
27 0.04%
22 1.27%
17 2.44%
19 3.76%
21 0.06%
1 0.11%
43 1.21%
31 0.36%
24 1.83%
16 1.91%
9 0.28%
35 4.46%
6 0.53%
30 3.10%
14 0.06%
32 1.02%
12 2.44%
23 5.71%
10 2.93%
40 0.70%
38 1.19%
36 3.18%
39 2.72%
15 13.38%
7 3.50%
18 0.93%
8 1.10%
25 6.31%
13 1.95%
41 7.28%
4 10.83%
5 2.91%
2 5.50%
3 2.38%
Dead-103
CyclinB1-139
TUJ1-141
cCASP-142
CD117-143
S100B-144
CD31-145
γH2AX-147
CD34-148
p-4EBP1-149
p-STAT5-150
BMX-151
p-AKT-152
p-STAT1-153
CD45-154
CD56-155
p-p38-156
p-STAT3-158
CD49F-159
CD133-160
PDGFRa-161
SOX2-163
CD15-164
EGFR-165
p-NFκB-166
CD171-167
Nestin-168
CD44-169
GFAP-170
p-ERK-171
p-S6-172
SOX10-173
HLA-DR-174
p-HH3-175
HH3-176

Supplemental Single Cell Patient Information

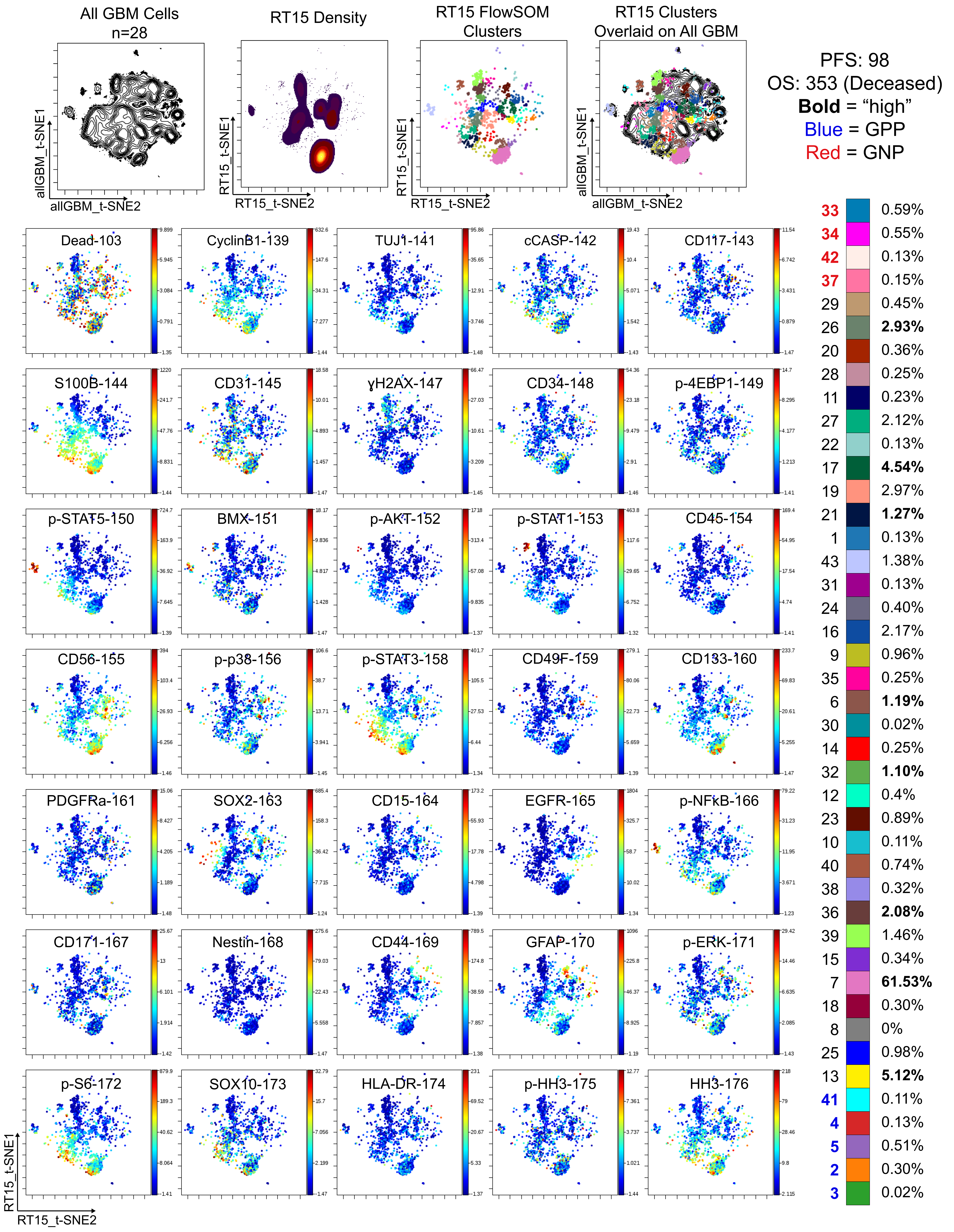

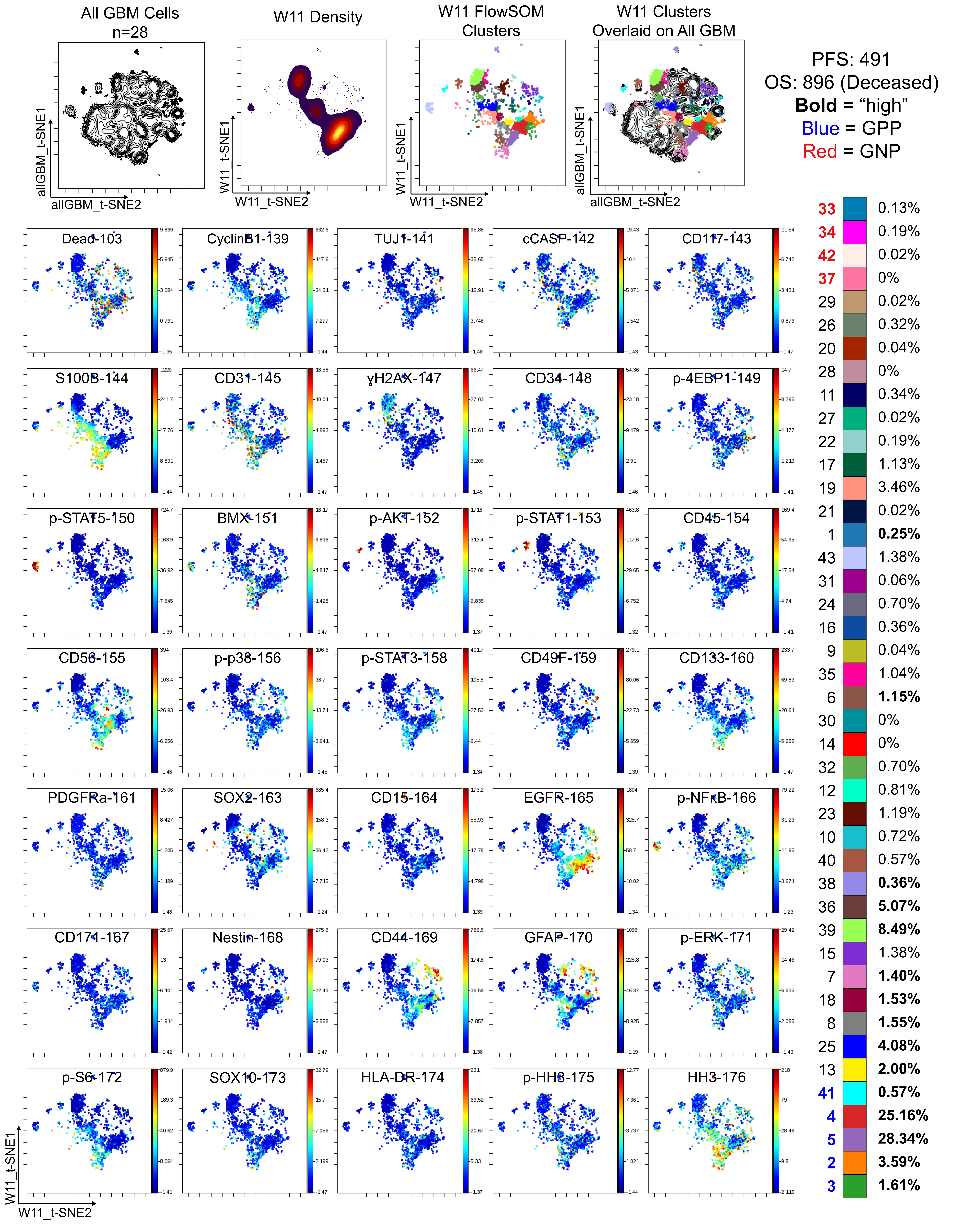
All GBM Cells
n=28
W11 Density
W11 FlowSOM Clusters
W11 Clusters Overlaid on All GBM
allGBM_t-SNE1
allGBM_t-SNE2
W11_t-SNE1
W11_t-SNE2
PFS: 491
OS: 896 (Deceased)
Bold = “high”
Blue = GPP
Red = GNP
33 0.13%
34 0.19%
42 0.02%
37 0%
29 0.02%
26 0.32%
20 0.04%
28 0%
11 0.34%
27 0.02%
22 0.19%
17 1.13%
19 3.46%
21 0.02%
1 0.25%
43 1.38%
31 0.06%
24 0.70%
16 0.36%
9 0.04%
35 1.04%
6 1.15%
30 0%
14 0%
32 0.70%
12 0.81%
23 1.19%
10 0.72%
40 0.57%
38 0.36%
36 5.07%
39 8.49%
15 1.38%
7 1.40%
18 1.53%
8 1.55%
25 4.08%
13 2.00%
41 0.57%
4 25.16%
5 28.34%
2 3.59%
3 1.61%
Dead-103
CyclinB1-139
TUJ1-141
cCASP-142
CD117-143
S100B-144
CD31-145
γH2AX-147
CD34-148
p-4EBP1-149
p-STAT5-150
BMX-151
p-AKT-152
p-STAT1-153
CD45-154
CD56-155
p-p38-156
p-STAT3-158
CD49F-159
CD133-160
PDGFRa-161
SOX2-163
CD15-164
EGFR-165
p-NFkB-166
CD171-167
Nestin-168
CD44-169
GFAP-170
p-ERK-171
p-S6-172
SOX10-173
HLA-DR-174
p-HH3-175
HH3-176

Supplemental Single Cell Patient Information

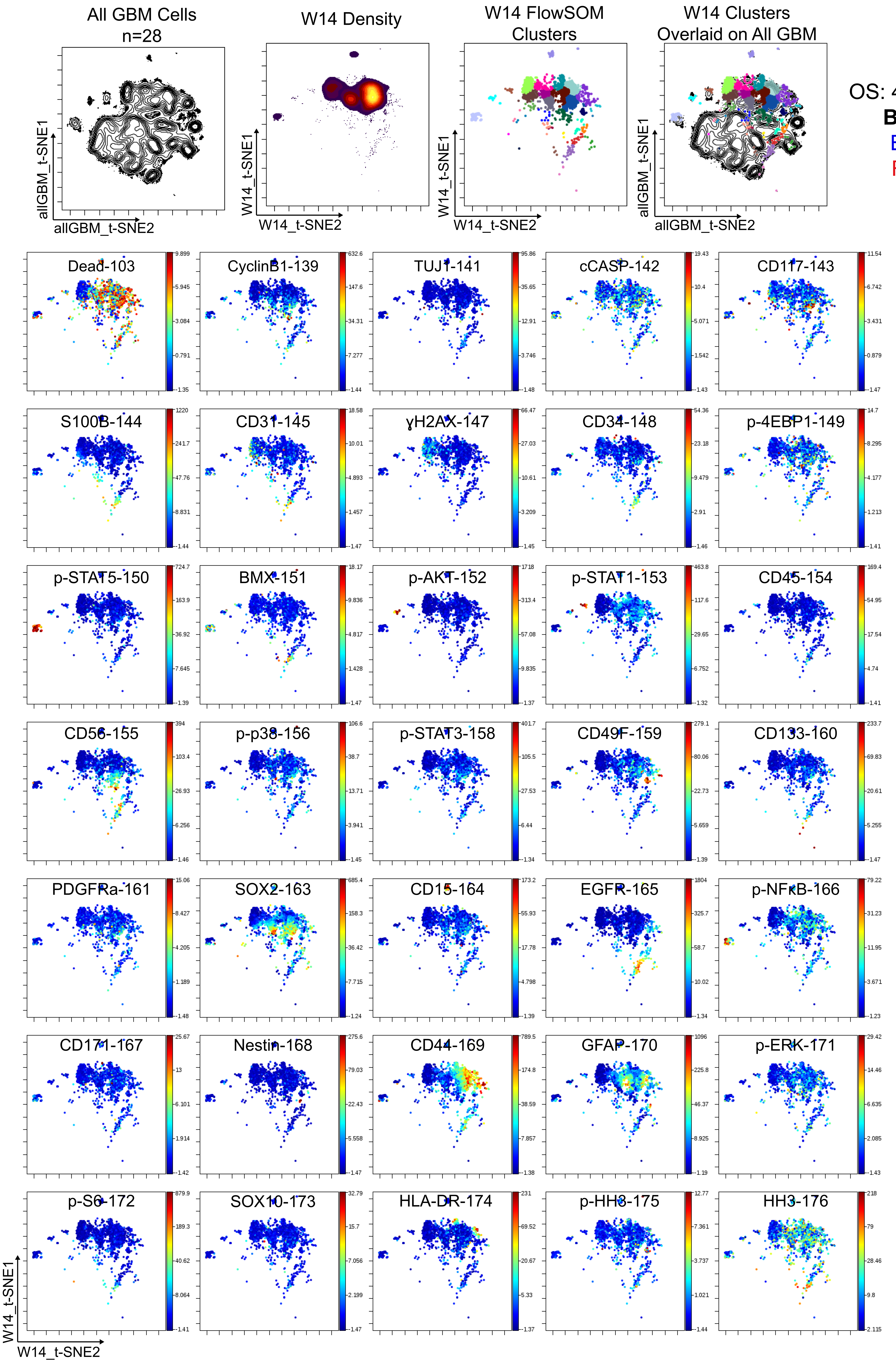


PFS: 125
OS: 488 (Deceased)
**Bold** = "high"
Blue = GPP
Red = GNP

| | |
|---|---|
| **33** | 0.06% |
| **34** | 0.06% |
| **42** | 0% |
| **37** | 0.02% |
| 29 | 0% |
| 26 | 0% |
| 20 | 0% |
| 28 | 0% |
| 11 | **1.13%** |
| 27 | 0% |
| 22 | **17.47%** |
| 17 | 2.02% |
| 19 | 0.19% |
| 21 | 0.02% |
| 1 | 0.02% |
| 43 | **1.87%** |
| 31 | **3.25%** |
| 24 | **9.75%** |
| 16 | **25.29%** |
| 9 | 0% |
| 35 | **3.14%** |
| 6 | 0.08% |
| 30 | **3.14%** |
| 14 | 0% |
| 32 | 0.40% |
| 12 | 0.42% |
| 23 | **10.83%** |
| 10 | 0.15% |
| 40 | 0.21% |
| 38 | **1.63%** |
| 36 | **3.91%** |
| 39 | **6.52%** |
| 15 | **5.92%** |
| 7 | 0.08% |
| 18 | 0.06% |
| 8 | 0% |
| 25 | 0.19% |
| 13 | 0.06% |
| **41** | **0.55%** |
| **4** | 0.36% |
| **5** | **0.66%** |
| **2** | 0.34% |
| **3** | 0.17% |

Supplemental Single Cell Patient Information

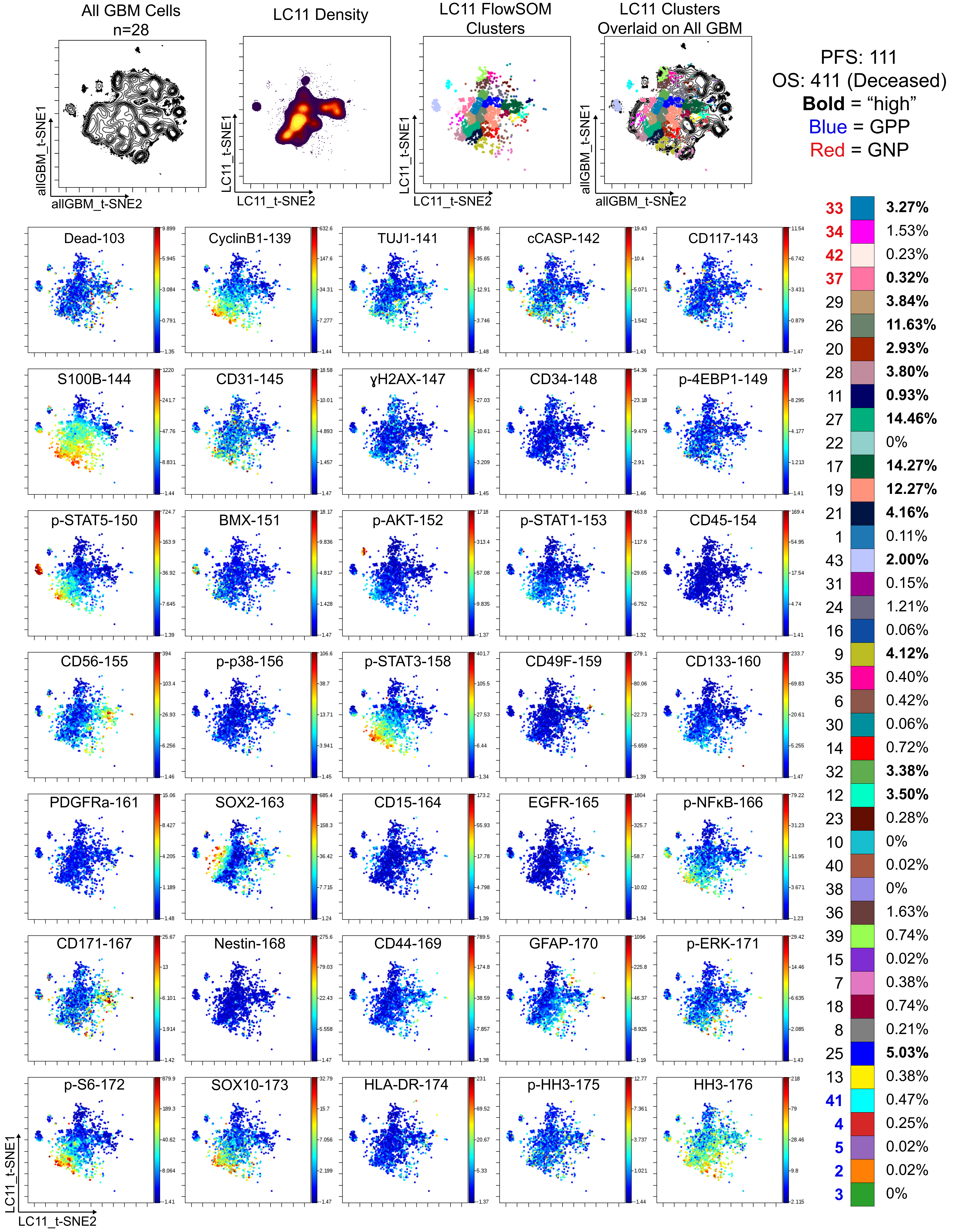

Supplemental Single Cell Patient Information

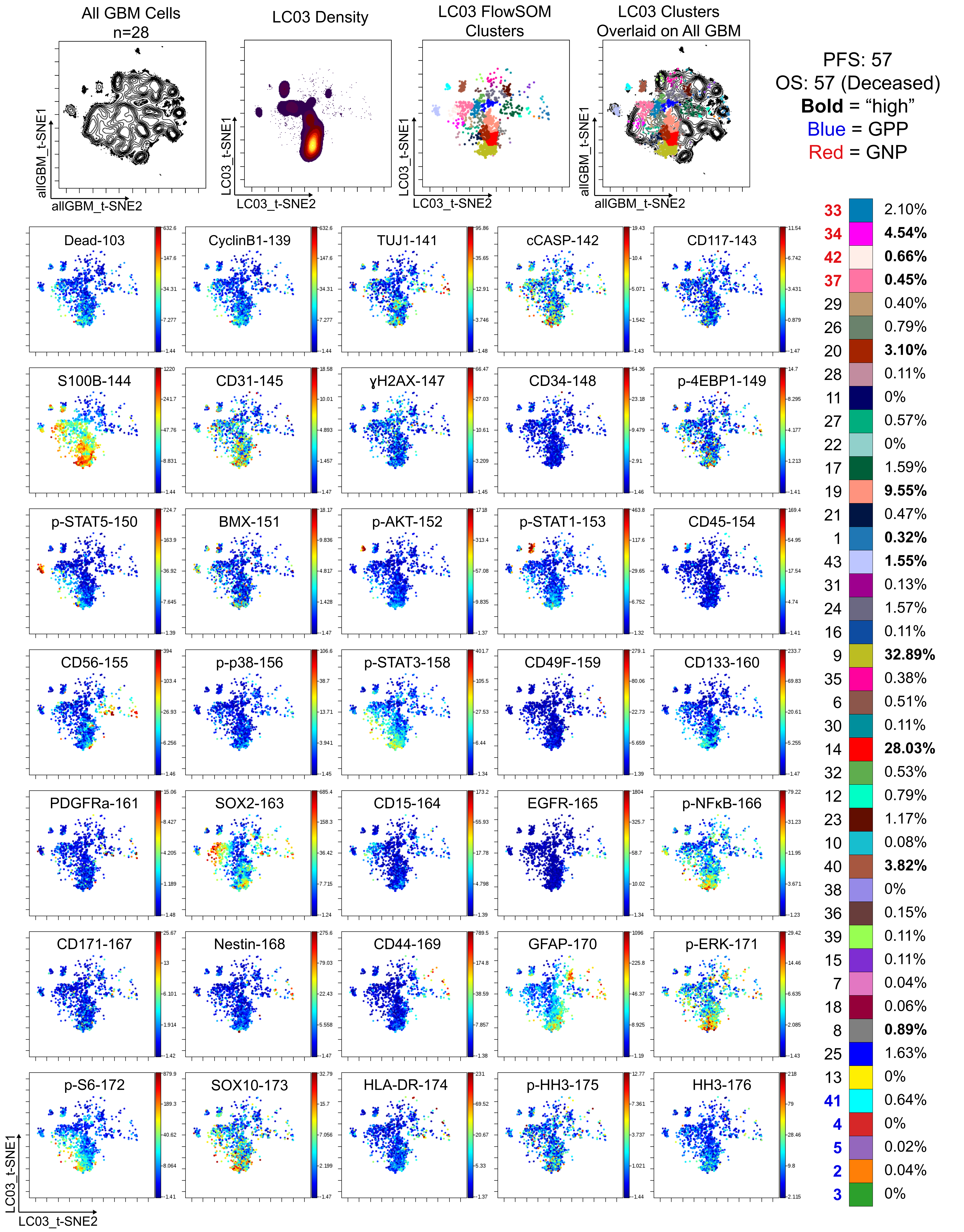

Supplemental Single Cell Patient Information

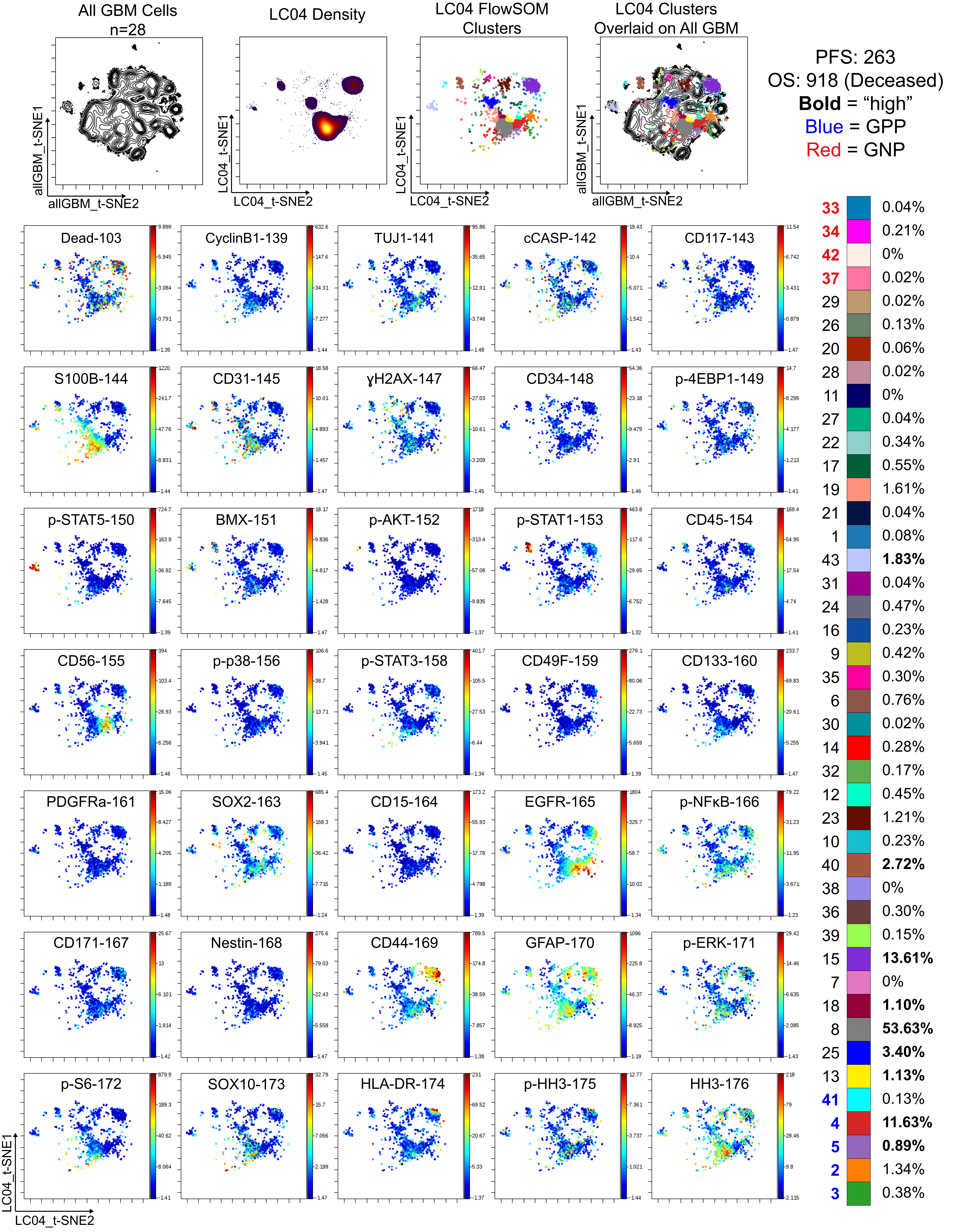

Supplemental Single Cell Patient Information

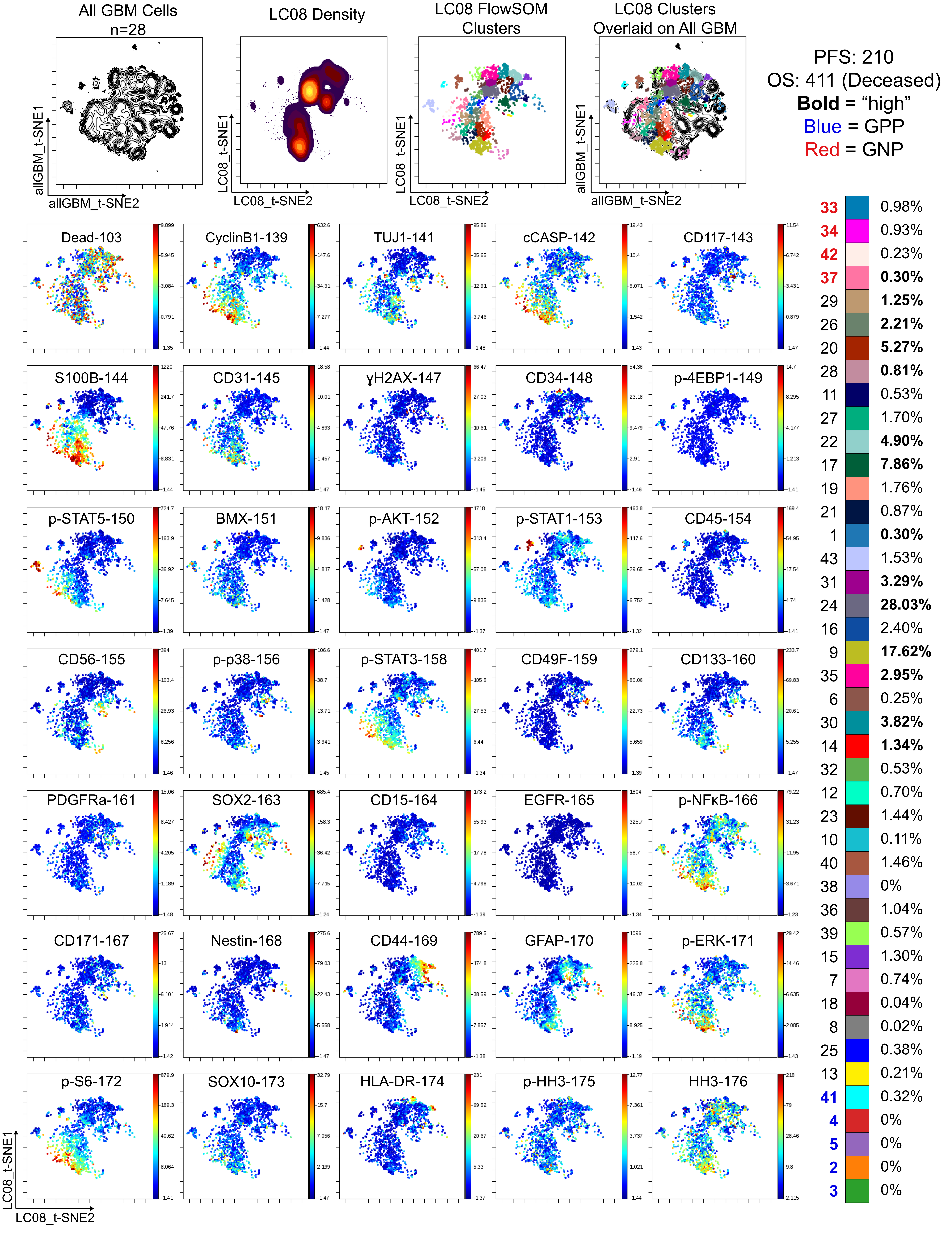

Supplemental Single Cell Patient Information

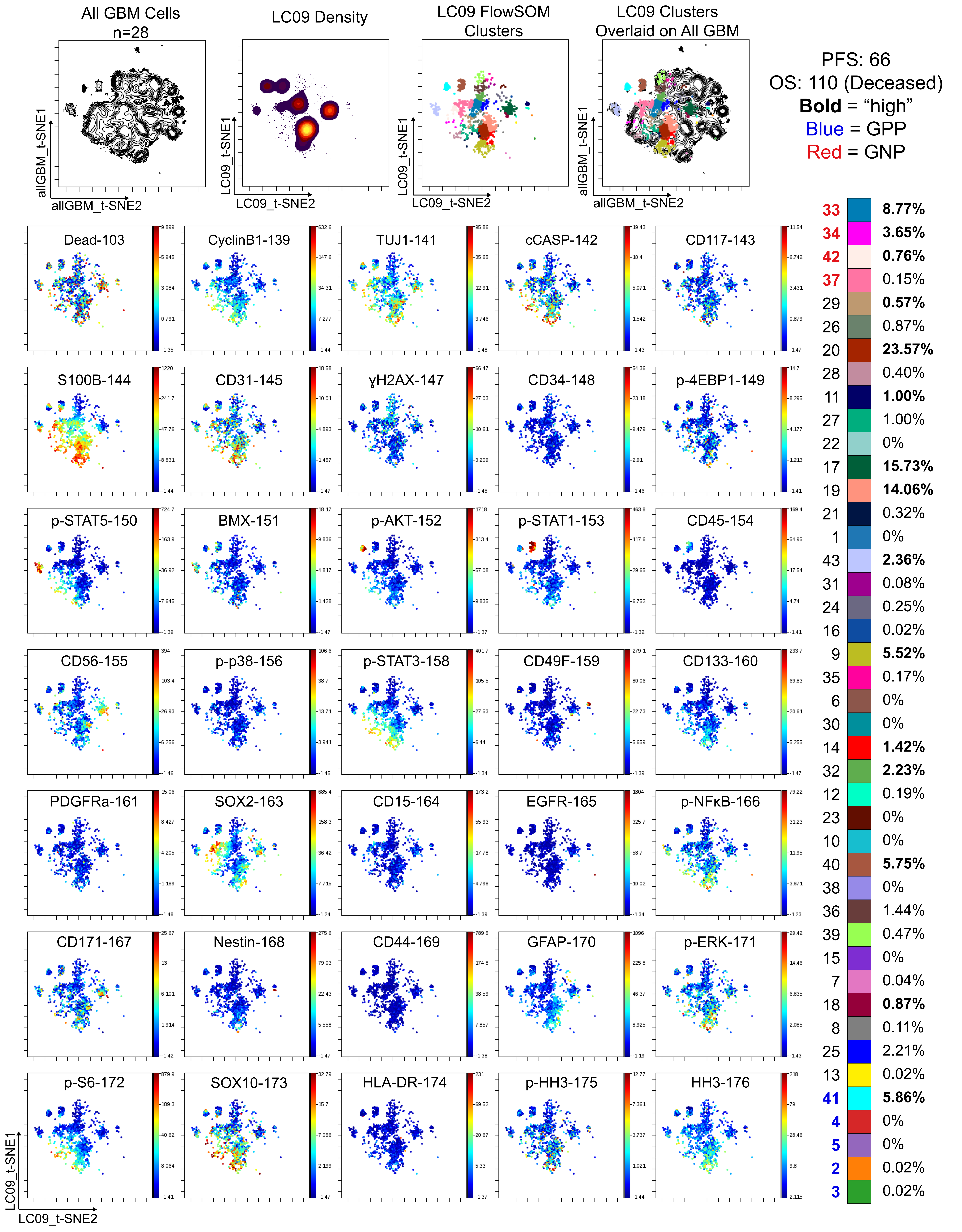

Supplemental Single Cell Patient Information

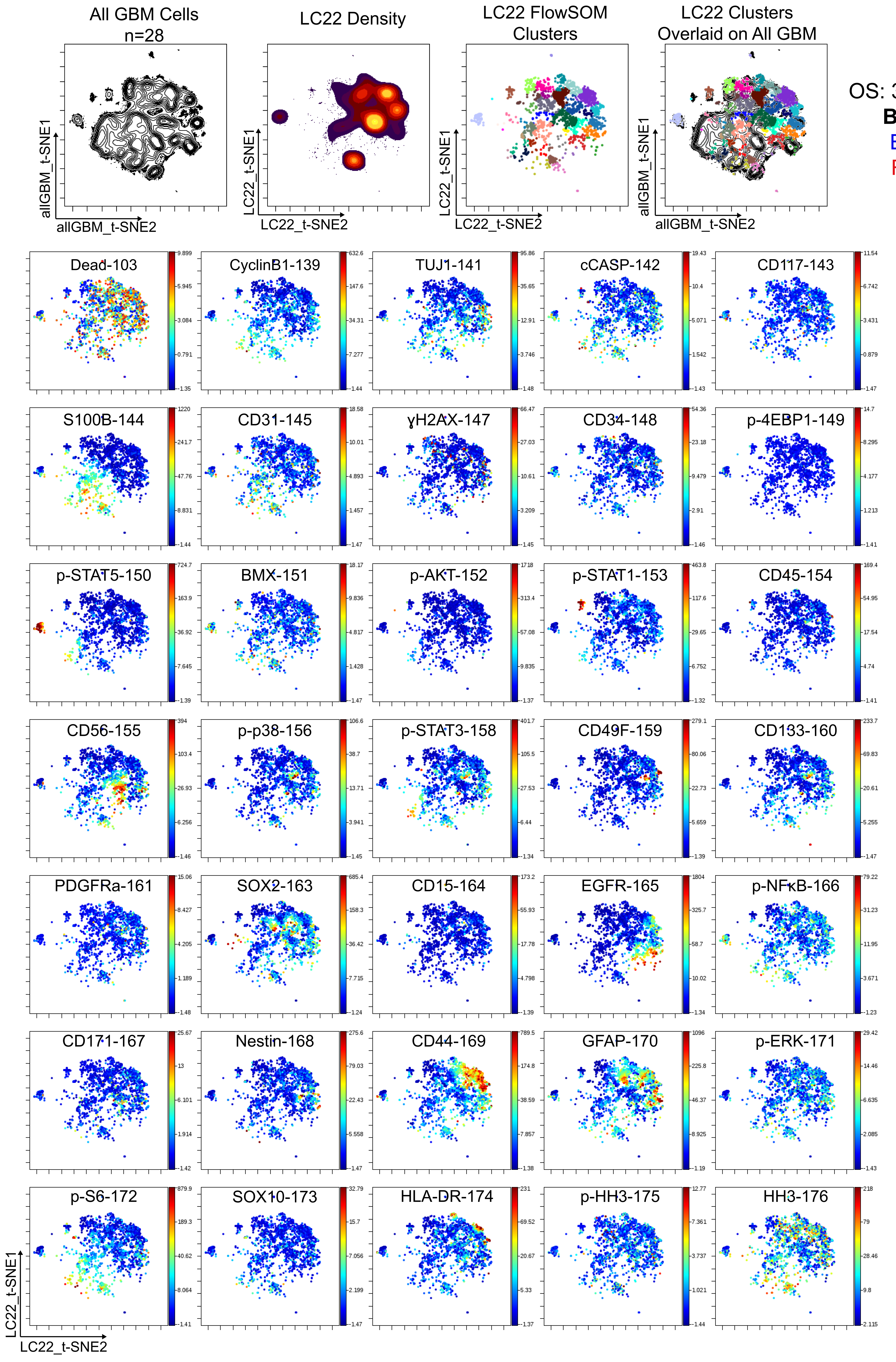


PFS: 263
OS: 366 (Deceased)
**Bold** = "high"
Blue = GPP
Red = GNP

| | |
|---|---|
| 33 | 0.13% |
| 34 | 0.19% |
| 42 | 0.06% |
| 37 | 0.02% |
| 29 | 0.15% |
| 26 | 1.63% |
| 20 | 1.00% |
| 28 | 0.06% |
| 11 | 0.62% |
| 27 | 0.51% |
| 22 | **3.72%** |
| 17 | **18.81%** |
| 19 | 3.69% |
| 21 | 0.51% |
| 1 | **2.42%** |
| 43 | **3.21%** |
| 31 | **0.59%** |
| 24 | **3.68%** |
| 16 | **4.16%** |
| 9 | 0.36% |
| 35 | 1.57% |
| 6 | **8.68%** |
| 30 | **2.46%** |
| 14 | 0.21% |
| 32 | 0.32% |
| 12 | **2.55%** |
| 23 | **7.79%** |
| 10 | **8.22%** |
| 40 | 1.40% |
| 38 | 0.08% |
| 36 | 1.15% |
| 39 | 1.42% |
| 15 | **12.70%** |
| 7 | 0.42% |
| 18 | 0.04% |
| 8 | 0.19% |
| 25 | 0.49% |
| 13 | 0.55% |
| 41 | 0.02% |
| 4 | **1.42%** |
| 5 | 0.13% |
| 2 | 2.46% |
| 3 | 0.23% |

Supplemental Single Cell Patient Information

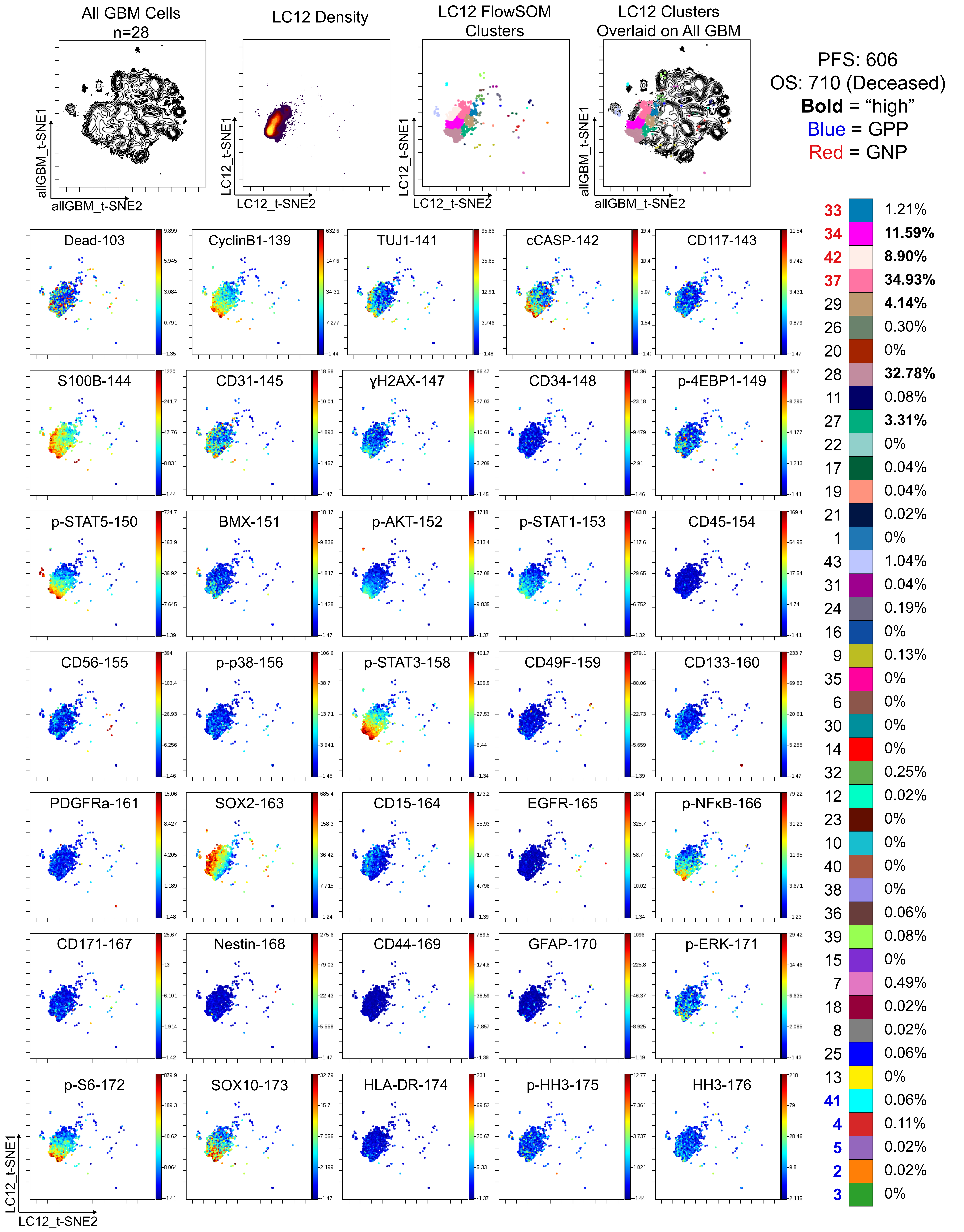

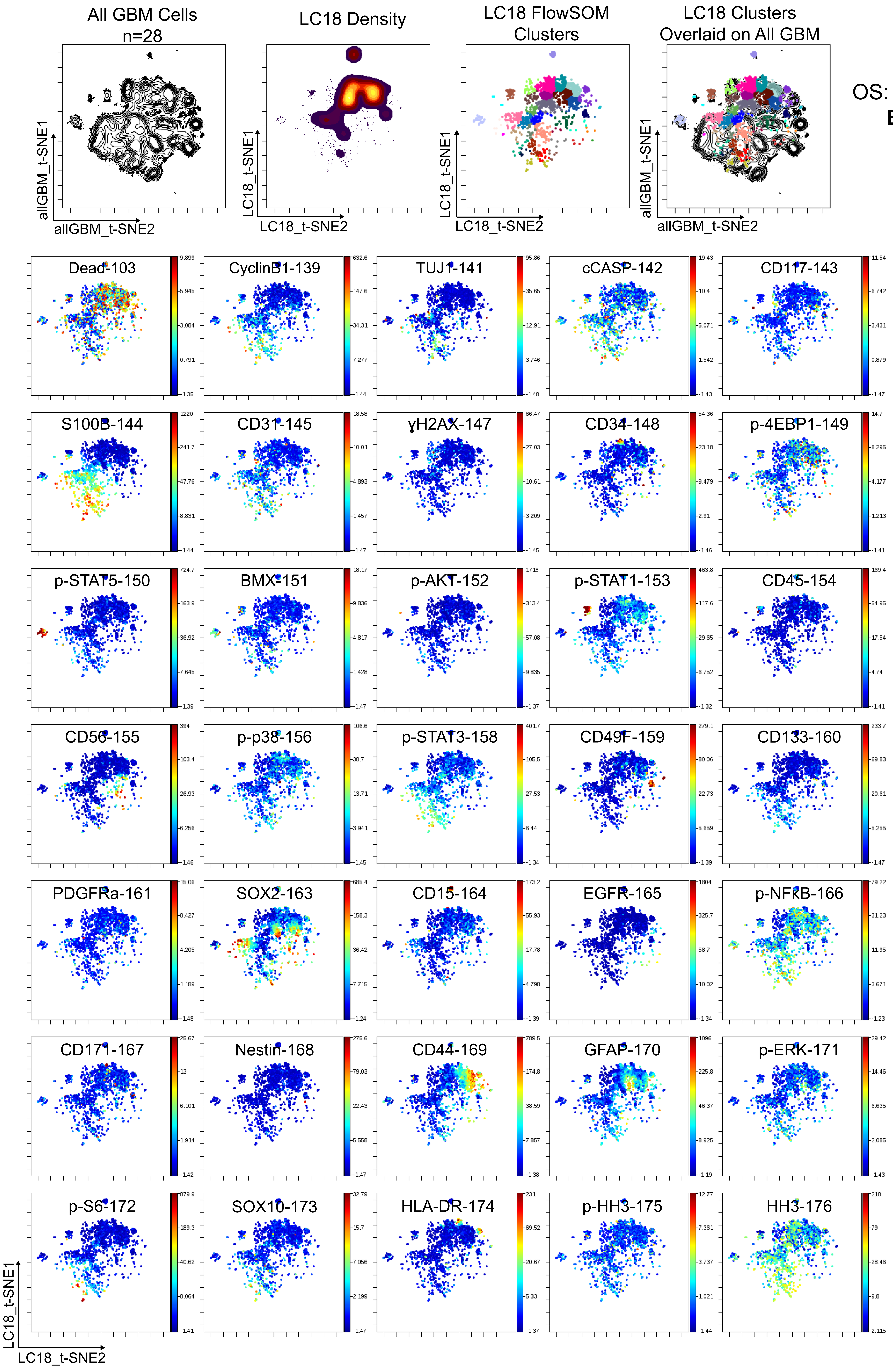


PFS: 133
OS: 240 (Deceased)
**Bold** = “high”
Blue = GPP
Red = GNP

| | |
|---|---|
| **33** | **3.10%** |
| **34** | **2.76%** |
| **42** | 0.36% |
| **37** | 0.04% |
| 29 | 0.25% |
| 26 | 0.51% |
| 20 | **1.25%** |
| 28 | 0% |
| 11 | **2.53%** |
| 27 | 0.21% |
| 22 | **13.18%** |
| 17 | 1.25% |
| 19 | 3.59% |
| 21 | 0.21% |
| 1 | 0.02% |
| 43 | 0.66% |
| 31 | **8.77%** |
| 24 | **12.82%** |
| 16 | **7.26%** |
| 9 | 0.47% |
| 35 | **9.94%** |
| 6 | 0.15% |
| 30 | **7.60%** |
| 14 | 0.83% |
| 32 | **1.85%** |
| 12 | 0.15% |
| 23 | **5.71%** |
| 10 | 0.13% |
| 40 | 1.10% |
| 38 | **5.07%** |
| 36 | 0.93% |
| 39 | 0.66% |
| 15 | **3.99%** |
| 7 | 0% |
| 18 | 0.08% |
| 8 | 0.11% |
| 25 | 1.91% |
| 13 | 0% |
| **41** | 0.28% |
| **4** | 0.17% |
| **5** | 0% |
| **2** | 0.06% |
| **3** | 0.02% |

Supplemental Single Cell Patient Information

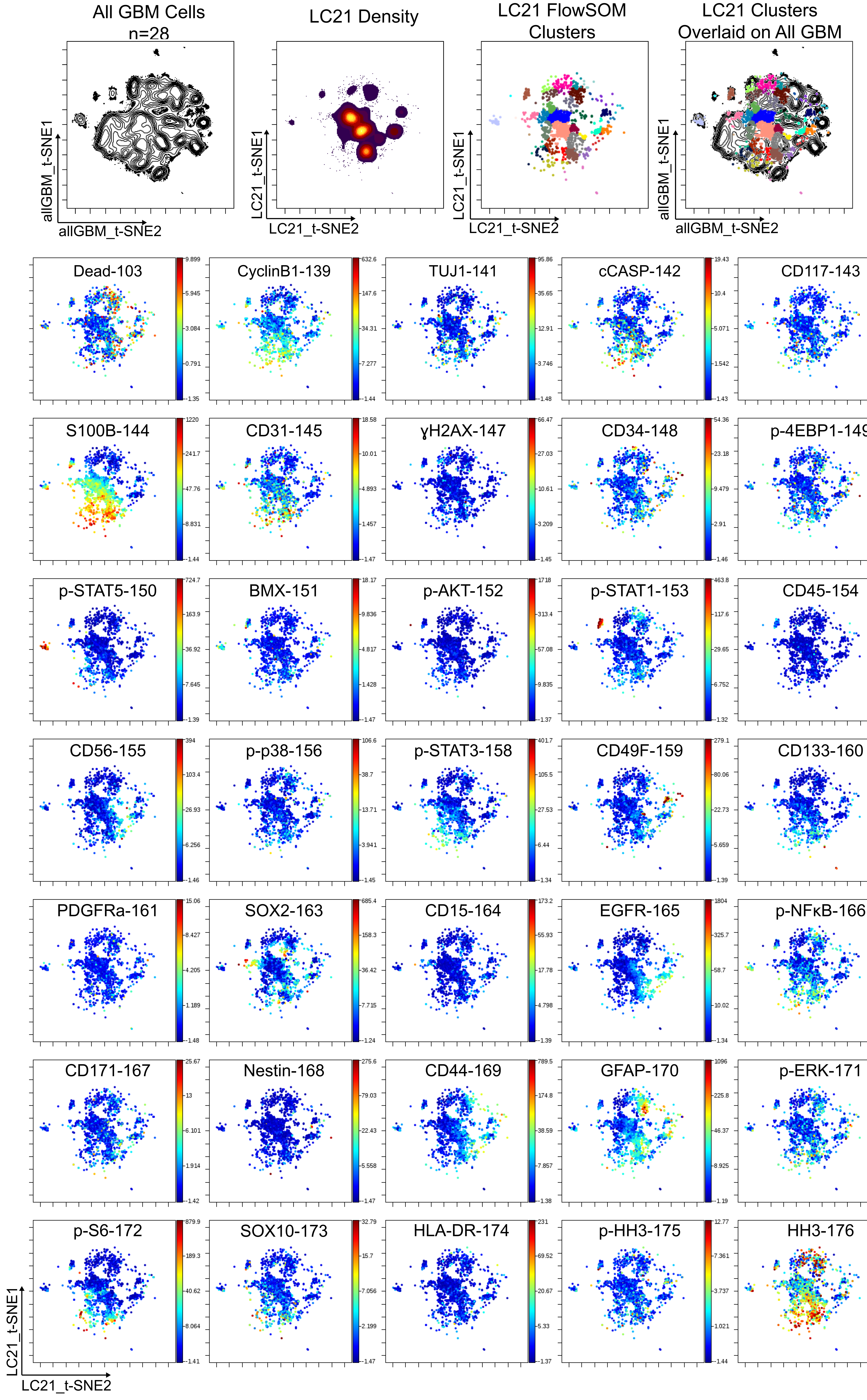


PFS: 267
OS: 267 (Deceased)
**Bold** = "high"
Blue = GPP
Red = GNP

| | |
|---|---|
| 33 | 1.83% |
| 34 | 0.89% |
| 42 | 0.02% |
| 37 | 0% |
| 29 | 0.06% |
| 26 | **1.97%** |
| 20 | 1.10% |
| 28 | 0% |
| 11 | **1.83%** |
| 27 | 0.11% |
| 22 | 0.11% |
| 17 | 0.87% |
| 19 | **21.76%** |
| 21 | 0.62% |
| 1 | 0.11% |
| 43 | 0.89% |
| 31 | 0.19% |
| 24 | 1.27% |
| 16 | 0.06% |
| 9 | 0.68% |
| 35 | 1.34% |
| 6 | **14.06%** |
| 30 | 0.47% |
| 14 | 0.45% |
| 32 | **2.68%** |
| 12 | **3.93%** |
| 23 | 2.48% |
| 10 | 0.17% |
| 40 | 1.00% |
| 38 | 0% |
| 36 | 0.98% |
| 39 | 0.36% |
| 15 | 0.23% |
| 7 | 0.13% |
| 18 | **8.11%** |
| 8 | **3.72%** |
| 25 | **21.80%** |
| 13 | **1.17%** |
| 41 | 0.02% |
| 4 | 0.72% |
| 5 | **0.57%** |
| 2 | 1.25% |
| 3 | 0% |

Supplemental Single Cell Patient Information

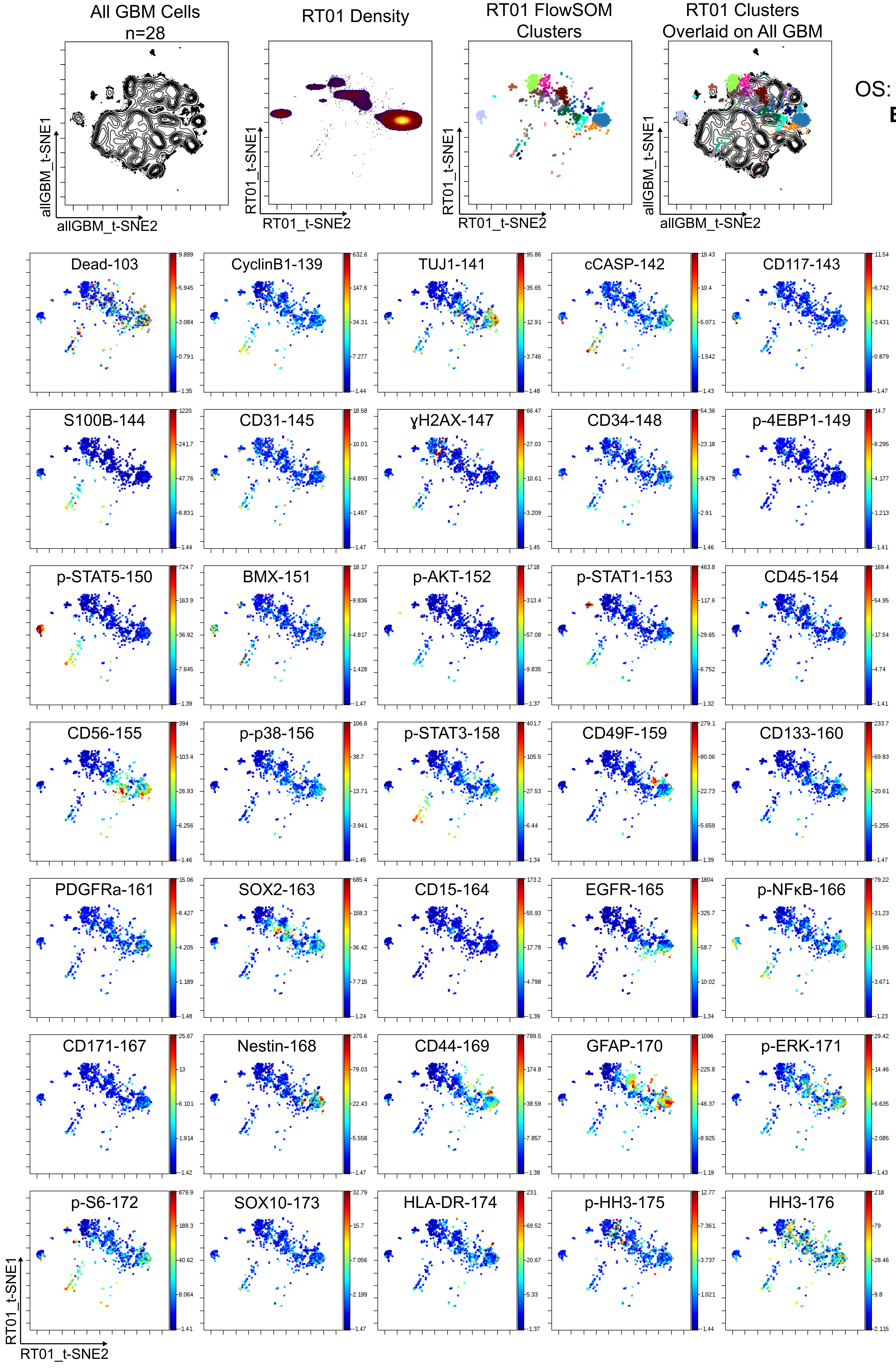


PFS: 162
OS: 198 (Deceased)
**Bold** = "high"
Blue = GPP
Red = GNP

| | |
|---|---|
| 33 | 0.02% |
| 34 | 0% |
| 42 | 0% |
| 37 | 0% |
| 29 | 0.08% |
| 26 | 0.21% |
| 20 | 0% |
| 28 | 0.21% |
| 11 | **1.23%** |
| 27 | 0.55% |
| 22 | 0.08% |
| 17 | 3.12% |
| 19 | 0.04% |
| 21 | 0.21% |
| 1 | **63.52%** |
| 43 | **4.76%** |
| 31 | **0.53%** |
| 24 | **4.01%** |
| 16 | 1.10% |
| 9 | 0% |
| 35 | 1.34% |
| 6 | 0.08% |
| 30 | 0.08% |
| 14 | 0% |
| 32 | 0.15% |
| 12 | **1.74%** |
| 23 | **4.84%** |
| 10 | **1.04%** |
| 40 | **2.72%** |
| 38 | 0% |
| 36 | **3.14%** |
| 39 | **3.72%** |
| 15 | 0.17% |
| 7 | 0.11% |
| 18 | 0.02% |
| 8 | 0% |
| 25 | 0.04% |
| 13 | 0.04% |
| 41 | 0.08% |
| 4 | 0.11% |
| 5 | 0.02% |
| 2 | 0.83% |
| 3 | 0.02% |

Supplemental Single Cell Patient Information

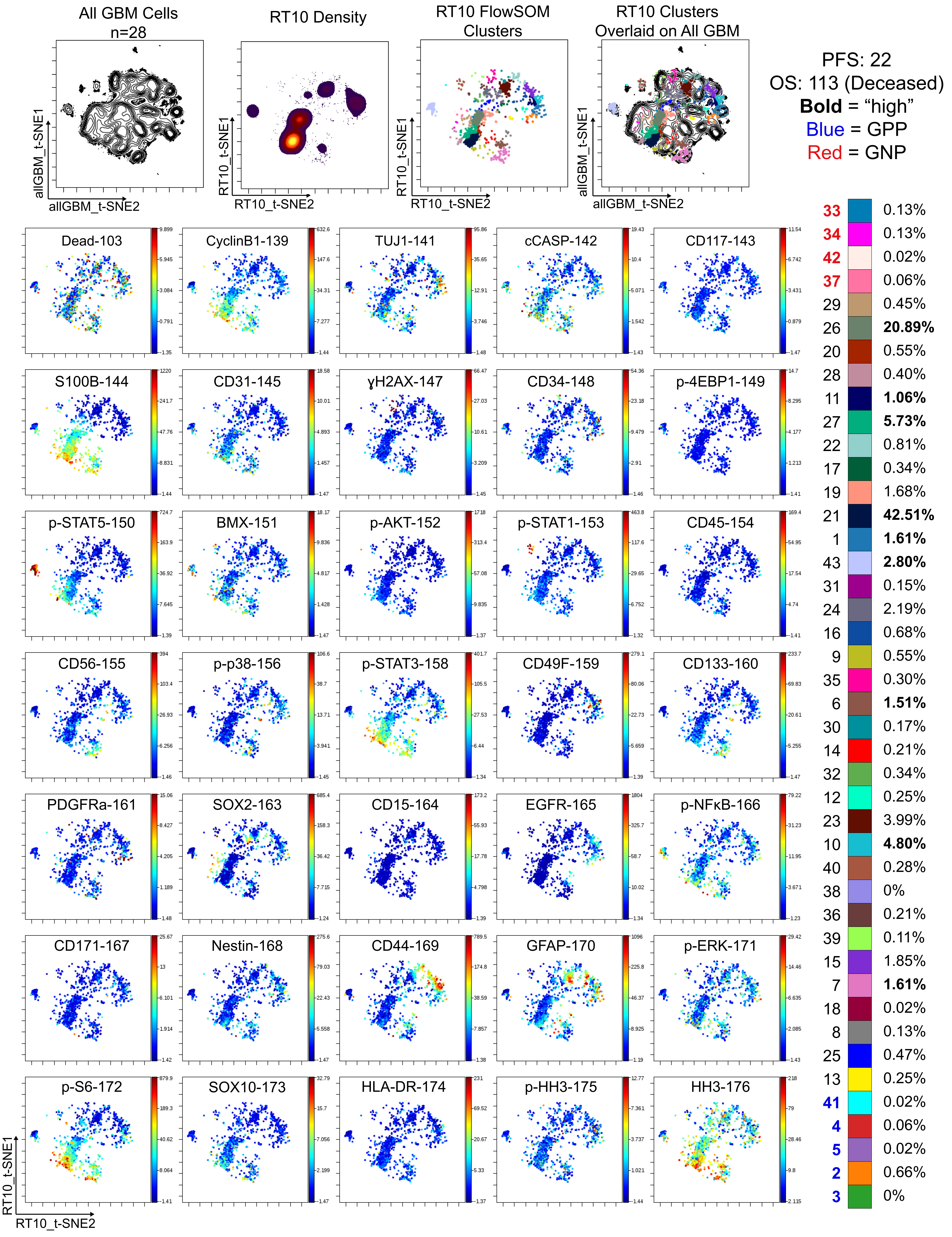

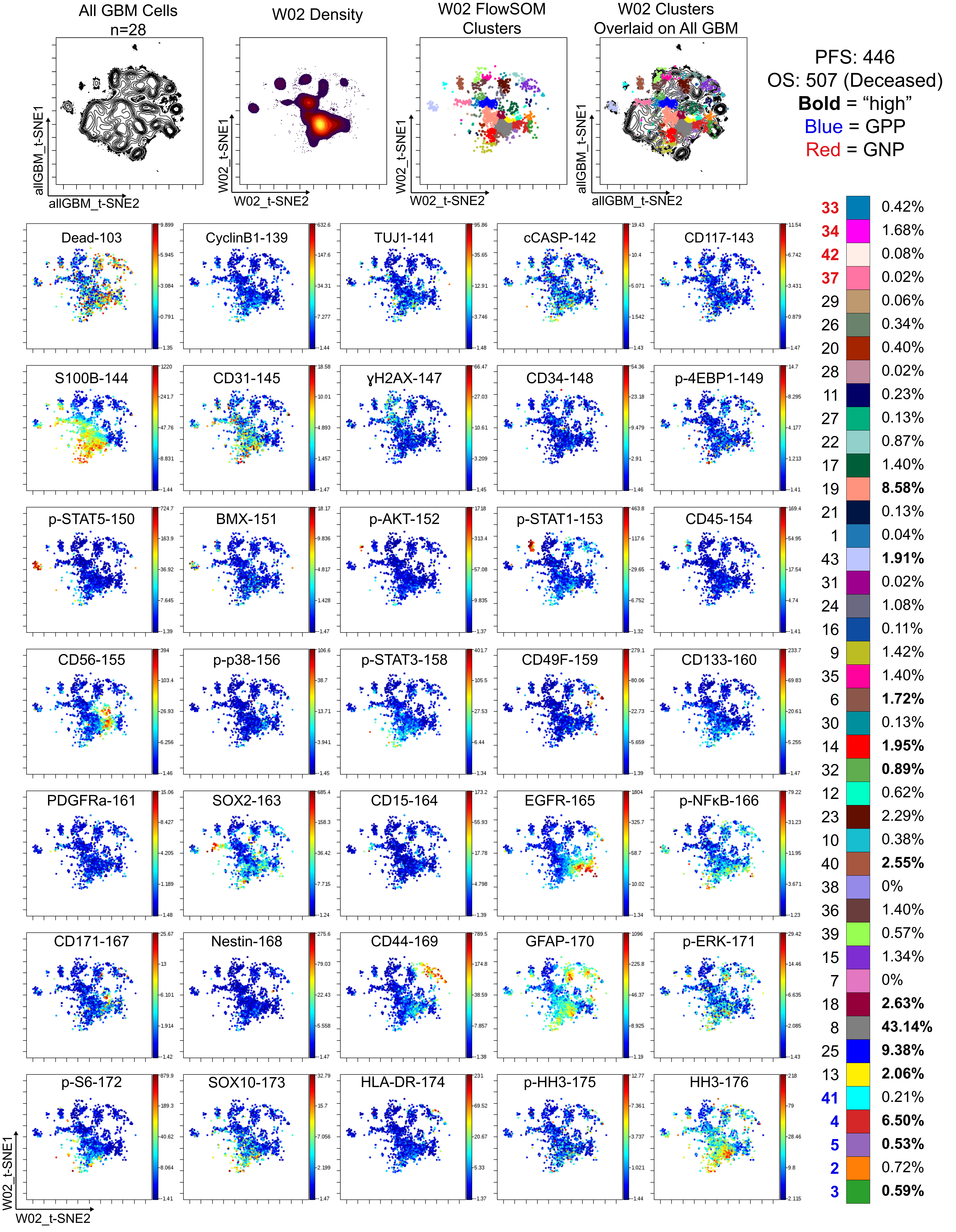

All GBM Cells n=28
W02 Density
W02 FlowSOM Clusters
W02 Clusters Overlaid on All GBM
allGBM_t-SNE1
allGBM_t-SNE2
W02_t-SNE1
W02_t-SNE2
PFS: 446
OS: 507 (Deceased)
Bold = "high"
Blue = GPP
Red = GNP
Dead-103
CyclinB1-139
TUJ1-141
cCASP-142
CD117-143
S100B-144
CD31-145
γH2AX-147
CD34-148
p-4EBP1-149
p-STAT5-150
BMX-151
p-AKT-152
p-STAT1-153
CD45-154
CD56-155
p-p38-156
p-STAT3-158
CD49F-159
CD133-160
PDGFRa-161
SOX2-163
CD15-164
EGFR-165
p-NFκB-166
CD171-167
Nestin-168
CD44-169
GFAP-170
p-ERK-171
p-S6-172
SOX10-173
HLA-DR-174
p-HH3-175
HH3-176
33 0.42%
34 1.68%
42 0.08%
37 0.02%
29 0.06%
26 0.34%
20 0.40%
28 0.02%
11 0.23%
27 0.13%
22 0.87%
17 1.40%
19 8.58%
21 0.13%
1 0.04%
43 1.91%
31 0.02%
24 1.08%
16 0.11%
9 1.42%
35 1.40%
6 1.72%
30 0.13%
14 1.95%
32 0.89%
12 0.62%
23 2.29%
10 0.38%
40 2.55%
38 0%
36 1.40%
39 0.57%
15 1.34%
7 0%
18 2.63%
8 43.14%
25 9.38%
13 2.06%
41 0.21%
4 6.50%
5 0.53%
2 0.72%
3 0.59%

Supplemental Single Cell Patient Information

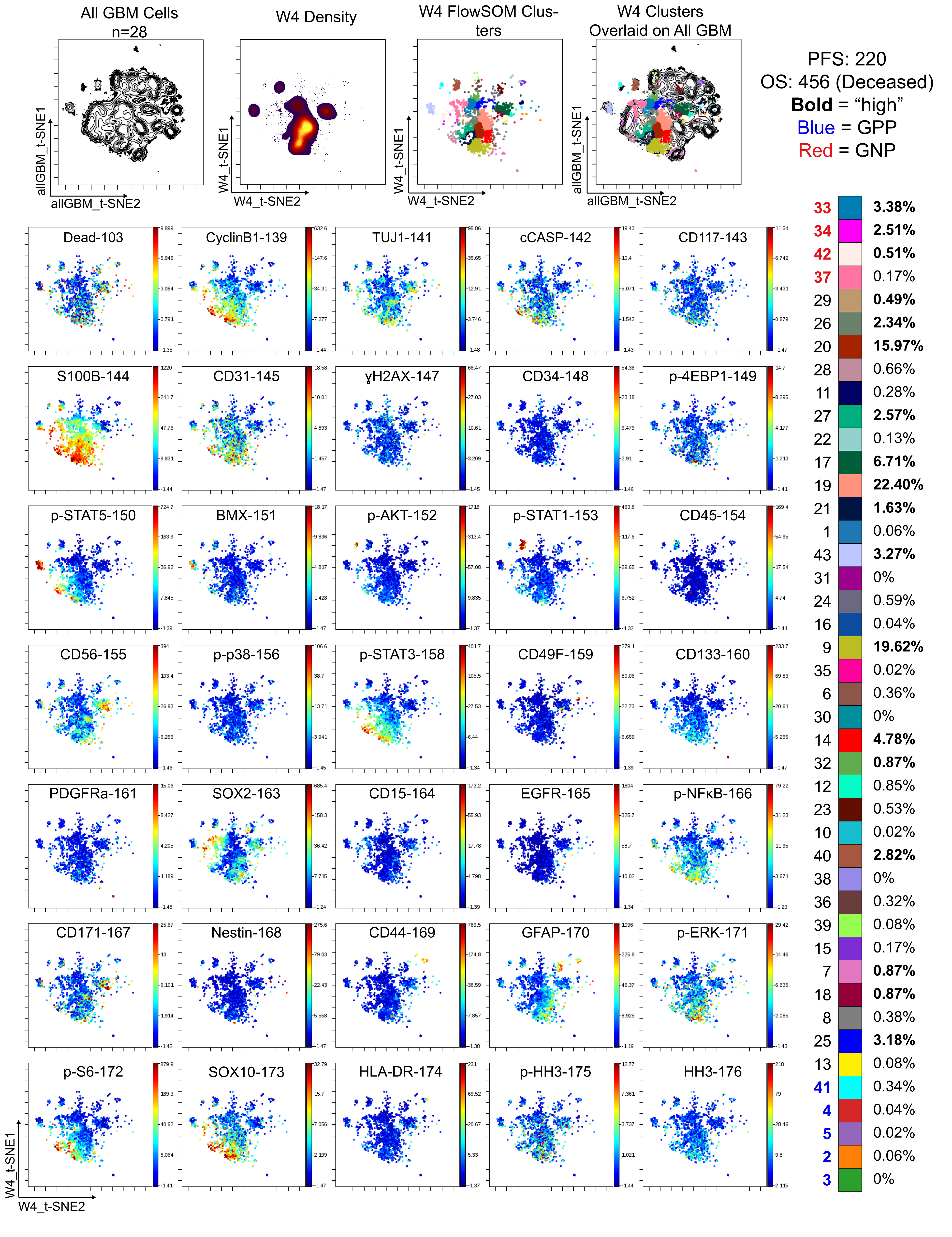

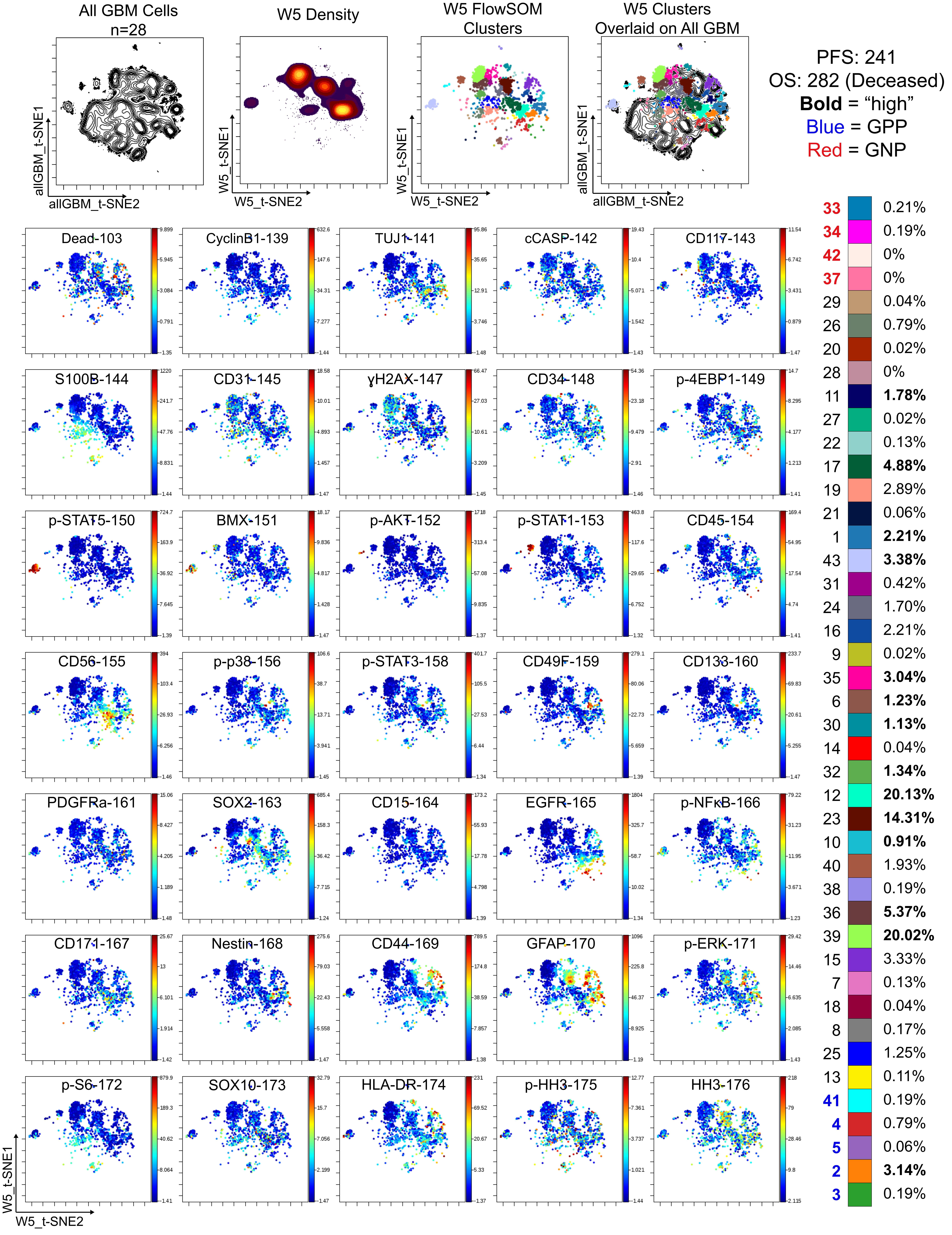
All GBM Cells
n=28
allGBM_t-SNE1
allGBM_t-SNE2
W5 Density
W5_t-SNE1
W5_t-SNE2
W5 FlowSOM Clusters
W5_t-SNE1
W5_t-SNE2
W5 Clusters Overlaid on All GBM
allGBM_t-SNE1
allGBM_t-SNE2
PFS: 241
OS: 282 (Deceased)
Bold = "high"
Blue = GPP
Red = GNP
33 0.21%
34 0.19%
42 0%
37 0%
29 0.04%
26 0.79%
20 0.02%
28 0%
11 1.78%
27 0.02%
22 0.13%
17 4.88%
19 2.89%
21 0.06%
1 2.21%
43 3.38%
31 0.42%
24 1.70%
16 2.21%
9 0.02%
35 3.04%
6 1.23%
30 1.13%
14 0.04%
32 1.34%
12 20.13%
23 14.31%
10 0.91%
40 1.93%
38 0.19%
36 5.37%
39 20.02%
15 3.33%
7 0.13%
18 0.04%
8 0.17%
25 1.25%
13 0.11%
41 0.19%
4 0.79%
5 0.06%
2 3.14%
3 0.19%
Dead-103
CyclinB1-139
TUJ1-141
cCASP-142
CD117-143
S100B-144
CD31-145
γH2AX-147
CD34-148
p-4EBP1-149
p-STAT5-150
BMX-151
p-AKT-152
p-STAT1-153
CD45-154
CD56-155
p-p38-156
p-STAT3-158
CD49F-159
CD133-160
PDGFRa-161
SOX2-163
CD15-164
EGFR-165
p-NFκB-166
CD171-167
Nestin-168
CD44-169
GFAP-170
p-ERK-171
p-S6-172
SOX10-173
HLA-DR-174
p-HH3-175
HH3-176
W5_t-SNE1
W5_t-SNE2

Supplemental Single Cell Patient Information

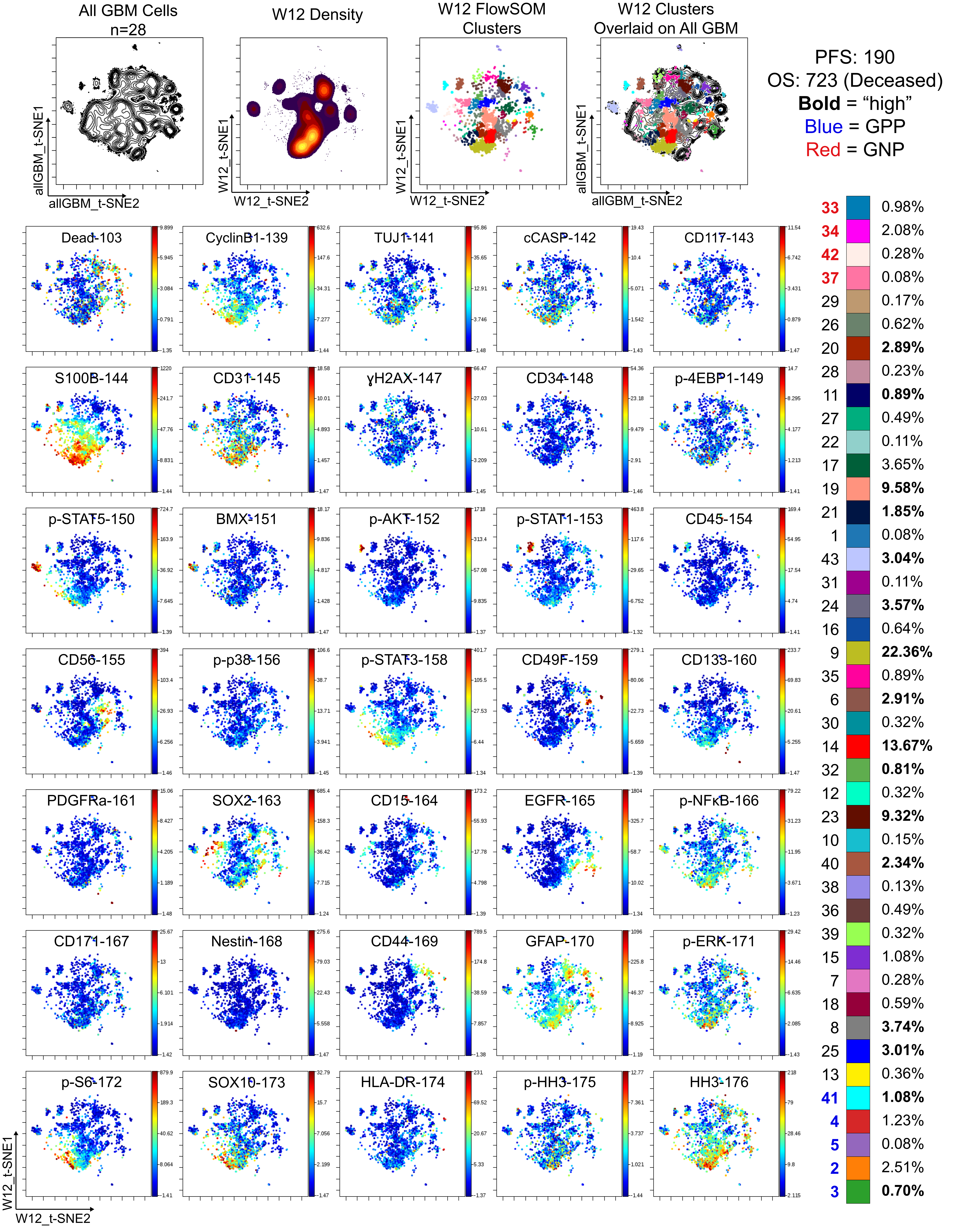
